# The Algorithm for Reversible Jump Inference of Motifs

**DOI:** 10.64898/2025.12.25.696506

**Authors:** Ali M. Farhat, Lydia Freddolino

## Abstract

Understanding sequence specificity in transcription factor binding is a critical step in understanding gene regulation. DNA-protein binding experiments such as ChIP-chip and ChIP-seq can reveal where on the genome a particular transcription factor of interest is bound, and similar experiments such as ATAC-seq or DNase hypersensitivity can reveal the profile of genome-wide protein binding and chromatin accessibility at the time of an experiment. However, demultiplexing the binding of multiple transcription factors represented in a single occupancy trace can be difficult. While many tools exist for extracting binding motifs from single transcription factor assays, comparatively few exist for deconvolving binding motifs from many transcription factor experiments, and they often require additional information on the number and/or nature of factors to be considered. Here, we developed The Algorithm for Reversible Jump Inference of Motifs (TARJIM), which translates DNA-protein binding data by inferring a set of sequence motifs that explain the data in question. By using a reversible jump Metropolis Hastings algorithm, TARJIM is able to infer both the number of motifs present and their sequence identity by using Bayesian techniques on the binding model itself. Using TARJIM, we have shown that not only can we deduce sequence motifs for known transcription factors, we are also able to extract sequence motifs from a mixture of sequence binding data, allowing us to extract information even from protein-DNA binding experiments where we do not know the number of transcription factors represented.

## 2 Introduction

Understanding how genes are regulated is an absolutely essential line of inquiry in modern genomics. Central to this line of inquiry in both bacteria [10, 29] and in eukaryotes [16, 37] is the analysis of transcription factors (TFs), proteins which bind DNA and regulate gene transcription. We can examine the locations of binding of TFs using Chromatin ImmunoPrecipitation (ChIP) experiments, whether on microarrays (ChIP-chip) [47] or with the use of sequencing libraries (ChIP-seq) [6, 26]. ChIP experiments are extremely powerful and can show precisely where a TF of interest binds under some specific physiological condition. There are many tools to take binding data from ChIP experiments and use them to predict sequence specific binding motifs, allowing us to understand how TFs encode their binding locations on the genome [5, 30]. Furthermore, the experimental toolbox is not limited by needing to identify a factor of interest *a priori*: there are now protein agnostic analyses of DNA bound by TFs, such DNAse-seq [9], ATAC-seq [11], and IPOD(-HR) [19, 46]. Protein agonistic binding experiments are extremely powerful, as they not only encode the binding on the genome that we know should exist, but also the binding of all the TFs we do not yet know about. If it were possible to deconvolve all of the binding modes represented in protein agnostic experiments, it would be possible to infer information about all TF binding the genome at the time of the experiment. This would make protein agnostic binding experiments powerful tools for discovery of both novel TFs and novel binding modes of existing TFs. However, there are very few TF binding signal deconvolution algorithms. For example, DIVERSITY [33] is a tool optimized for inferring multiple binding motifs of single transcription factors. It requires a list of DNA fragments containing peaks, and performs multiple inferences on different numbers of binding modes before performing an *a posteriori* comparison of the results from each. CENTIPEDE [38] takes in a set of known motifs and DNAse-seq data, predicting binding profiles. CENTIPEDE can be used for some motif inference when there are many known motifs accounted for, but this requires a great deal of *a priori* knowledge of transcription factor binding modes to expect. MEME-ChIP [30] requires a set of sequences from pre identified peaks and benefits greatly from an equally sized set of sequences specifically without binding (which we will call “null sequences”) and from all the sequences supplied being of equal width. ChromeBPNet [36] has a focus on recapitulating a model of chromatin accessibility to recapitulate a binding profile in eukaryotic genomes. In ChromeBPNet, transcription factor motifs are a secondary concern which are extracted by feeding ChromeBPNet’s predicted regions of occupancy into a clustering algorithm, which does not itself account for avoiding matching null sequences in its predicted motifs.

While each demultiplexing tool mentioned above provides its own unique and important advantages, each of them also require a great deal of prior information (CENTIPEDE) [38], fail to leverage information granted from null sequences (DIVERSITY) [33], or only leverage information from null sequences in equal proportion to the binding sequences (MEME-ChIP, ChromeBPNet) [30, 36] despite genomes on average having more unbound regions than bound regions in experiments (see the coverage of identified peaks in [14, 19, 28, 46]). These algorithms also primarily report motifs based on base pair frequency in peak regions. This is not fundamentally problematic; however, DNA-protein binding occupancy traces encode a great deal about the biophysics of transcription factor binding [48], and peak finders must ultimately set an arbitrary threshold when determining whether or not a region is a peak. We assert that it should be a higher priority for a motif to match a highly peaked region than a mildly peaked region, and that is more important for a motif to avoid matching a region that is purely experimental noise than a region that holds a feature that is possibly a peak but below an arbitrary cutoff; following our argument, an approach that directly matches a binding profile to the quantitative, experimentally observed binding data would be ideal. These problems are more easily mitigated when looking for single transcription factor binding behavior, but demultiplexing binding from multiple transcription factors is more complicated. This complexity is compounded by the fact that estimating the number of motifs to demultiplex from a trace is non-trivial. DIVERSITY attempts to solve this by performing separate inferences for every count of motifs up to some user defined maximum, then comparing them *a posteriori*. CENTIPEDE and ChromeBPNet each use clustering algorithms to attempt to compress the number of motifs they infer, but do not further reference null sequences, so the representative motif of a large enough cluster could match null sequences more strongly than they should. MEME-ChIP runs multiple algorithms to try to capture motifs that each reference the null sequences, but these algorithms do not have redundancy pruning when they find have found potentially overlapping motifs.

To address the need for a tool specifically designed for precision motif demultiplexing from protein agnostic binding experiments, we have created The Algorithm for Reversible Jump Inference of Motifs (TARJIM). TARJIM seeks to address the problems mentioned above by leveraging several different theoretical approaches. First, TARJIM treats transcription factor sequence motifs as describing binding energy optima and the surrounding energy landscape, using the language of DNA sequences. In particular, a single TARJIM motif consists of one “best” sequence with a list of penalties coming from deviating from that sequence. If a sequence deviates too far from the binding energy optimum, then it is assumed that binding at that sequence cannot be described by that motif, and is dominated either by nonspecific interactions or by a different energy optimum, which would be represented as a different motif. A transcription factor need not have a one-to-one relationship with this definition of a sequence motif. For example, if a transcription factor binds DNA mainly by shape [13, 42], the sequence motifs TARJIM might infer from a ChIP experiment of that transcription factor might represent that as multiple sequence optima with the same shape but different sequences. If there are strong position dependent effects—e.g., suppose a motif position is better as an A if its neighbor is a C, but better as a G if that neighbor is an A [27]—these sequence motifs could split into multiple motifs to model this—such as the prior example by having one motif which heavily favors AC and another which heavily favors GA. However, simply treating motifs as energy optima presents a particular challenge: the more granularly we model the energy landscape by increasing the number of motifs, the better we can fit the experiment. But the fewer motifs TARJIM infers, the more understandable the binding behavior of these motifs becomes, and the less overfitting in our inference. To mitigate the problem of determining motif count, TARJIM is a Reversible Jump Metropolis Hastings algorithm [22], which allows us to infer the number and lengths of the motifs as part of the inference itself. We accelerated this inference with parallel tempering, which we prove below to be compatible to Reversible Jump Metropolis Hastings. To further avoid overfitting and account for experiments being fundamentally noisy, TARJIM’s likelihood function is based not on product of independent location likelihoods, but on fitting a whole data set background noise distribution while minimizing matching to null sequences. To quantify the distance between the background distribution and residual distribution induced by a motif set, we use a modification of the Anderson-Darling statistic which was first derived by Ahmad, Sinclair, and Spurr and most often used in flood prediction models [1, 43], as it is particularly sensitive to likely peaks while being less sensitive to likely noise. As this is a whole distribution parameter and predicting whole occupancy traces is computationally intense, TARJIM computes the residuals it requires precisely for sites with binding and approximately for null sequences. We will first show the correctness of TARJIM in inferring binding motifs from relatively simple data: a TrpR ChIP-chip and an ArgR ChIP-chip in E. coli [14]. We will then demonstrate its ability to deconvolve binding modes by running TARJIM on an artificial mix of both of these ChIP-chip data sets, emulating the data that could provide protein-agnostic occupancy across the genome. We further follow up by demonstrating its utility in capturing variable binding behavior from global regulator binding data by using TARJIM to analyze Lrp ChIP-seq experiments that demonstrated changes in Lrp binding specificity across changing physiological conditions [28]. And finally, we will demonstrate how TARJIM can deconvolve the binding of many transcription factors across the whole genome when it is fed data from a protein agnostic DNA-protein binding experiment such as IPOD-HR, using the data from the original IPOD-HR experiment in *E. coli* [19].

## 3 Results

TARJIM’s fundamental theoretical basis conceives of DNA-protein occupancy traces as a composition of two phenomena: a signal based on sequence motifs which provoke protein binding, and a noise trace which is added onto the signal. Sequence motifs are not necessarily one-to-one with transcription factors: one factor can potentially have multiple binding modes [28, 51] or bind DNA in a complex way only partially related to sequence, which might require multiple sequence motifs to account for [13, 27, 42]. Conversely, a multi-protein complex binding a single consistent sequence would only produce one motif. To account for this complication, TARJIM performs inference on the number of motifs itself using a Reversible Jump Metropolis Hastings algorithm [22]. This algorithm allows us to account for the possibility of uncertainty in the number of binding motifs while minimizing overfitting using a restrictive prior on the motif count itself, allowing TARJIM to naturally infer a proper model size.

To predict binding from a set of motifs, TARJIM assumes that every sequence in the genome has the opportunity to show binding signal from each of the motifs. In general, ChIP data sets tend to have peaks that are roughly Gaussian shaped, with relatively consistent widths based on the experiment: see Figure 1A for example ChIP-chip data from [14]. To calculate an occupancy trace, TARJIM assigns each motif a maximum peak height, and assumes that a binding peak will have that peak height only if the underlying sequence perfectly matches the motif. TARJIM assigns smaller peaks to secondary matches, consistent with predicted free energy penalties, similar to the characterization of binding in [48]. To account for all motifs in the set, the occupancy traces of each individual motif are added together. The assumed additivity of occupancy traces comes with a fundamental assumption of independence of binding phenomena: this may not be strictly true, but we are assuming that it is generally unlikely for multiple binding modes to strongly compete for the same sequence. To calculate the likelihood of a particular motif set, we calculate the upper tail version of the modified Anderson Darling statistic first derived by Ahmad, Sinclair, and Spurr [1, 43] (which we label ð). For the sake of convenience, we propose calling ð the “Ahmad Statistic”, and its low tail counterpart the “Sinclair statistic.” The Ahmad statistic should be particularly sensitive to missed or mispredicted peaks and has the following form:

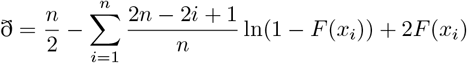

**Figure 1.**
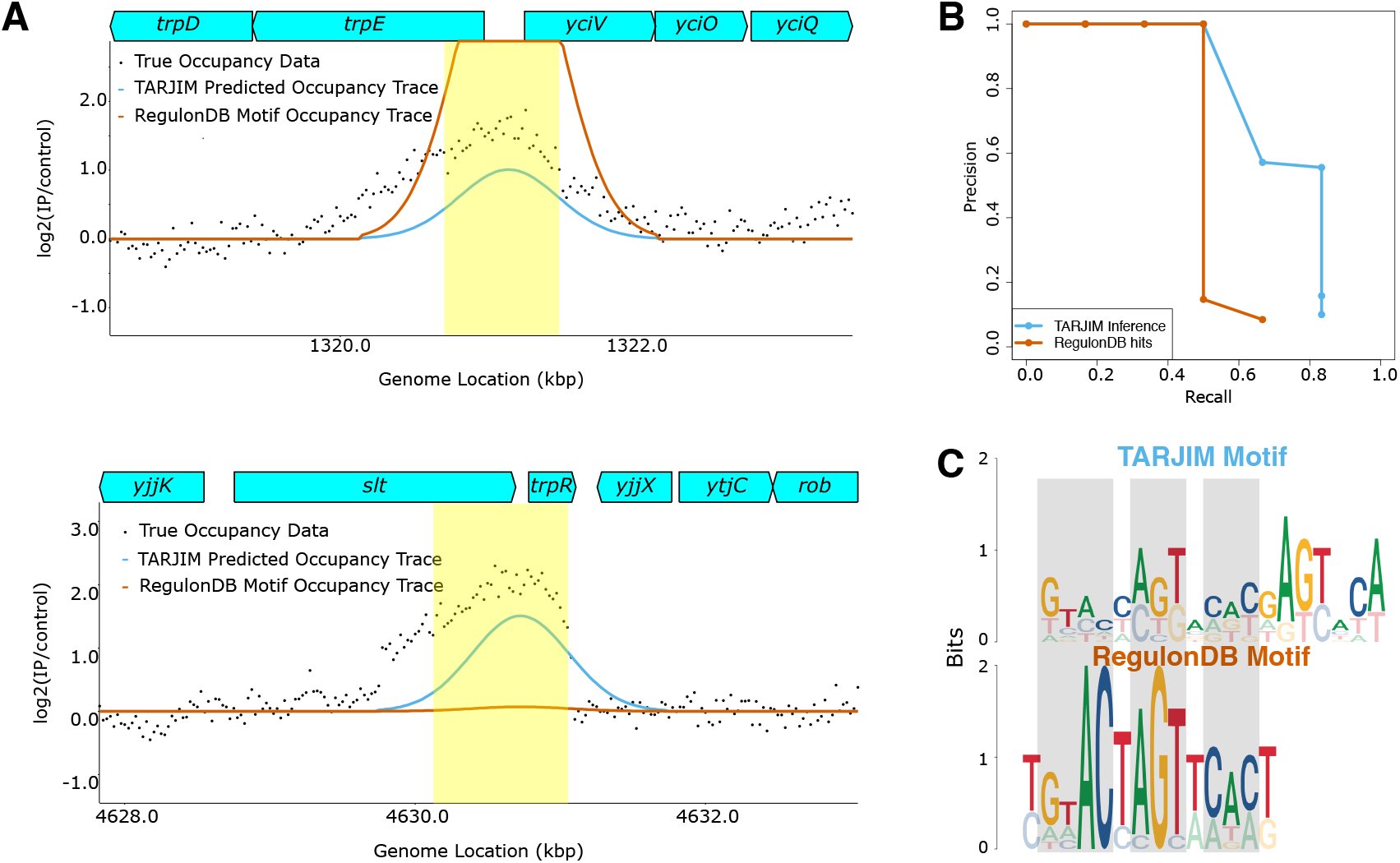
TARJIM’s analysis of TrpR binding data.**A** The occupancy of TrpR at the *trp* operon (top) and near the *trpR* locus itself (bottom), with the [14] data as black dots, macs3’s peak calls shaded in yellow, TARJIM’s predicted occupancy trace in blue, and the [40] occupancy in red. Note that loci in occupancy traces were only shown if they were larger than 5% of the plotting window: e.g., the *trpL* locus was omitted from the top plot. **B** The precision-recall curves for the TrpR peaks from the [40] PWM (black) and TARJIM’s predictions (blue). **C** TARJIM’s predicted TrpR motif (top) as compared to the RegulonDB [40] motif for TrpR (bottom). The grey shading shows where the TARJIM motif’s primary sequence is the same as that of the RegulonDB motif.

Where *x*_*i*_ is the *i*th smallest absolute value of the residual accounted for in the motif trace, *n* is the number of points accounted for in the motif trace, and *F*(*x*_*i*_) is the CDF of the absolute values of the noise, which we fit to a half normal distribution before the inference.

We use the probability distribution of the Ahmad statistic as our likelihood function. More details can be found in the methodology section.

### 3.1 TrpR Binding

In order to demonstrate the application of TARJIM to sequence-specific local regulators in bacteria, we applied TARJIM to a set of ChIP-chip data for the amino acid-responsive local regulators TrpR and ArgR [14]. TrpR is a well characterized transcription factor in *E. coli* which regulates tryptophan metabolism. Its binding is extremely localized: using macs3 [49], we selected six primary binding locations from the ChIP-chip data in [14], two of which are shown in Figure 1A. Since Jacob and Monod [25], TrpR has been studied as a paradigmatic example of a gene regulatory protein in *E. coli*, and it has a well characterized binding motif, published in RegulonDB [40], and is thus available for comparison with our inferred motifs.

Using TARJIM, we derived a binding motif set for TrpR from the data from [14]. The maximum *a posteriori* prediction from TARJIM contains one motif, pictured in Figure 1. The motif clearly corresponds to the RegulonDB motif [40]. However, it is different from the [40] motif in key ways: it deviates from main 6-mer, preferring an ACCAGT over an ACTAGT in that location, and it also contains additional specification on the right hand side of the motif. To compare our predicted binding, we used macs3 [49] to call gold standard peaks for the TrpR data shown here, as well as both the ArgR and mixed data we will show later. We then used these gold standard peaks to generate precision-recall curves: we generated binding locations with FIMO [21], and considered a location a hit if it was inside the peak location generated by macs3. Further details are elaborated on in the Methods section. The area under the precision-recall curve (AUPR) for the RegulonDB [40] motif’s peak prediction locations is 0.52, while the TARJIM inference has an AUPR of 0.72.

### 3.2 Disentangling ArgR’s Binding Modes

ArgR is another transcription factor in *E. coli*. It regulates multiple facets of arginine metabolism, along with histidine transport, and has even been implicated in plasmid stability [12, 44]. It is more promiscuous than TrpR, binding roughly ten times as many locations [14, 40]. When using TARJIM with the ArgR ChIP-chip from [14], TARJIM infers many more motifs than in the case of TrpR: 8 total motifs. We can also use the likelihood function of TARJIM to our advantage, and report motif sets in order of which motifs most improve the likelihood function. To understand TARJIM’s inference, we analyzed both its full inferred ArgR motif set and the set from taking only the top binding motif.

The motif set for ArgR recapitulates important aspects of binding from the occupancy trace more accurately than the RegulonDB motif (Figure 2A), despite the fact that the top TARJIM ArgR motif resembles the RegulonDB ArgR motif (Figure 2B) [40]. Comparing the precision-recall curves for the ArgR peaks in Figure 2C for the RegulonDB [40] motif against the motif sets from the full TARJIM inferences reveals an important idea: the RegulonDB motif [40] matches binding peaks because it matches many parts of the genome in general. After it matches its top few locations, its precision drops to about two-thirds. The failure mode for matching binding locations is significantly different for the motif sets inferred by TARJIM: TARJIM’s precision hovers near unity as it identifies more binding locations until it hits a ceiling, dropping the precision to zero with no improvement in recall. This is a function of an important tendency of TARJIM: it strongly selects against motif sets with significant off target binding, and will avoid proposing such motif sets even at the cost of failing to match binding sites. To show how our inference improves with more included motifs, we have plotted the AUPRs when we take different sized subsets from the full TARJIM motif set. With one motif, TARJIM’s AUPR is already is comparable to the RegulonDB motif despite matching fewer peaks, and with even just the top two motifs, the TARJIM AUPR is strictly superior (Figure 2D).

**Figure 2.**
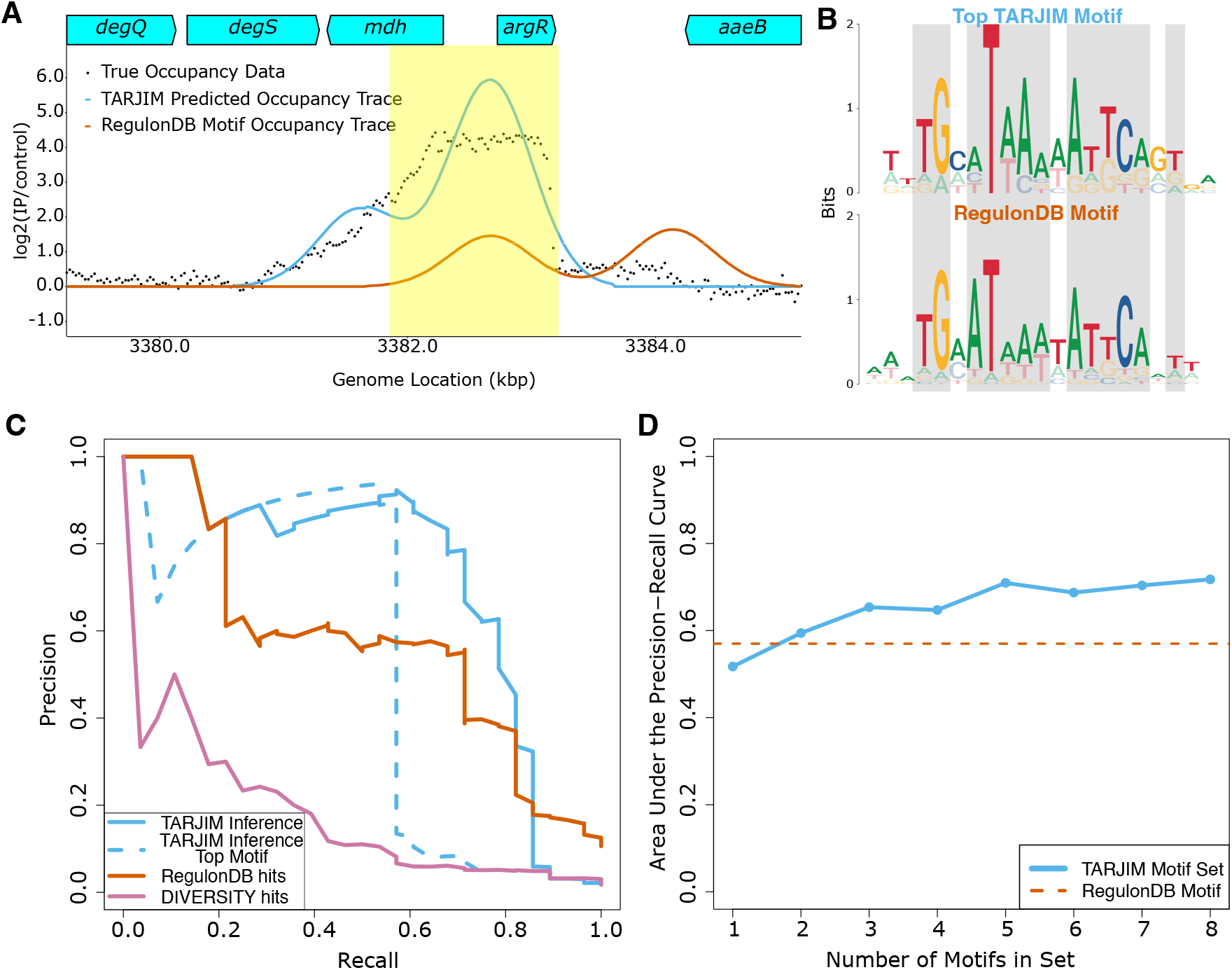
TARJIM’s analysis of ArgR binding data. **A** The occupancy of ArgR at the *argR* locus, with the [14] data as black dots, macs3’s peak call shaded in yellow, TARJIM’s predicted occupancy trace in blue, and the [40] occupancy in red. **B** TARJIM’s strongest predicted ArgR motifs, aligned with the [40] motif. The grey shading shows where the TARJIM motif’s primary sequence is the same as that of the RegulonDB motif. **C** The precision-recall curves for the ArgR peaks from the [40] PWM (black) and TARJIM’s predictions (blue). **D** A comparison of the areas under the precision-recall curves as we add each of TARJIM’s strongest motif predictions for the ArgR data in order.

### 3.3 TARJIM permits deconvolution of mixed TF binding signals

We artificially mixed the TrpR and ArgR data sets by taking the peak regions of the TrpR data, multiplying that data by 4, and substituting it for the data in the matching location of the ArgR data set. When we ran TARJIM on this artificially mixed ChIP-chip trace, we found that the number of motifs inferred was more than additive. Because of the relaxed prior on the number of motifs, we inferred 18 motifs on the mixed data set. Our precision-recall curve for the mixture further demonstrates how TARJIM is particularly selective about choosing motifs: the failure mode present in the ArgR motif set with limited motifs is also replicated in the mixture set with limited motifs. Namely, TARJIM motifs generally do not make strong off target binding predictions, even though this also limits the ability to predict positive binding with few motifs, as shown in Figure 3B. The top motifs of the mixture inference mostly corresponded to ArgR’s binding behavior: the top two motifs correspond with an ArgR motif like the [40] motif, as we show in Figure 3C, and it takes eight motifs before we get a conclusive TrpR motif, as we show in Figure 3D. These motifs capture the behavior of ArgR and TrpR, but we note that these motifs are notably more specific than their comparative motifs in the single target binding, as there are more motifs to compensate for other motif hits. The TARJIM inference needs more motifs to match the AUPR of the combined RegulonDB motifs (at least 6 motifs), but the multiple motifs we infer are specifically more selective than they would be in an inference with more stringent requirements for motifs.

**Figure 3.**
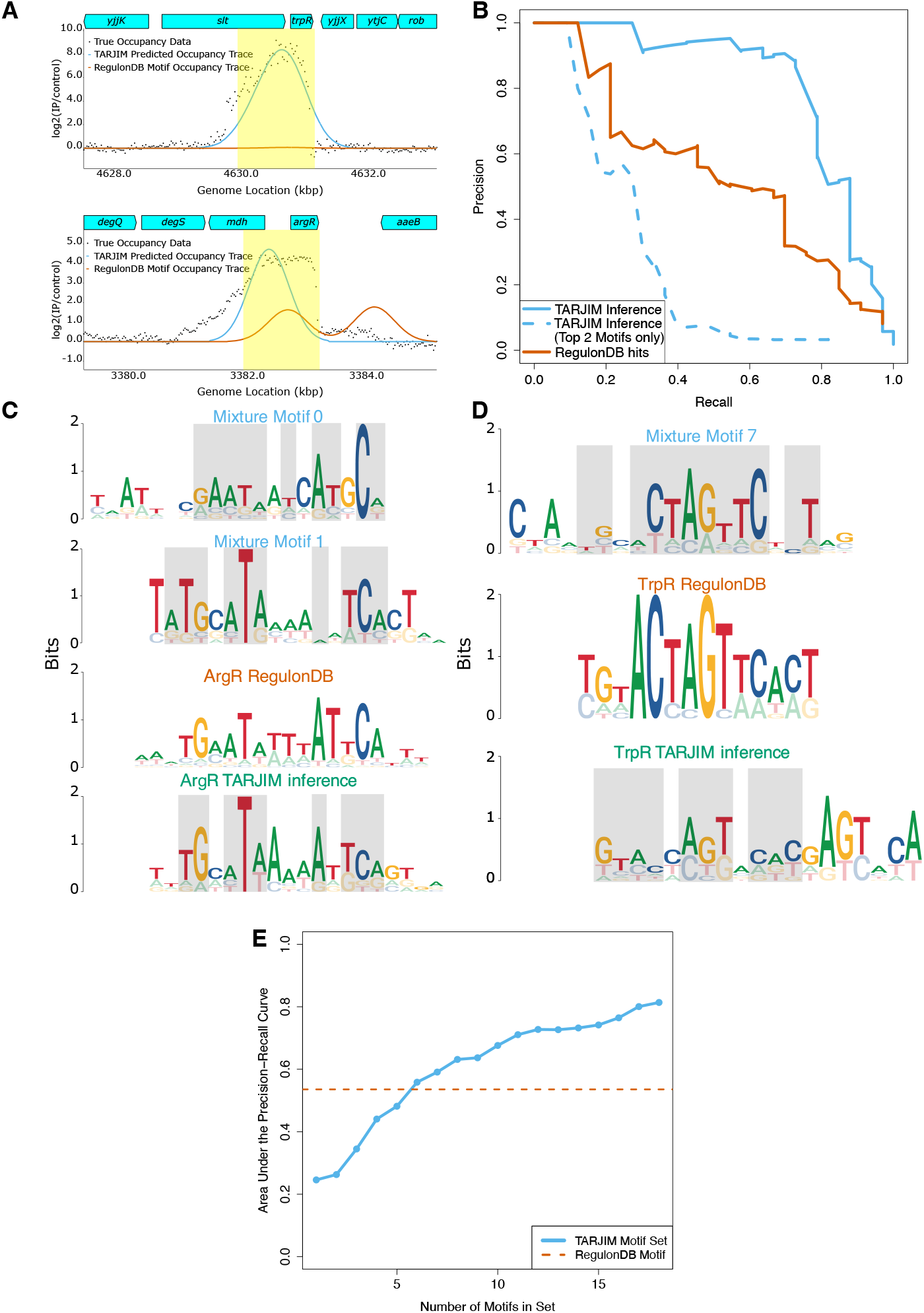
TARJIM’s analysis of the mixed ArgR-TrpR binding data. **A** Predicted binding near the *trpR* locus (top) and *argR* locus (bottom); c.f. Fig. 1A and Fig. 2A. **B** Precision-recall curves of the all of the TARJIM inferred binding sites and the TARJIM inferred binding sites of only the two highest priority motifs, as compared with the combined RegulonDB ArgR-TrpR loci. **C** Motif comparisons of the top two motifs TARJIM inferred from the mixture data (top 2), the ArgR RegulonDB motif (middle), and the top motif TARJIM inferred from the ArgR data alone (bottom). The grey shading shows where the other motifs’ primary sequences is the same as that of the RegulonDB ArgR motif. **D** Motif comparisons of one of the motifs TARJIM inferred from the mixture data (top), the TrpR RegulonDB motif (middle), and the top motif TARJIM inferred from the TrpR data alone (bottom). The grey shading shows where the other motifs’ primary sequences is the same as that of the RegulonDB TrpR motif **E** Plot of AUPRs of the TARJIM inferred motif set as we add motifs as compared to the combined ArgR-TrpR RegulonDB motifs

### 3.4 Lrp

The protein Lrp in *E. coli* is a global transcriptional regulator which is responsive to both the availability of amino acids and the growth state of the cell [28, 50, 51]. It has broad regulatory reach: up to a third of the *E. coli* genome is either directly or indirectly regulated by it [28]. Furthermore, Lrp’s activity is dependent on its exposure to leucine, which can induce a change in oligomerization state in the protein which can result in it changing which genes it regulates [51]. There is evidence that Lrp is both an activator and repressor depending on the genomic context of its binding [28]. Not only are Lrp and other members of the same family broadly conserved amongst bacteria, they also have homologs in archaea [51]. Lrp is a critical protein in bacteria with a great deal of complexity in its binding behavior.

To attempt to understand and model this complex binding behavior, we applied TARJIM to *E. coli* Lrp ChIP-seq data from Kroner et al. [28]. These data include experiments with three nutrient conditions– minimal media (min), minimal media with branched chain amino acids (LIV) and rich defined media (RDM)– taken at three growth points: logarithmic growth (log), stationary phase (stat), and the middle of the transition point from logarithmic to stationary phase (mid). When we performed our inference, it was immediately obvious that TARJIM agrees with [28] that Lrp has variable binding behavior in the three nutrient conditions. TARJIM assigns between 30 and 50 motifs to the min data sets, but only up to 10 motifs for the LIV data sets and about 20 for the RDM sets (Figure 4A). This is clearly far more than the same assignments done by DIVERSITY and MEME-ChIP both, even though both algorithms were allowed to infer at least as many motifs per condition as the largest TARJIM set per condition (DIVERSITY and both the MEME and STREME options for MEME-ChIP were allowed to infer up to 25 motifs for the RDM datasets, 12 for the LIV data sets, and 55 for minimal media data sets). It appears that particularly for a case like Lrp with degenerate binding behavior, TARJIM responds by using a larger set of motifs to build up a reasonable model for the observed occupancy landscape. We calculated AUPRs using the list of peaks published in the Kroner paper for each condition [28]. In general, TARJIM clearly outperformed both algorithms in all data sets except for the LIV log, where its AUPR was worse than MEME-ChIP’s, and LIV mid, where its AUPR was slightly worse than both alternative motif finders. Part of TARJIM’s advantage is the ability to productively infer additional motifs when given the opportunity: while it is better able to predict occupancy than both DIVERSITY and MEME-ChIP (Figure 4A, B), its AUPR suffers when we only restrict TARJIM to only its top motifs (Figure 4D). When we restrict TARJIM to its top 3 motifs, the same number as the MEME-ChIP inference, its AUPR is roughly 0.05, comparable to MEME-ChIP’s AUPR of 0.06. This disparity in the numbers of inferred motifs also affects how well inferred sets can predict not only the peaks identified by [28] from their own data sets, but also the peaks that were identified from other data sets. In general, the areas under the precision-recall curves were highest for peaks and sets that came from the same condition, but the TARJIM inferred set from the min conditions also did relatively well with the peaks identified in the other conditions, while the TARJIM inferred sets from LIV generally did more poorly (Figure 4B). However, we noticed that LIV log inference seemed to do especially poorly, even on its own peaks. When we examined further, we noticed that the top hit from the most important LIV log motif corresponded to a peak near the *waa* operon, and that TARJIM generally inferred the *waa* operon to contain binding peaks in the LIV log data (Figure 4C, top). This top motif was also relatively similar to a motif from the min log data set which also had near the *waa* operon (Figure 4D). However, the data near the *waa* operon were not registered as a peak in the bed files shared by Kroner et al. in any condition, including LIV log or min log [28]. We suspect that this is partially because the shape of the peak is far wider than normal, and partially because by direct admission of Kroner et al., their “ChIP-seq peak calling pipeline is designed to err toward being conservative,” [28]. Lrp does indeed bind the *waa* operon, even in the presence of branched chain amino acids [45], so this hit suggests strongly that TARJIM’s combined motif inference and occupancy trace prediction is able to detect peaks that are unusually shaped but which still seem to be the result of true binding. Several other of the top hits of the TARJIM inferred motifs from the LIV log data are also on locations that are arguably peaks but are not considered such, similar to the results near the *waa* operon. In essence, the LIV log data inference is not one where TARJIM purely fails: instead, TARJIM disagrees with the peak calling pipeline on how the line should be drawn between peak and non-peak.

**Figure 4.**
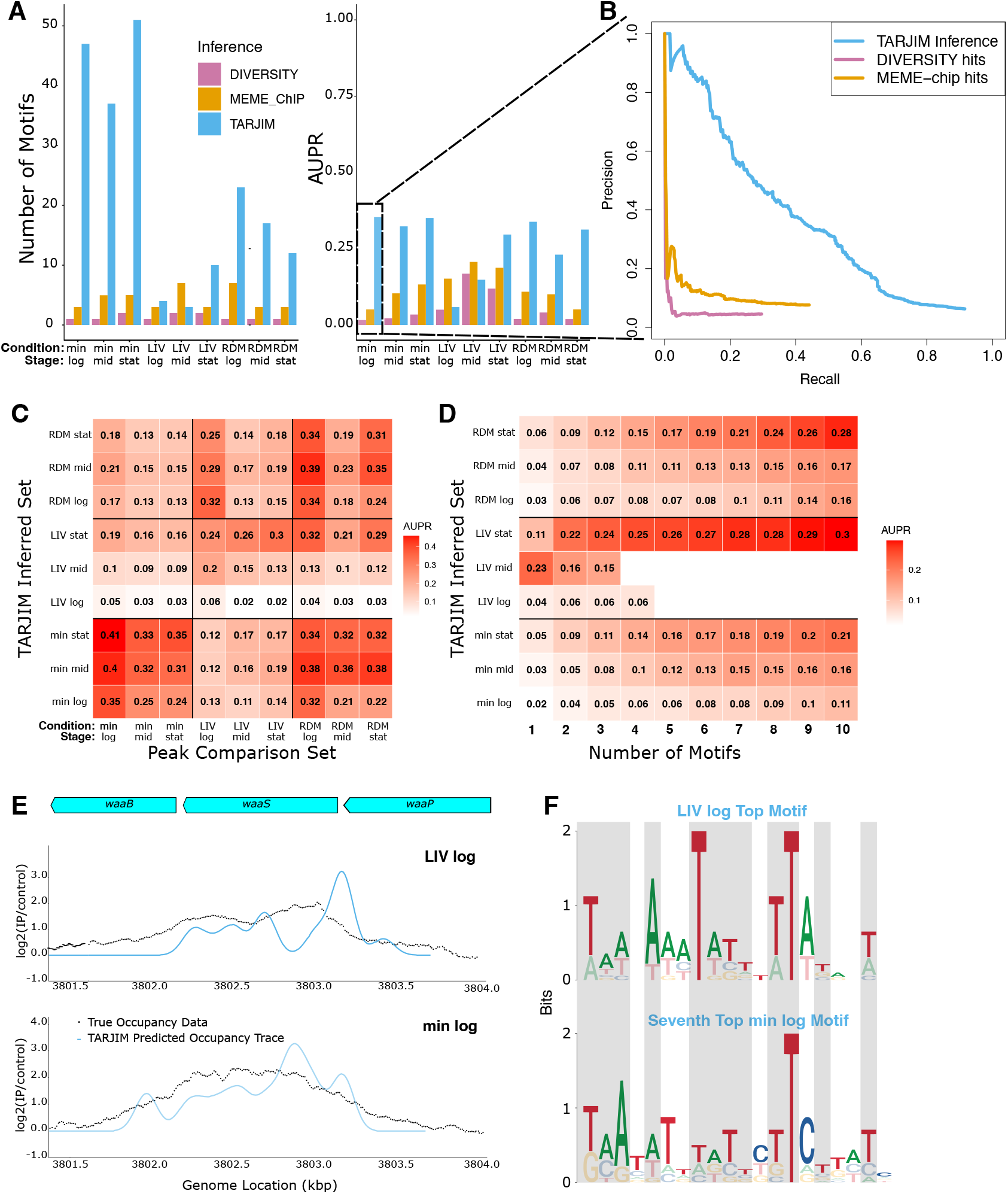
TARJIM’s performance on the Kroner, et al. Lrp ChIP-seq data [28]. **A** Comparisons of TARJIM’s performance on the Lrp data as compared to both DIVERSITY [33] and MEME-ChIP [30], comparing both ultimate motif count (left) and AUPR (right). **B** Sample PR curves comparing the min log inference from TARJIM, DIVERSITY, and MEME-ChIP **C** Areas under the PR curve for each TARJIM inference compared to the peaks seen in Kroner et al. [28] for each data set **D** Areas under the PR curve for each TARJIM inference as a function of number of motifs from each inference included in the PR curve calculation. **E** The TARJIM predicted occupancy traces of the LIV log and min log data near the *waa* operon. **F** The top TARJIM inferred LIV log motif as compared to its closest analogue in the min log motif set. The grey shading shows where motifs’ primary sequences are identical to one another.

We sought to use the TARJIM inferences to better understand the similarities and differences between Lrp binding in different conditions and growth phases. We started by delving manually into examples of how well and poorly TARJIM’s inference performed at a locus that is similar across the Kroner et al. experiments, such as the occupancy upstream of *pepD* and *gpt*, which we show for stationary phase across conditions in Figure 5A. We also wanted to confirm that TARJIM can detect both binding and failure to bind at loci with differential binding behavior, as we demonstrate in Figure 5B upstream of the *lrp* locus itself. During the logarithmic phase, there is clear binding at this locus in minimal media, while the association of Lrp at this locus in rich defined media at logarithmic phase is weak and more consistent with noise than with true binding. In minimal media with branched chain amino acids, the data is in a grey area: the published peak calls from Kroner et al. indicate that a narrow band of the absolute highest value data at this locus is a peak [28], but TARJIM fails to call at a peak at this locus, and it is on the low end of intensity for called peaks. Biologically, it is known that at loci where Lrp has variable binding in response to amino acid conditions, Lrp disassociating from a locus tends to be induced by leucine [51]. The rich media data suggests that the *lrp* locus itself is one of loci where Lrp binding varies in response to amino acid conditions. Thus, we posit that TARJIM may be correct to not call a peak at this locus in the LIV log data: it is biologically consistentthat the presence of leucine would cause Lrp to disassociate from this locus, regardless of the presence of other amino acids. This further reveals a potential strength of TARJIM: if it is ambiguous whether or not locus has a true peak or if increased occupancy at that locus is simply noise, TARJIM can use the sequence information at that peak to help resolve the ambiguity.

**Figure 5.**
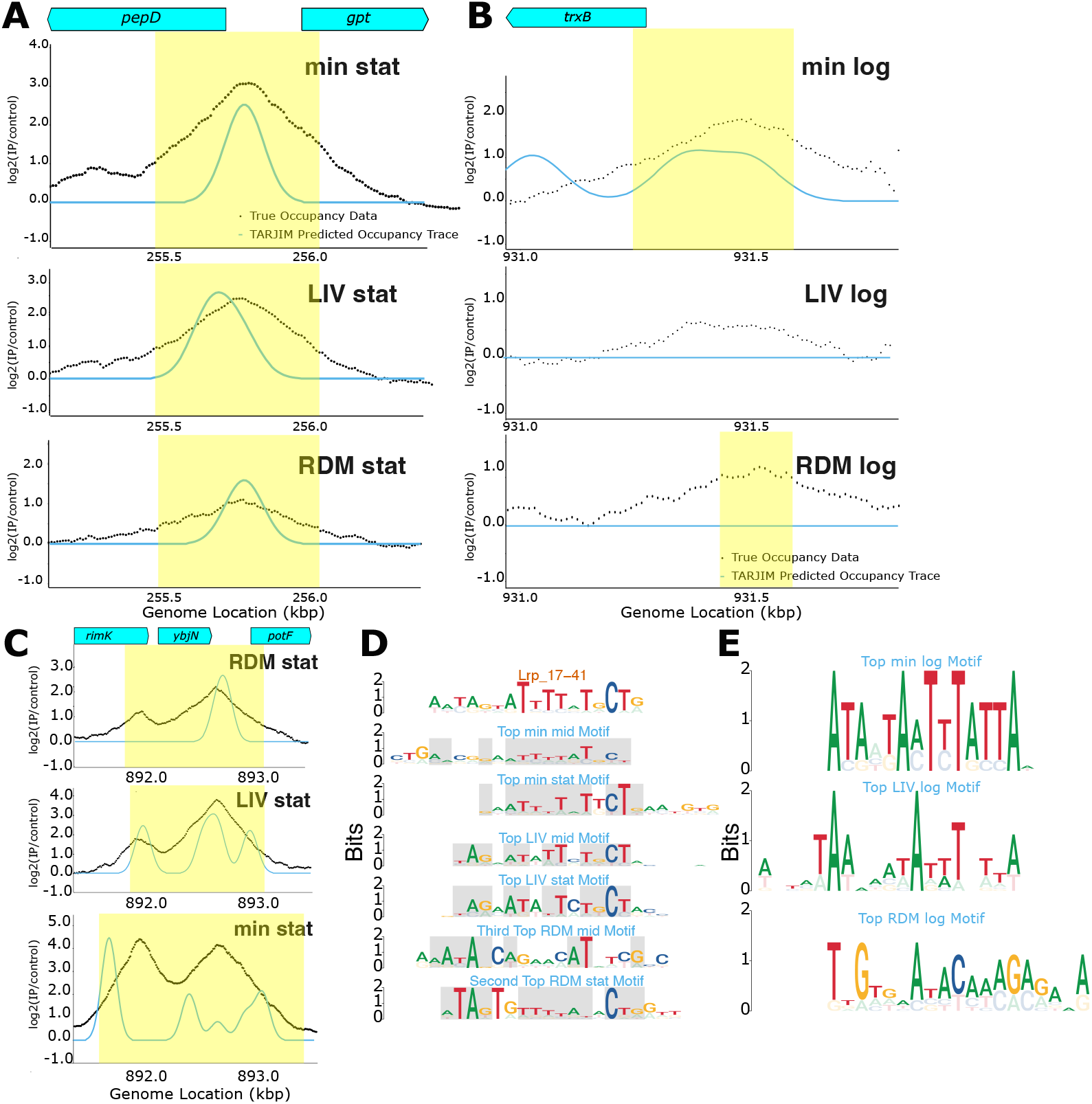
TARJIM’s analysis of the Kroner, et al. Lrp ChIP-seq data [28]. **A** The TARJIM predicted occupancy traces from the stationary phase compared to [28] data near the *pepD* locus, where its occupancy is fairly similar across all experimental conditions. All peak calls published by Kroner, et al. are shaded in yellow in this plot. **B** The TARJIM predicted occupancy traces of the logarithmic phase Lrp ChIP-seq data from Kroner et al. [28], upstream of the *lrp* locus itself, where its binding changes across conditions. **C** The TARJIM predicted occupancy traces of the stationary phase compared Lrp ChIP-seq data from Kroner et al. [28] near the *potF* locus. **D** The TARJIM predicted motifs in the non logarithmic phase experiments matching the SwissRegulon 17-41. The grey shading shows where the other motifs’ primary sequences is the same as that of the SwissRegulon Lrp 17-41 motif **E** The top TARJIM predicted motifs in the logarithmic phase experiments.

However, TARJIM also has clear weaknesses in the face of the Lrp data. The likelihood function for TARJIM specifically prioritizes the shape of the noise distribution in areas with potential peaks. On the one hand, this is good at making sure that TARJIM limits peak calls only to loci with particularly clear evidence of peaks. On the other, this means that TARJIM’s likelihood function prioritizes having matches at those peaks at all, more than precisely matching the shape of the peaks. In the comparatively simple TrpR and ArgR inferences, this was not as large a concern: the data were sufficiently simple that TARJIM’s inferred motifs could fairly precisely match the occupancy trace (Figures 1A, 2A, 3A). However, there are clearly loci in the Lrp inference where this prioritization hurts the inference more, as shown in Figure 5C. The motifs inferred by TARJIM at the *potF* locus in stationary phase are able to correctly call peaks in all three stationary phase experiments, and they even correctly call the binding at the *nfsA* locus in the min stat data. However, with the amount of fitting used for the Lrp data, the inference had not converged to the point of better fitting the shape of the occupancy at *potF* and *nfsA* loci. Nonetheless, TARJIM still detected the peaks there, though imperfectly.

We turn our attention towards the actual motifs inferred across the Lrp datasets, and compare them to Lrp motifs collated by Kroner, et al. [28] by the use of TOMTOM [23]. TARJIM inferred many motifs which were not consistent with any previously known Lrp motif. However, two patterns in the TOMTOM runs on the motif sets stood out. First, across conditions in both the transition and stationary phases, high priority TARJIM motifs matched specifically to Lrp 17-41 from the SwissRegulon database [35] (Figure 5D). In the motifs shown for RDM mid and min mid that TOMTOM matched to the Lrp 17-41 motif, the matches were somewhat weak, with q-values of 0.3 and 0.2, respectively, but Lrp 17-41 was still the best matching motif in the SwissRegulon database. The other, non logarithmic phase data sets’ motifs matched the Lrp 17-41 motif with q-values clearly less than 0.05, and indeed below 0.0001 in the LIV mid data set. For the logarithmic growth phase data, however, there was not a single motif that whose best match was Lrp 17-41. In fact, between the three logarithmic phase data sets, there was exactly one motif which most strongly matched to any specific Lrp motif in the SwissRegulon database at all. However, the top motifs for the logarithmic growth phase data are fairly consistent with being tracts of AT richness (Figure 5E), another indicator of Lrp binding. This suggests that Lrp’s binding behavior shifts to be more specific in *E. coli* as the bacterium transitions from logarithmic growth to stationary phase, a conclusion consistent with the conclusions reached through the manual analysis performed by Kroner et al. [28]. We include the full outputs of TOMTOM on the TARJIM inferences on these Lrp ChIP-seqs in Supplemental File 3, labeled by experimental condition.

### 3.5 IPOD-HR

After our analyses of the Lrp data, we turned our attention to analyzing protein agonistic DNA binding experiments. In particular, we analyzed the IPOD-HR from Freddolino et al. [19]. TARJIM was clearly superior to MEME-ChIP in inferring motifs that reproduce the observed occupancy traces, with TARJIM’s AUPR on the minimal media (rich media) IPOD-HR at 0.19 (0.24), while MEME-ChIP’s AUPR was 0.06 (0.08) (Figure 6A, B, top). We also checked what happened with peak detection if we ignored hits that landed on Extended Protein Occupancy Domains (EPODs), counting them as neither hits nor misses, as those are the loci that are likely least protein specific and that TARJIM is most likely to detect. The AUPR for both motif finders dropped, with TARJIM having an AUPR of 0.11 (0.13) and MEME-ChIP having an AUPR of 0.05 (0.04) (Figure 6A, B, bottom). The observation that TARJIM’s performance sufferred a great deal more when EPOD hits are excluded is unsurprising: the algorithm’s likelihood function is tuned to fit occupancy such as EPODs, while the MEME-ChIP algorithm is tuned to fit identified peaks from a peak finder, which EPODs frequently have the wrong shape for. But even when excluding EPOD hits, TARJIM still performs a great deal better than MEME-ChIP.

**Figure 6.**
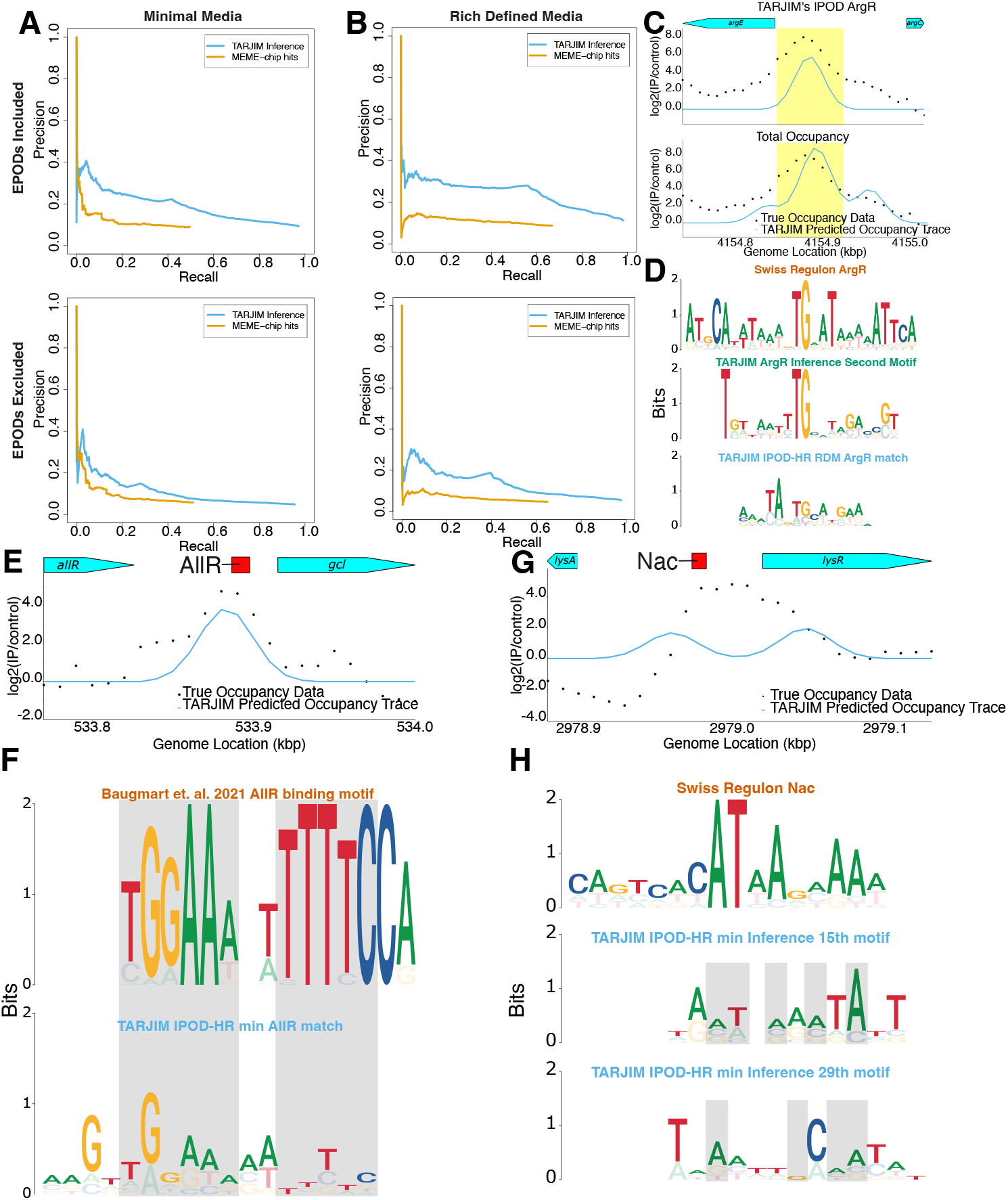
TARJIM’s analysis of the IPOD-HR data from Freddolino et al. [19]. **A** Precision-recall curves for occupancy predictions of the minimal media data, including EPOD matches (top) and excluding them (bottom). **B** Precision-recall curves for occupancy predictions of the rich defined media data, including EPOD matches (top) and excluding them (bottom). **C** The TARJIM predicted occupancy traces of the rich defined media data as compared to the Freddolino et al. [19] data near the *argE* and *argC* loci. All peak calls published by Freddolino, et al. are shaded in yellow in this plot **D** The TARJIM predicted motif from the rich defined media data with occupancy most corresponding to ArgR, as compared to the SwissRegulon ArgR motif [35]. **E** The TARJIM predicted occupancy traces of the minimal media data as compared to the Freddolino et al. [19] data near the *gcl* locus. The red box is the AllR binding site. **F** The TARJIM predicted motifs from the minimal media data with occupancy most corresponding to AllR, as compared to the Baugmart et al. AllR motif [7] The grey shading shows where the motifs’ primary sequences are identical to one another. **G** The TARJIM predicted occupancy traces of the minimal media data as compared to the Freddolino et al. [19] data near the *lysR* locus. The red box is the Nac binding site. **H** The TARJIM predicted motifs from the minimal media data with occupancy near the *lysR* locus, as compared to the SwissRegulon Nac motif [35]. The grey shading shows where the other motifs’ primary sequences is the same as that of the SwissRegulon Nac motif

**Figure 7.**
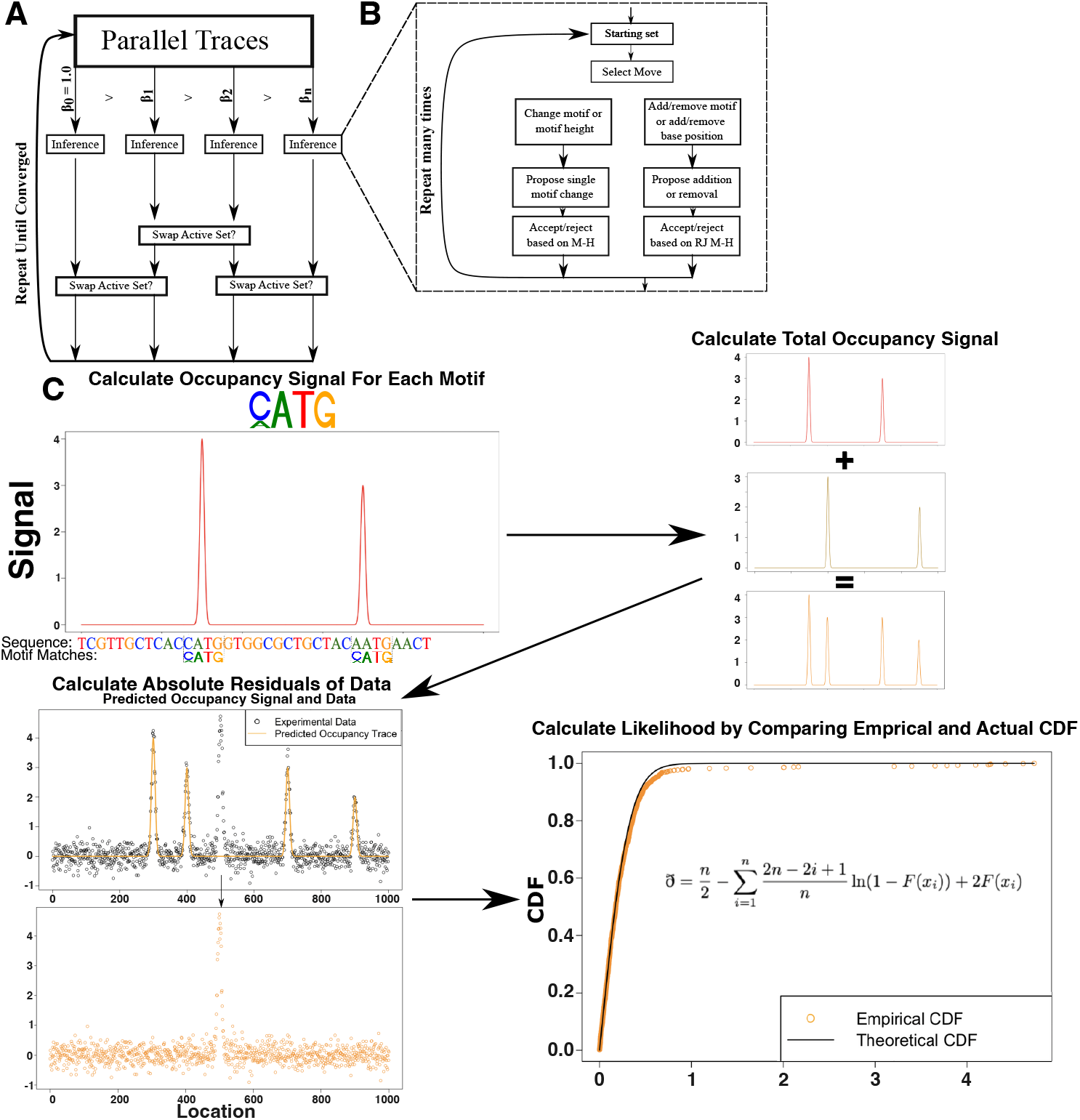
Algorithmic overview of TARJIM. (A) TARJIM’s inference is based first on Replica Exchange Monte Carlo. TARJIM samples from ln posterior distributions with different thermodynamic betas: a thermodynamic beta of 1 is the true distribution, while smaller betas can sample states more easily. At each large step, the betas perform normal inference. Then, they attempt to swap as the replica exchange method [18]. (B) TARJIM’s inference algorithm at each individual thermodynamic beta. At each smaller Monte Carlo step, TARJIM decides to either perform a normal Monte Carlo move, altering its current motif set, or a reversible jump move, changing the number of parameters it is inferring. (C) TARJIM’s likelihood calculation. Given a motif, TARJIM is able to generate a theoretical occupancy trace (top left). It then adds the occupancy traces from each of the other motifs linearly (top right). It takes the total occupancy trace and subtracts it from the data to produce a set of residuals (bottom left). Finally, it takes the absolute value of the residuals and compares their distribution to the background distribution from unoccupied locations (bottom right).

We then wished to understand the motifs TARJIM inferred. We did this by analyzing the occupancies of individual motifs and how clearly they intersected with known transcription factor occupancies from the RegulonDB database [40], as well as comparing the occupancies to the EPODs identified in Freddolino et al. [19]. Importantly, the TARJIM inferences on both sets replicate known differences in transcription factor behavior in different nutrient conditions: the RDM inference has no motifs whatsoever with occupancy near enough to the *lysR* locus to regulate it, and none of the motifs inferred from the minimal media experiment have occupancy most consistent with ArgR over another factor or simply being EPOD associated. On the other hand, the minimal media inference quite clearly has occupancy near the *lysR* locus (Figure 6G, Supplementary Files 1 and 2), and the RDM inference infers a motif whose predicted occupancy is strongly consistent with ArgR (Figure 6D). TARJIM quite clearly is able to predict differential occupancies in different conditions.

For motif identification, TARJIM is most able to extract motifs with high sequence specificity. In the minimal media, for example, there are two motifs out of 134 in the minimal media inference with occupancy near *lysR*: both are most consistent with being EPODs, but the 15th motif of the minimal media inference also generally has a strong overlap in binding with Nac. We thus compare both of those motifs to the SwissRegulon database’s Nac motif [35], and we do not see sequence similarity in motifs despite the occupancy overlap (Figure 6F). As a factor, there is evidence that Nac binds over 400 loci on the E. coli genome [40]. When using the SwissRegulon Nac motif to call motif hits with FIMO [21], the motif calls over 900 hits, only 30 of which intersect these 400 loci. We posit that this is a fundamental weakness of using sequence motifs to capture Nac binding, which TARJIM attempts to compensate for by proposing many, more specific sequence motifs. On the other hand, the 119th motif of the minimal media inference has occupancy most consistent with AllR, which only binds in 3 locations [34]. This includes a locus upstream of *gcl, allRgcl*, which the inferred motif’s occupancy trace captures (Figure 6E). When we compared the TARJIM inferred AllR motif to the AllR motif from the supplementary information in Baugmart et al. [7], we notice that TARJIM’s predicted motif agrees quite strongly: everywhere the motifs overlap, TARJIM’s primary motif agrees exactly with the Baugmart motif, except for the highly ambiguous 7th Baugmart base and a slight disagreement with the 8th Baugmart base (Figure 6F). This increased motif accuracy from TARJIM when transcription factors are more sequence specific is further substantiated when we look at the RDM experiment. From the TARJIM RDM IPOD-HR inference, there are three motifs whose occupancy is most consistent with ArgR. The most important of those motifs is particularly striking: it has strong occupancy near the *argE* and *argC* loci making up the lion’s share of the total TARJIM predicted occupancy at that locus (Figure 6C). Furthermore, the motif also bears a resemblance to both the SwissRegulon ArgR motif and one of the motifs inferred in the ArgR specific TARJIM inference (Figure 6D). Yet, this is clearly not as strong as the resemblance in the minimal media inference’s AllR motif. TARJIM is extremely capable of extracting specific motifs from protein agnostic DNA-protein binding experiments such as IPOD-HR. However, its extraction best matches transcription factors whose binding is sequence specific, such as in the case of AllR (Figure 6F). ArgR has more promiscuous binding, with some sequence specificity, so the TARJIM inference of the ArgR motif from the IPOD-HR data only somewhat resembles both the SwissRegulon motif [35] and TARJIM’s own inference from the ArgR ChIP-chip from Cho et al. [14], as we see in Figure 6D. And when binding at a locus is consistent with EPOD occupancy or more promiscuous factors, TARJIM’s explanations for the binding will suffer, both with the predicted binding matching less perfectly and with the motif inferred not matching known motifs, as shown in Figure 6H. In summary, TARJIM is good at capturing sequence motifs, but performs less well as binding becomes less consistently explained by sequence alone.

## 4 Discussion

As it stands, the problem of transcription factor motif inference has been generally difficult, which has only been exacerbated by the fact that single transcription factors can have complex binding not captured simply by DNA sequences. Motif inference must not only struggle with disentangling binding modes, but also to explain as much of the energy landscape of those binding modes as possible while remaining interpretable. The main algorithms for that could perform this unmixing prior to TARJIM were ChromeBPNet, where transcription factor motifs are a secondary concern and thus have minimal attention to their precision [36], MEME-ChIP, which requires uniformly trimmed sequences and often predicts motif sets with redundancies [30], and DIVERSITY, which requires running separate models for each possible motif count, is not designed to deconvolve binding from multiple transcription factors, and cannot account for where transcription factors specifically do not bind [33].

TARJIM offers a fundamentally different approach: it treats motifs as first class citizens and strongly refines their precision, unlike ChromBPNet, needs only an occupancy trace and a genome to produce a set of motifs where each motif in set accounts for the others, unlike MEME-chip, and it can itself perform inference on the number of motifs represented in a set to minimize the number of models it needs to consider, unlike DIVERSITY. TARJIM’s method of inference also allows it to rank the importance of each motif in the sets it infers, further allowing the user to take the information that they deem relevant. Its output also indicates where each motif binds, and how that binding compares to the actual ChIP data trace, which allows the user to find the motif or motifs explaining particular binding phenomena that they are interested in.

One particular trend we noticed with TARJIM is that it has a tendency to predict substantially more motifs than there are transcription factors. This trend was particularly apparent in the Lrp min stat experiment, which predicted over 50 motifs for a single transcription factor. There are multiple potential reasons for this behavior. First and foremost, binding motifs are an abstraction meant to model energy minima of binding, not serve as a single physical explanation for a binding mode. When TARJIM infers a motif, it can only infer an energy minimum which explains part of an occupancy trace. If it is inferring many motifs, the conclusion is that many different local energy minima are needed to capture the full complexity of the occupancy trace without causing off target binding. There are many potential reasons for this complexity. For example, some transcription factor binding behavior is better explained with the shape of DNA rather than specific sequence motifs [41]. There are many different DNA sequences with similar shapes, and if Lrp is a transcription factor that binds mainly through shape, TARJIM would naturally predict many sequence motifs corresponding to a single shape motif. In addition, because Lrp itself is known to bind promiscuously and differently under different conditions, it is likely has at least two binding modes corresponding to different oligomer states, and it is also known to generally prefer AT richness in the abstract [28, 51]. This sequence limited approach also makes itself especially apparent in the IPOD-HR data: in the rich defined media experiment, TARJIM infers nearly 300 motifs, only about an eighth of which seem to correspond more to specific transcription factors than to EPODs more generally (Supplementary File 2). Ultimately, any researcher using TARJIM on their own binding data would do well to pay attention to motifs that TARJIM reports first when trying to draw conclusions about binding, and/or to perform inference on multiple biological replicates to identify which motifs reoccur across both data sets.

There are important caveats to the results we infer with TARJIM. First and most importantly, TARJIM selects very heavily against motifs that would bind to parts of the genome that a particular data set does not have binding in. This is good for ensuring precision, but it also means that TARJIM has a tendency to prefer multiple, more specific motifs over single, more generic motifs. Second, TARJIM does not account for potential steric or competition effects of transcription factors impacting binding: it assumes that all binding events do not interfere with one another so that it can treat occupancy traces from multiple motifs as additive. This also results in TARJIM having difficulty predicting the existence of a transcription factor motif if that transcription factors binds overlapping a strict subset of another transcription factor: for example, in the context of a global protein occupancy experiment like IPOD-HR, TARJIM would likely never predict a motif consistent with LysR, even in a minimal media experiment where it would be active, as both of its main sites are also occupied by Nac [40]. Third, TARJIM can only directly capture sequence based binding. We see this directly in the minimal medial IPOD-HR inference: AllR has binding most easily captured with a single sequence motif and is easily inferred, while Nac binds more promiscuously and its binding is captured by an ensemble of motifs which do not match its traditional single-motif binding sequence. And finally, the theoretical basis of TARJIM assumes that every base position in a motif independently contributes to the energy of binding. This is frequently not the binding mechanism of transcription factors, where certain bases might correlate with one another to produce an ensemble binding effect [31].

TARJIM is able to perform admirably for the purposes of deconvolving binding from multiple motifs. This makes a tool with potentially powerful applications. First, by applying to the ChIP of global regulators such as Lrp, we can use TARJIM to tease apart different binding modes. Not only might these binding modes simply be selective for different regions, if we compare them with different inferences across different experiments, we can potentially disambiguate the regulation of these different methods of binding. The inferences performed by TARJIM are likely not sufficient for disambiguating all binding. However, by analyzing broad binding landscapes with TARJIM, it becomes possible to orient future analyses of binding behavior: if TARJIM predicts a particular binding motif, it becomes possible to check for it with other experiments, such as with a Gel Shift Mobility Assay [4] or with a PICh experiment [17]. As an algorithm, TARJIM also has a clear avenue for improvement: it is best at capturing binding based on sequence, and requires a great deal more model complexity to capture binding that is not directly sequence based. If a future version of TARJIM incorporates shape motifs [41] or motifs where positions correlate with one another [27], it is likely to improve the interpretability of TARJIM’s results. This will expand the horizon of what our protein-DNA binding experiments can probe for, allowing us to better understand the intricacies of gene regulation.

## 5 Methodology

### 5.1 Data Normalization

Our algorithm generally requires two sources of input data: the genome of the organism used in the experiment, and an occupancy trace which assigns binding propensities to genome locations such as ChIP-chip, ChIP-seq, ATAC-seq, IPOD(-HR), or DNase-seq. The experimental data must have both an experimental condition (e.g., a ChIP pulldown) and a control (e.g., an input sample) to compare against. For every such data set, we take the data values for each location in the experimental setup and divide them by the data values in the corresponding location by the value in the control, and take the binary logarithm of the ratios. For the IPOD-HR data sets specifically, we performed the inference on the robust z scores of these ratios.

For the ChIP-chip experiments, the reference genome was the *E. coli* MG1655 genome (GenBank U00096.2). For the Lrp ChIP-seq experiments, the reference genome was the *E. coli* MG1655 genome (GenBank U00096.2), modified to match the ATCC 47076 variant reported in Freddolino, et al. [20]. For the IPOD-HR data, the reference genome was the current *E. coli* MG1655 genome (GenBank U00096.3).

To determine how to find potential peaks and normalize the data, we fit a “background distribution” to the data. To do this, we fit a normal distribution by the Nelder-Mead algorithm to minimize the square residuals of the densities in the lower three-quarters of the data. From the data, we subtract the mean we derived from this algorithm. As our likelihood function ultimately uses the absolute values of the residuals, the background distribution is set as the half normal distribution where the scale parameter equals the fit normal distribution. We do not normalize the resulting trace further.

Because it would be computationally inefficient to perform all steps of our inference on the whole genome, we narrow in on the parts of the genome with particularly high or low values of data. These binding events influence the data both upstream and downstream of the precise binding location, so we split the potentially bound genome into discrete parts such that each of the parts with potentially binding are isolated from one another. Our preprocessing starts by removing locations that have no associated data. Then, it prunes redundancies by taking the mean of any data that matches to the same location, and finally ensures that the data is uniformly spaced by location through linear interpolation. Our algorithm decides whether to prune between two locations or omit them based on a user supplied fragment length estimate: if two successive locations are more than 2*fragment length+5 base pairs apart, they are separated into independent chunks. The user supplied fragment length as some measure of central tendency for the fragment lengths of their experiment or as roughly half of the width of a typical singlet peak in their occupancy trace.

We take the result and do rudimentary peak finding: if a location has data with an absolute value greater than the data set’s threshold, the one sided *q* = 0.005 value of the background distribution, it is considered a potential peak location, and it and all data points within a fragment length of it in both directions are included in the possible peak data. This specifically includes data with negative peaks: while we do not fit negative peak with TARJIM, we also want to specifically avoid placing positive peaks on top of peaks with negative values. Peak data are sorted into blocks that do not overlap: whenever there is overlap between two or more blocks, they are merged into a single block. We use the term “null sequence” refer to genomic locations which are not contained in one of these blocks, and use the term “sequence block” to refer to genomic windows that are considered potential peak locations. To narrow TARJIM’s search space, a motif is considered invalid, with zero probability density, unless its primary sequence— i.e., the sequence of base pairs it binds with its minimum binding energy—is a sequence of base pairs in a sequence block that can be associated with some positive data point greater than the data set’s threshold.

To perform our inference, we assume that the background distribution is the cause of any difference between a motif set’s theoretical occupancy and the actual data, and the likelihood of any given motif set is given by the probability density that these residuals were indeed generated by said background distribution, as described in more detail below.

### 5.2 Theoretical Occupancy Trace Calculation

We took binding motifs as the primary abstraction by which we characterized binding. These motifs are not necessarily one-to-one with transcription factors. For example, Lrp displays behaviors consistent with several binding modes, [28], while multi protein complexes can be required to come together to bind single DNA location [39]. As we were working with experiments in which bound DNA sequences are primary, we took DNA sequence motifs as our primary driver for binding, and did not attempt to disambiguate this further.

We characterized our binding motifs by using a slightly modified position weight matrix (PWM). Instead of characterizing motifs by information content, we parameterized motifs based on the dimensionless free energy difference 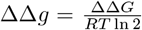 of each position, which we parameterized as *binary energy units*. For any motif, we defined its ΔΔ*g* = 0 as the energy of binding its best possible sequence. For the sake of simplicity, we assumed that the binding energy of each individual position *i* is independent of every other position, and characterized PWMs as being composed of *base vectors*. Each base vector has a ΔΔ*g* = 0 for the best possible base for its position to bind, and all other bases have a ΔΔ*g*_*i,B*_ ≥ 0. We also refer to −ΔΔ*g*_*i,B*_ as the *base penalty* of base *B* in position *i*. We define the *binding score S*_*M*_ (*j*) of a motif *M* of length *L* at a position *j* as

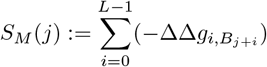

To characterize binding, we modify a result from [48], directly using the formulation based on the free energy of binding. Using this directly—ignoring the simplifying assumption of using the average energy of other sites—the probability that the site at *j* is bound by a transcription factor with a motif M is:

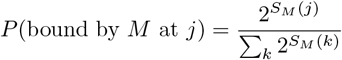

We assume that the ratio of the signals between the experimental condition and the control condition at position *j* is proportional to probability of a motif binding at that position, so long as sequence binding is the primary generator of signal in the experimental condition. Further, we associate every motif with a maximum height *H*(0) of binding, which is defined as the binding signal in a position which perfectly matches it. Because we take the binary logarithm of the signal/control ratio, multiplying the amount of binding 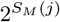 with the maximum amount of binding has the same effect as adding *S*_*M*_ (*j*). Hence, *H*(*S*_*M*_ (*j*)) = *H*(0) + *S*_*M*_ (*j*) so long as motif binding is the primary generator of signal at that position. This requires at least that *S*_*M*_ (*j*) ≥ −*H*(0). The minimum possible *H*(0) in our ChIP (IPOD-HR) inferences is 3.0 (1.0). As such, if *B*_*M*_ (*j*) is smaller than −3.0 (−1.0), then we assume that *H*(*B*_*M*_ (*j*)) = 0.

We assume that a signal of value *H*(*S*_*M*_ (*j*)) ≥ 0 indicates binding, which we represent with a binding kernel in the occupancy trace. The shape of this binding kernel depends on the experiment. In the case of the experiments shown in this paper, we take the binding kernel to be a Gaussian such that 3*σ* is set equal to a user supplied estimated fragment length. If we wished to adapt TARJIM to an experiment like DNAse-seq, we would likely need associate one of several peak shapes to each motif [9].

We assume that binding events are essentially independent of one another, and thus that the occupancy trace created by two motifs is the sum of occupancy traces created by each individual motif, added together. This assumption fails when binding events in sequencing data had a tendency to interfere with one another: while most binding peaks are far enough from one another to ignore this, there can be exceptions, which this tool might be less accurate in capturing [32]. It is important to note that two factors binding the same motif are not problematic if they consistently bind together. However, if two factors are competing for motifs near one another, the linearity assumption is less likely to be accurate. We assume that this situation is rare enough that our inference should be mostly unhindered.

For most of our inferences, we assume that if the total binding free energy of a motif in some position is greater than or equal to 3 binary energy units, the energy minimum this motif represents is no longer the dominant factor influence binding at this position, and thus we do not add any binding kernel to that position. This is a relatively small cut off for our approximation: according to [31], the free energy of binding for a transcription factor binding its best motif might be about 6 binary energy units (roughly 2.5 kcal/mol). However, following this metric precisely would likely be counterproductive: it would naively predict a minimum peak height of roughly 6, which is notably taller than we have observed empirically in the various data sets shown here, and used in development and testing of the method. We also note that, according to [31], it generally takes about three base substitutions for binding to come mostly from nonspecific interactions, which is consistent with how we have defined our model here. However, the IPOD-HR data is meant to capture binding from more transcription factors, and thus there are more DNA fragments competing for detectability, reducing the resolution of a single motif. For the IPOD-HR data sets, we assumed that if the total binding free energy of a motif was at least 1 binary energy unit, we no longer add any binding kernel to that position based on that motif.

To estimate the binding in null sequences where we are not explicitly comparing the predicted trace to the data, we save the sequence information for motif finder locations that do not have noticeable binding in the protein occupancy data. When there is binding at a null sequence, we measure it by finding the Gaussian kernel that this binding would induce and assuming the values from the Gaussian kernel that are at least three times the standard deviation parameter of the background normal distribution to be the residues at that position. We then add them to the full likelihood calculation. This is technically inaccurate if there is a great deal of null sequence binding. However, TARJIM generally selects quite strongly against null sequence binding (see Results for details), ensuring that this situation is not usually relevant for our inference.

We also use this method of calculating null sequence binding to calculate the distance of an occupancy trace from the data: the distance is the square root of the sum of squared residuals in sites with suspected binding and the squared values from ectopic binding Gaussian kernel.

### 5.3 Conversion Between Our Motifs and PWMs

There is technically already a bijection between our motif parameterization and standard position weight matrices. However, if we attempt to do a direct Boltzmann inversion, the PWMs would imply that there are base mismatch positions with an infinite free energy difference, as there are plenty of PWMs which observe no density of binding in any given position. Thus, we convert our motifs to PWMs slightly differently. This type of conversion is usually handled with a pseudocount, but our conversion does not correspond to a single pseudocount across the PWM: instead, the “pseudocount” on a per position basis would be the maximum value of the PWM in each position. This is a consequence of setting our ΔΔ*g* = 0 for best possible sequence for the motif to bind.

We assume that the worst possible energy difference between the best and worst possible base in any given position is 1 binary energy unit, which is equal to *RT* ln(2) = 0.4105 kcal/mol. This corresponds to a single base pair mismatch only being able to halve the amount of binding in a given location at most. To convert our motif parameterizations to PWMs, we take each base identity *b* in each position *i* and compute 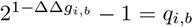 We then compute the probability of seeing a base in each position *p*_*i,b*_ = *q*_*i,b*_*/*Σ_∀*B*_*q*_*i,B*_ to attain the PWM representation for that base. This computation asserts that each base probability is 0 if ΔΔ*g*_*i,b*_ = 1, and that if a probability of seeing a base in a particular position is 1, then that corresponds to a position in the PWM having all maximum ΔΔ*g*_*i,b*_ except for the best base, where ΔΔ*g*_*i,b*_ = 0.

This conversion has physical merit for the ChIP experiments. However, it is less valid for the IPOD-HR experiments, which require a non-physical subtraction of RNA polymerase occupancy. The motifs we display from these inferences are displayed as though this conversion is still valid, but we use a different method to identify what the motifs might truly be (see “Occupancy Based Identification of Transcription Factor Motifs for IPOD-HR” later in this section for details)

### 5.4 Occupancy Based Identification of Transcription Factor Motifs for IPOD-HR

To identify what transcription factors the motifs in the IPOD-HR experiments might correspond to, we took a list of transcription factor occupancies from RegulonDB [40], and for each motif, we calculated the probability that any specific hit of that motif would happen to intersect within 20 base pairs of the sites of each transcription factor. We then calculated p values of each motif corresponding to a transcription factor of interest using the binomial distribution. If *s* is the probability that any specific motif hit would happen to intersect within 20 base pairs of the sites of a transcription of interest, *k* is number of hits of that motif which actually intersected with the transcription factor, and *N* is the number of hits of that motif, then the p-value was calculated as:

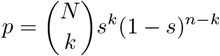

We performed the same calculations with the EPOD occupancies reported in Freddolino et al. [19] as though the minimal media EPODs and rich defined media EPODs were each one transcription factor. For each motif, we identified it with the EPOD or transcription factor with the smallest p-value. For those using TARJIM to identify the specific presence of particular transcription factor binding patterns, we strongly recommend the use of a multiple testing correction for hypothesis testing: we only used this identification to narrow down candidate motifs for manual analysis.

### 5.5 Ln Likelihood Calculation

TARJIM assumes that if it has the complete correct set of motifs causing binding in an experiment, the residual between the theoretical occupancy trace of these motifs and the actual occupancy of the experiment should be noise. The distribution of the noises in each position should therefore match the noise distribution at null sequences: we fit the null sequence distribution to a normal distribution, then subtracted the fit mean from all data points.

We assume that the experimental data is created by this random noise plus some underlying true binding occupancy of protein on DNA. So, we assume the likelihood of a motif set is a function of the probability of the noise necessary to make its theoretical occupancy trace match the experimental data. If we assess the likelihood as the probability of a set of independent samples of a normal background distribution, then the best possible trace would be one that perfectly fit every data point. This ran contrary to our intuition: real life experiments are noisy by definition. In fact, if we found a real life experiment with perfect occupancy traces and no noise, our intuitions would find this alarming and cause us to be skeptical about the experimental setup. Furthermore, early attempts with such a likelihood function frequently inferred tens of motifs even in highly constrained artificial data with only one motif (data not shown), strongly suggesting that this likelihood function would result in an overfit inference.

Thus, instead of attempting to maximize the likelihood of the individual observations, we sought to maximize the likelihood of the aggregate noise distribution in the experiment. There are multiple established methods to do this, but many were ill suited to this problem. The Kolmogorov-Smirnov statistic [2] is the maximum difference between the empirical cumulative density functions of the noises as compared to the theoretically correct distribution, but this statistic is most sensitive near the mean in the distribution, while we want to specifically capture deviations in the tails. The Cramer von-Mises statistic [15] is the sum of the deviations across the statistic, so it is more sensitive than the Kolmogorov-Smirnov statistic, but ultimately is still most sensitive to the middle of the distribution. The Anderson-Darling statistic [3] is a weighted sum of distances in the probability density function, designed to be more sensitive to deviations in the tails, but it was not helpful in capturing the dynamics of the occupancy trace. In particular, when we attempted to use the AD statistic on the signed noises of the distribution in our motif finder, the motif finder maximized the fit to the noise distribution by “compensating” for peaks it could not easily fit with peaks that were specifically located in parts of the genomes without binding, as this would create noises of the opposite sign, making the empirical noise distribution more symmetric and better fitting the symmetrical background noise distribution. When we attempted to use the Anderson Darling on the absolute values of the noises instead, the motif finder displayed erratic behaviors in its fit, as the Anderson Darling statistic does not converge for probability density functions with nonzero density at their tails: this includes the half normal distribution that arises by taking the absolute value of normally distributed noises. We needed a statistic that could work on the unsigned noises, with high sensitivity at only the upper tail. Thus, we used the upper tail modified Anderson Darling statistic first derived and used by Ahmad, Sinclair, and Spurr [1, 43]. For the sake of convenience, we call it the “Ahmad statistic” and label it with the letter ð (“eth”). For a set of data points *x*_1_, *x*_2_, …*x*_*n*_ ordered from smallest to greatest that supposedly follow a distribution with CDF *F* (*x*_*i*_), it has the following form:

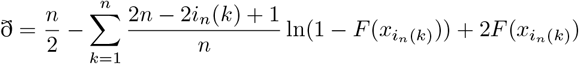

To calculate the likelihood of a particular set of motifs given an occupancy trace, we take the residuals of the predicted occupancy against the true occupancy and then calculate the probability density of that the residuals match the theoretical background distribution of noise.

We use the likelihood calculated from the Ahmad statistic to compute a posterior density for any particular set of motifs, and then use a Metropolis Hastings with Reversible Jump algorithm to iteratively improve our motif set estimate. The ability to perform reversible jump moves is vital for TARJIM: it allows it to identify not only what the key motifs to explain the data are, but also to infer the number of motifs likely to be present in the data itself.

We found that it would be impractical to attempt to find an analytical expression for the probability density of the Ahmad statistic. We could only find an analytic expression for its characteristic function, which gives following form for its PDF:

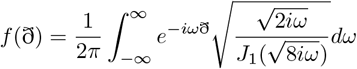

Where *J*_1_(*z*) is the first order Bessel function of the first kind. We were unable to find or derive an analytical solution to this integral. Hence, we used numerical approximations instead.

For the upper tail of the Ahmad statistic, we use the same approximation that Ahmad, Sinclair, and Spurr used, based on Zolotarev’s approximation, to derive the approximate ln PDF of ð [1, 43, 52], which comes out to:

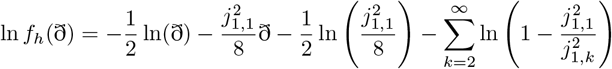

Where *j*_1,*k*_ is the *k*th smallest positive zero of *J*_1_. We make the following approximations:

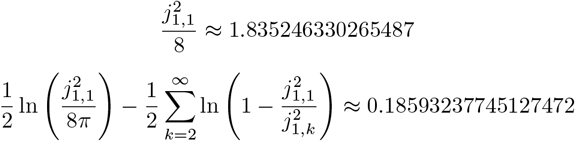

We thus use the following function for the high tail of the natural logarithm of the PDF:

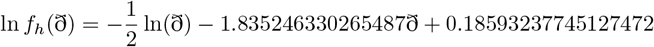

For the low tail, we took advantage of the fact that the characteristic function, in product form, does match a known random generating process: it is a linear combination of infinite chi square distributed random variables with one degree of freedom and coefficients of 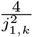. We simulated this distribution ten thousand times using the first one million coefficients (data not shown), and fit a function of the form:

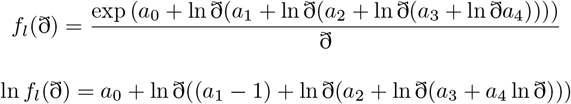

With our approximations, we obtained that

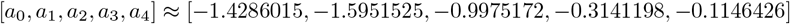

For the sake of continuity in our likelihood function, we defined a weight function *w*(ð) = (1+(ð*/*1.97)^16^)^−1^ to transition smoothly between the low and high tails of the approximation. We observed that the weight function was definitely less than a 64 bit machine epsilon when ð ≥ 20, so we defined our full function as

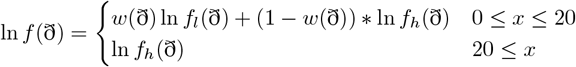

In all practical cases we have observed in the inference, we exclusively operate in the ð ≥ 20 regime.

#### 5.5.1 The Ahmad Statistic Composes

We wish to focus our inference on only the parts of the data containing peaks, which should comprise a minority of the data. We must establish that, if most of the data in a set comes from the background distribution, with only a minority of deviant data points (the peaks), it is mostly accurate to compute ð based calculating solely on the deviant data points. We refer to this as the Ahmad statistic *composing*.

Suppose we have a set of *n* + *j* points and a set of *n* points, such that the first *n* points of the *n* + *k* points are the same points as the set of *n* points. If we declare *i*_*n*_(*k*) to be the order of the *k*th data point for the set with *n* points, the ð of the smaller set is:

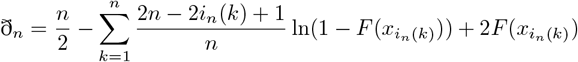

And for the larger set of data, this statistic is:

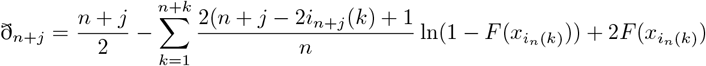

Thus:

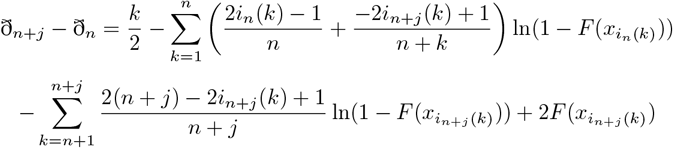

If *k* ≤≤ *n*, then 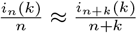 for points shared between the two sets. Furthermore, 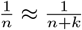 So:

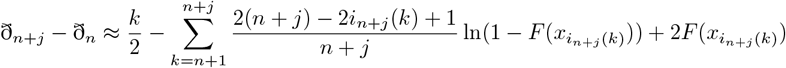

So, as long we do not have a large amount of off target binding as compared to the length of our genome, ð can simply be calculated as mostly just the difference of the occupancy traces in the binding parts of the genome, plus the extra from the off target binding. In general, for higher likelihood motif sets, TARJIM selects against off target binding, so this is generally a safe approximation.

### 5.6 General Algorithm For Motif Inference

Our algorithm is based on several versions of the Metropolis-Hastings algorithm [24]. Given some starting set of motifs, we have several moves which propose a new set of motifs to sample. Note that motif sets are unordered during inference: a motif set with motifs {*A, B, C*} is identical to a motif set with motifs {*B, A, C*}. We report motifs in a particular order based on their contribution to the likelihood function, but this is a post processing convention, and if that reporting order would give a motif sequence (*C, A, B*), it would give that sequence regardless of the permutation of {*A, B, C*} that is present in the actual inference.

Because we must account for the fact that we do not know *a priori* precisely how many motifs are needed to account for the binding events in a set of sequencing data, we must resort to Reversible Jump Markov Chain Monte Carlo (RJMCMC) [22]. This algorithm will allow us to infer both the number and identity of motifs in our inference in a fully Bayesian framework.

Below we enumerate the move types used in our method, followed by the overall algorithm followed for sampling from the posterior distribution on our motif sets. Note that we use *ϕ*(*x*) as the PDF of the standard normal distribution at a real number *x*, Φ(*x*) as the CDF of the standard normal distribution evaluated at a real number *x*, and Φ^−1^(*p*) as the inverse of this CDF evaluated at a probability *p*.

#### 5.6.1 Motif Scramble Moves

We have three moves which scramble motifs’ base vectors.

First is a “base leap” move. This move tries to change the primary sequence of a motif to another valid primary sequence within a maximum Hamming distance, generally preferring primary sequences close to the current motif’s primary sequence. This maximum Hamming distance is chosen so that we are generally choosing from approximately 200 possible primary sequences, based on the distribution of valid primary sequences of each motif length in a given data set. We balance this move by explicitly calculating the probability of the reverse proposal, as required to maintain detailed balance.

##### Algorithm 1

Base leap move

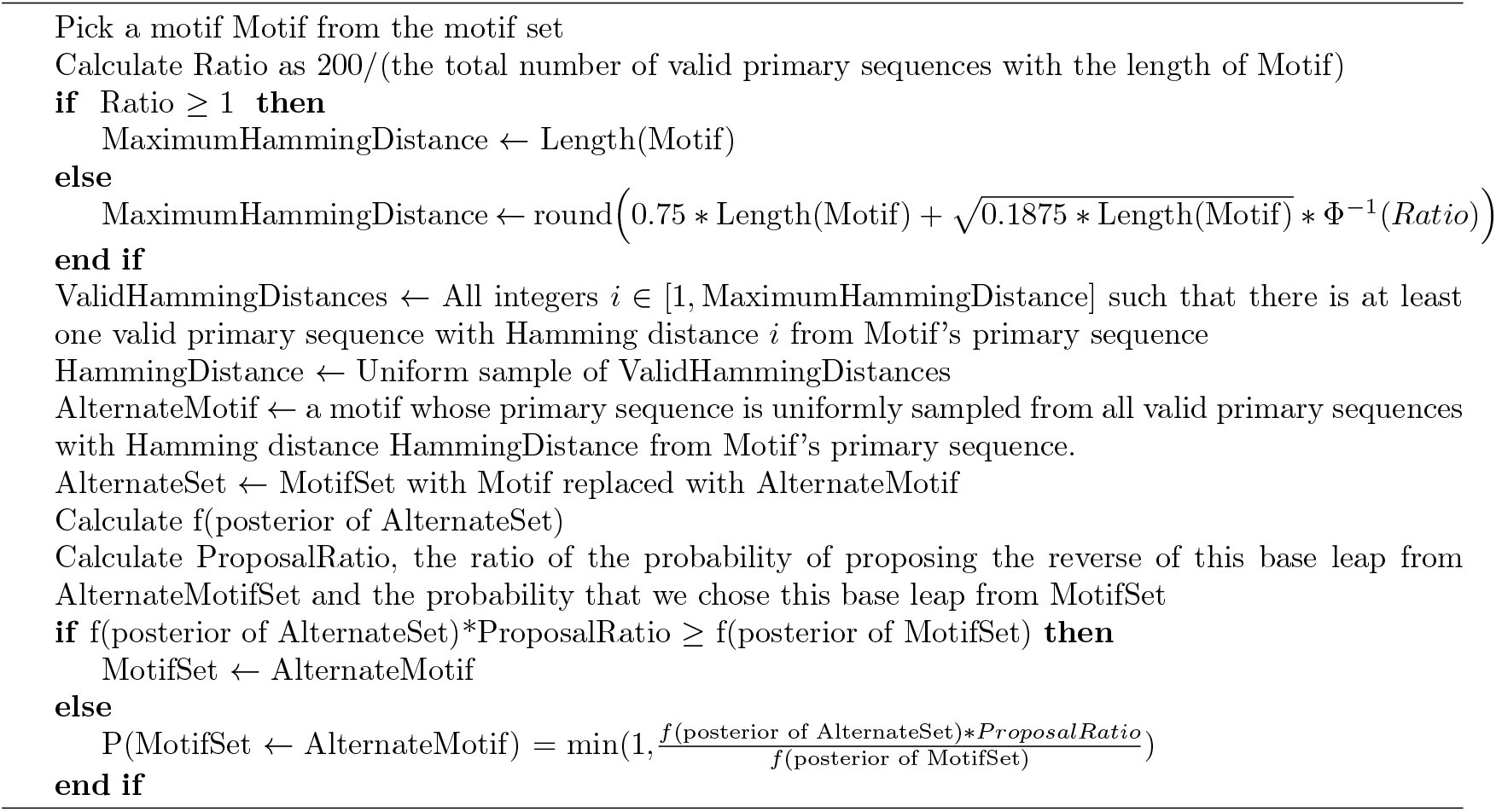

Second is a “motif by residuals” move, which is the other way that a motif can change its best base in its positions. Instead of preferring a motif based on closeness to a current motif, it instead attempts to select motifs that are likely to correspond to peaks. We balance this move by explicitly calculating the probability of the reverse proposal, as required to maintain detailed balance.

##### Algorithm 2

Motif residuals move

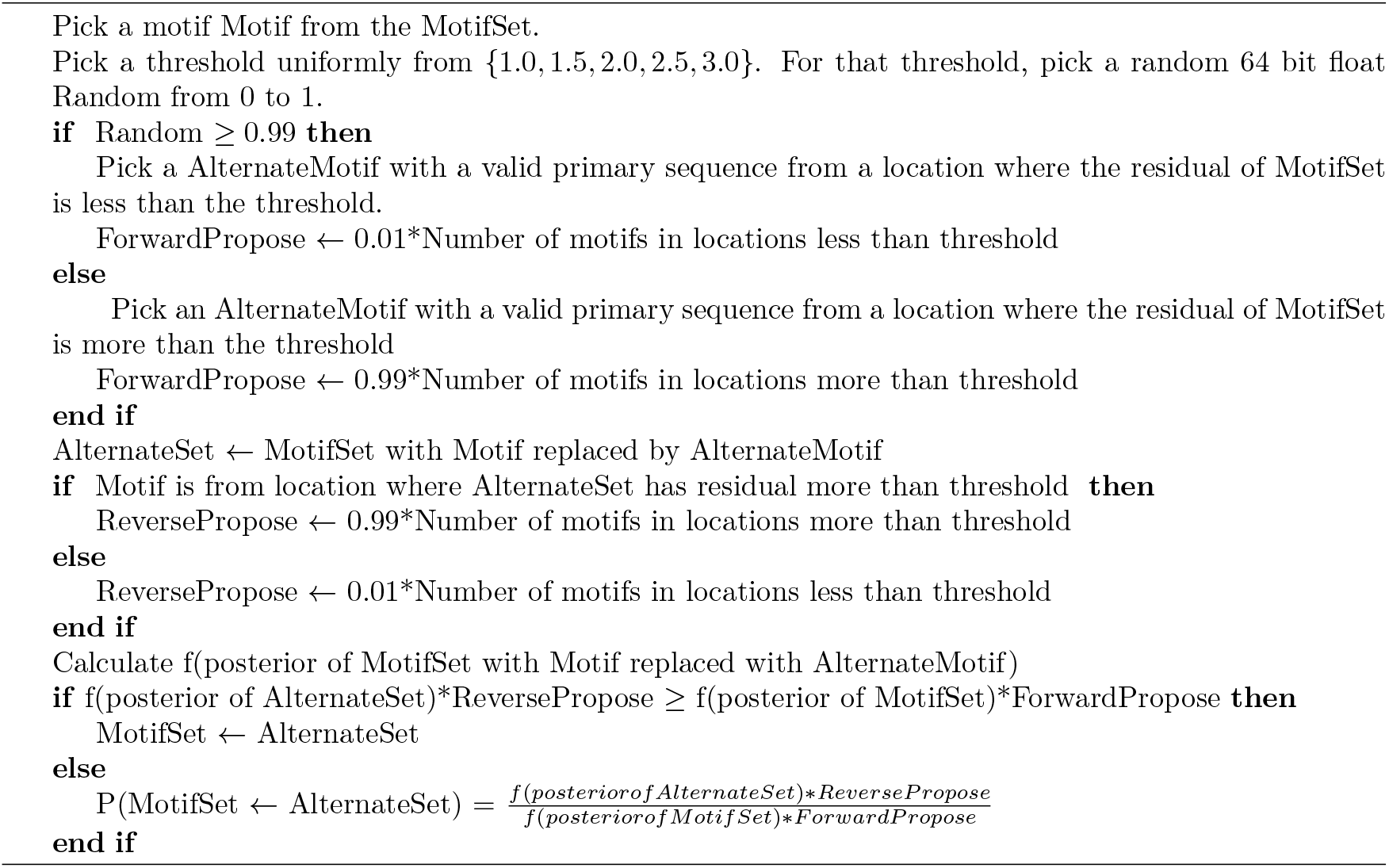

We have a “secondary shuffle” move, which attempts to change scramble of a motif’s penalties without changing its primary sequence. It is meant to accelerate convergence by allowing a motif to vary quickly without changing its primary sequence.

##### Algorithm 3

Secondary shuffle move Pick a motif Motif from the motif set

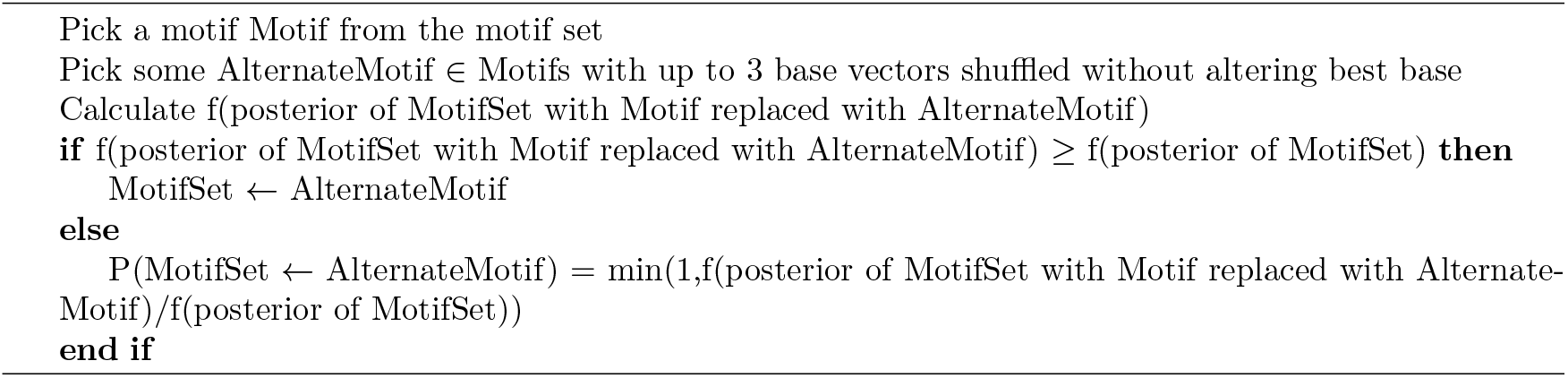

#### 5.6.2 Other Metropolis Hastings Moves

We also have moves which follow more standard Metropolis Hastings. These moves all work by proposing a motif set such that all but one motif is identical to the current motif set, with that one motif modified in some way.

One attempts to change a single energy penalty of a motif.

##### Algorithm 4

Penalty flip move

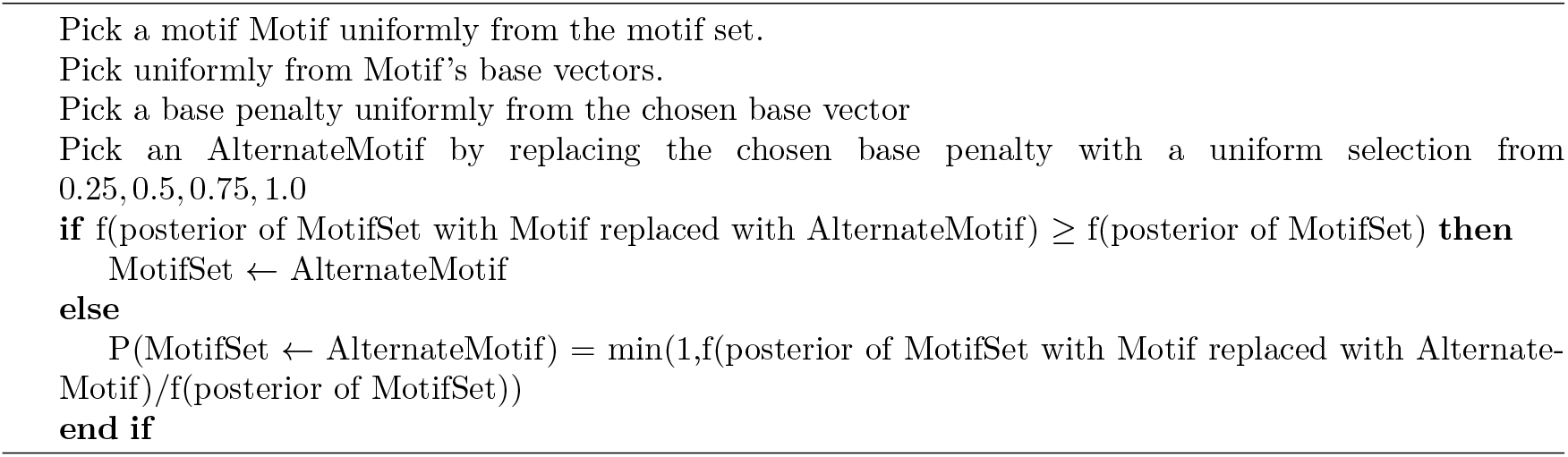

Another attempts to change the maximum height of a motif:

##### Algorithm 5

Motif height move

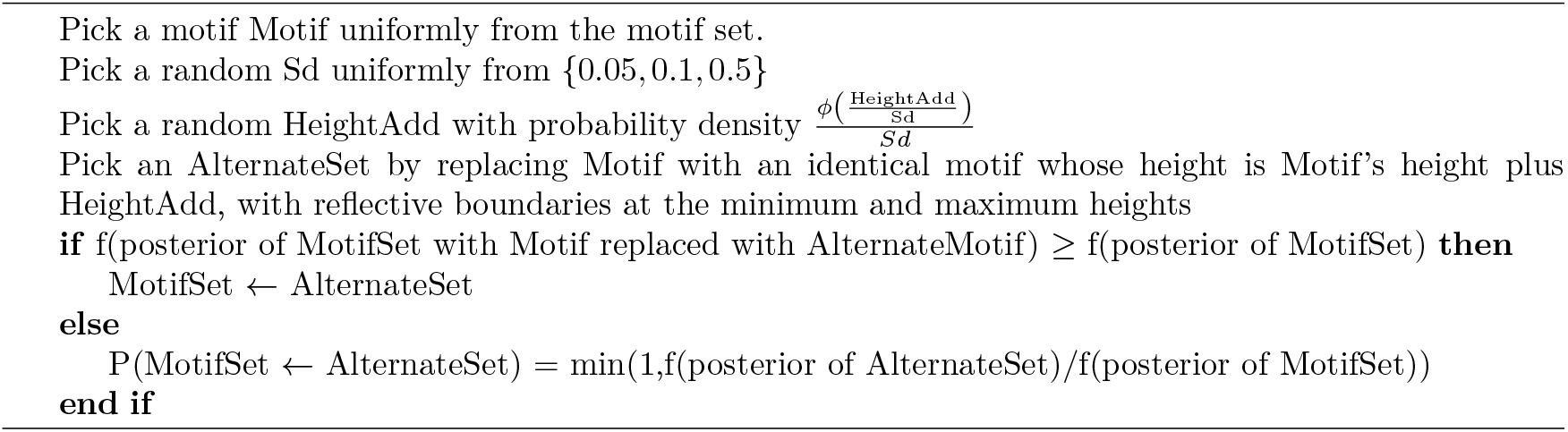

#### 5.6.3 Reversible Jump Moves

To account for the possibility of varying numbers of transcription factors with varying lengths, we include two pairs of Reversible Jump moves, based on [22]. This modifies the odds ratio to account for both a prior on the hyperparameter (in our case, the number of transcription factors and their lengths), and to account for any changes in geometry between changes in dimensionality, accounting for random variables used in the generation of larger sets from smaller ones. These moves are always balanced by explicitly calculating the probability of their reverse proposals, as required to maintain detailed balance.

We have one pair of reversible jump moves which changes our motif lengths by adding or removing a base vector from either end of a single motif. The extension move is:

##### Algorithm 6

Motif extension move

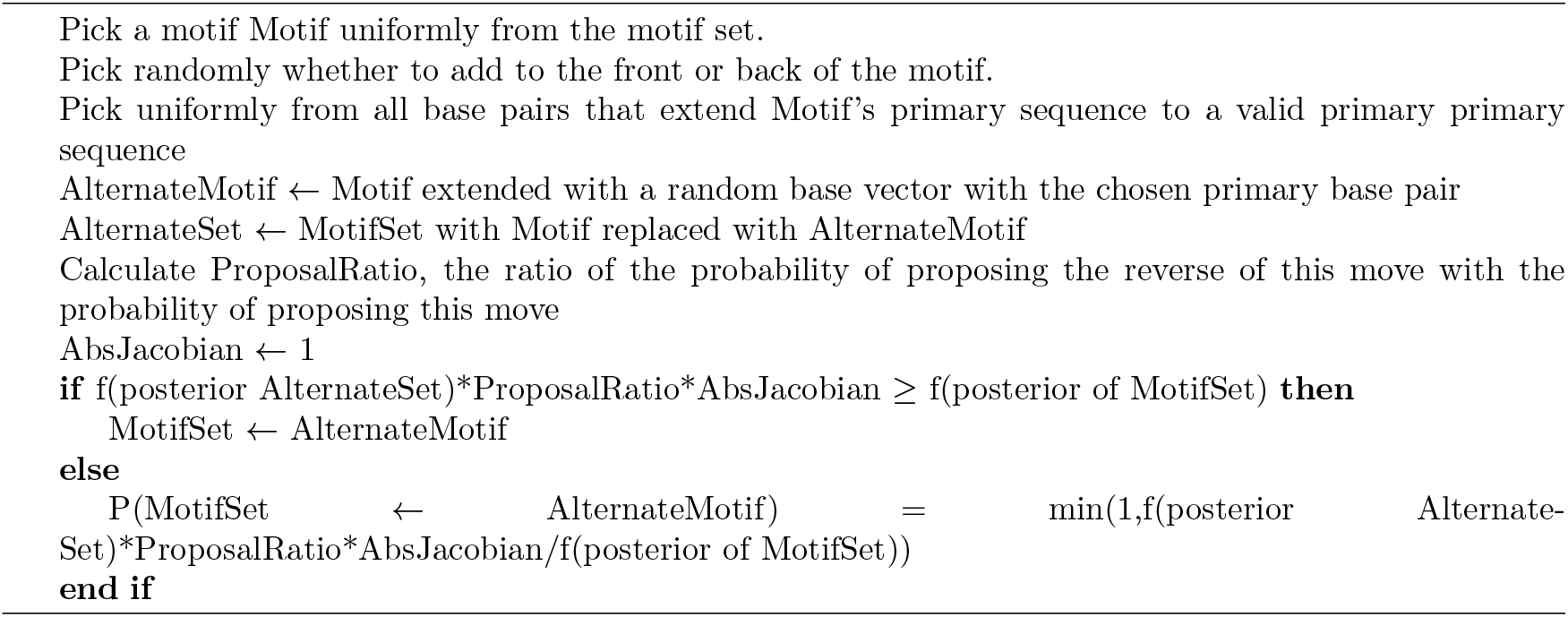

The contraction move which reverses the extension move is as follows. We note that, in the version of TARJIM used in this paper, there was a small programming error which could very rarely cause the contraction move to be rejected when it might have been accepted. This error does not affect any of the results, as we used TARJIM solely as a search algorithm. We have corrected this error, in case other users want to use TARJIM to generate full posterior distributions.

##### Algorithm 7

Motif contraction move

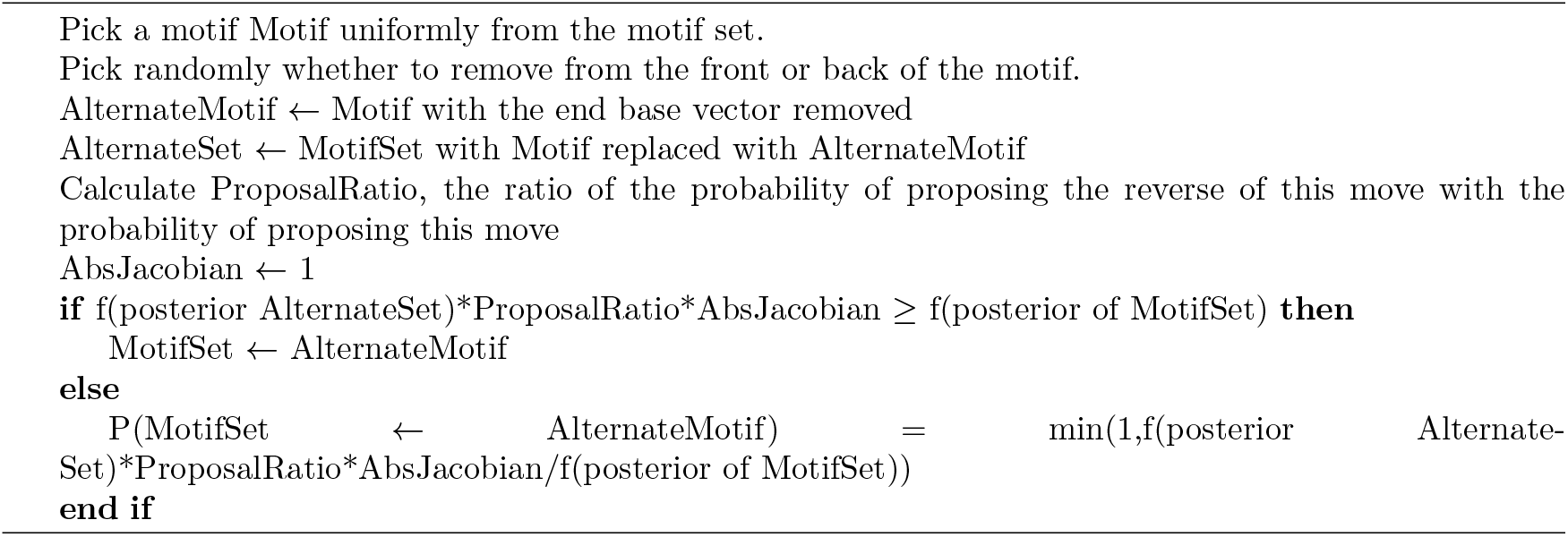

We have two pairs of reversible jump moves which add or remove motifs. Both depend on generating a motif length, then a valid primary sequence of that length, and then a motif with that primary sequence. However, one chooses a random valid primary of the requisite length directly, while the other uniformly chooses a random 8mer and then uniformly chooses a random valid primary with the requisite length that begins with that 8mer. The motif addition moves are otherwise identical, and have the following algorithm:

##### Algorithm 8

Motif addition moves

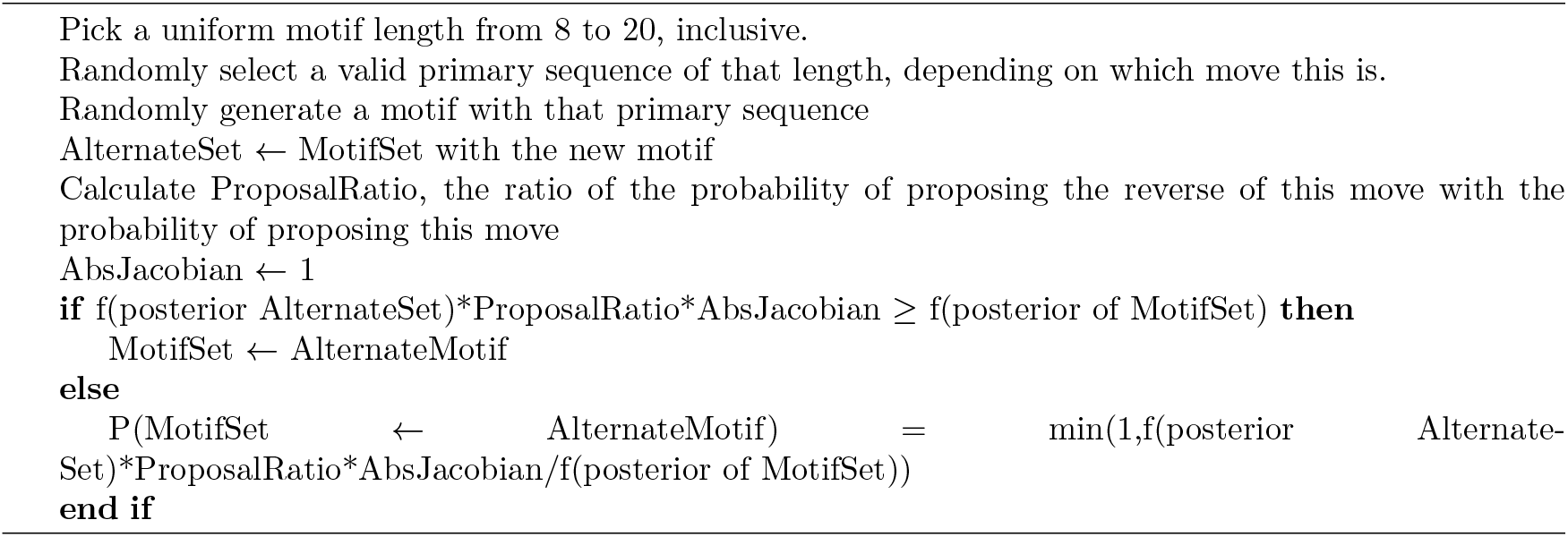

The motif removal moves are algorithmically identical, save for the calculating the ratio of the proposal of their reverse move to their forward move, and have the following algorithm:

##### Algorithm 9

Motif removal moves

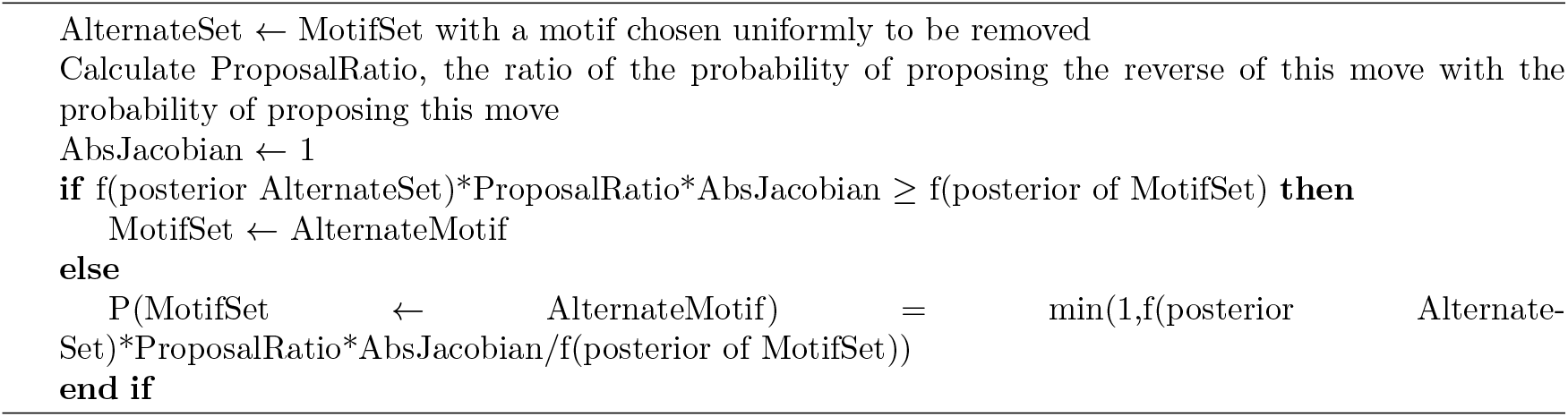

### 5.7 Priors

The prior on the number of motifs is set to be a geometric distribution with a support of the positive integers. However, the specifics of this prior should be set by the users prior expectations of the maximum number of motifs that should be represented in a particular binding experiment. The probability parameter *p* is chosen by the user by considering the a number of motifs *N* that the user thinks should be an approximation of the number of motifs in the experiment. The probability parameter is then set by the relation 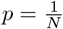 if *N* ≥ 1. If *N* = 1, the parameter should be chosen such that *p* approaches 1. For the TrpR and ArgR inferences, we set *p* = 1 − exp(−50), on the assumption that one transcription factor should generally have 1 motif. For the mixture inference and the Lrp inferences, we set *p* = 0.5, on the assumption we are expecting roughly 2 motifs. For the IPOD-HR inferences, we set *p* = 0.005, on the assumption that we are expecting roughly 200 motifs.

We set a uniform prior on the motif lengths, so that the length *L* of a motif is uniformly distributed on the integers in [8, 20].

A motif’s probability density is set to 0 if its primary sequence is invalid or if it does not have measurable binding in at least two positions, each found in some sequence block—this can be two occurrences of its primary sequence, or one occurrence of its primary sequence alongside another site with weaker binding. This probability density is also set to 0 if the motif would induce more null sequence binding than some maximum: 1000 for all the inferences other than the IPOD-HR inferences, where this maximum was set to 200 instead. For the ChIP inferences, this prior was simply a programmatic limitation which did not impact the final result, as the inference strongly selected against null sequence binding. However, this prior did impact inference for the IPOD-HR inferences.

For all motifs whose probability densities are not automatically set to 0, the motif prior is proportional to a product of 3*L* uniform distributions on the base penalties [− 0.25, −0.5, −0.75, −1.0] for all of the other base pairs and positions. We did this because we observed empirically that binding behavior tended to change in discrete steps far more than in smooth changes, most likely because of our discrete cutoff for binding.

The distribution of motif heights *H* is set to follow a truncated log normal distribution with ln *µ* = 10, ln *σ* = 0.25, and limited to *H* ∈ [3, 15] for the ChIP inferences. For the IPOD inferences, we used the same distribution except for using the truncation *H* ∈ [1, 15]

### 5.8 Accelerating Convergence

For all the inferences we show, we ran 4 independent identical Monte Carlo chains at the same time. Each chain used replica exchange Monte Carlo [18] with 128 different thermodynamic *β*s, with a true chain with *β*_0_ = 1 and supplementary chains such that the *i*th chain had a 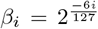 for *i* ∈ [0, 127] ∩ ℤ. We ran individual chains using 200 iterations of all previously mentioned Metropolis Hastings moves, randomly chosen at each of the iterations, then calculated swaps using the replica exchange criterion. We ran the ChIP inferences for 6 days, with break points every 2 days where we reinitialized the replica exchanges with the motif set from the chain with unit thermodynamic beta. For the IPOD-HR inferences, we ran for 10 days, with the same break point and reinitialization pattern. All inferences were run on 127 cores of an AMD EPYC 7742 on EXPANSE. The approximate number of Monte Carlo steps for each independent chain and inference is reported in this table:

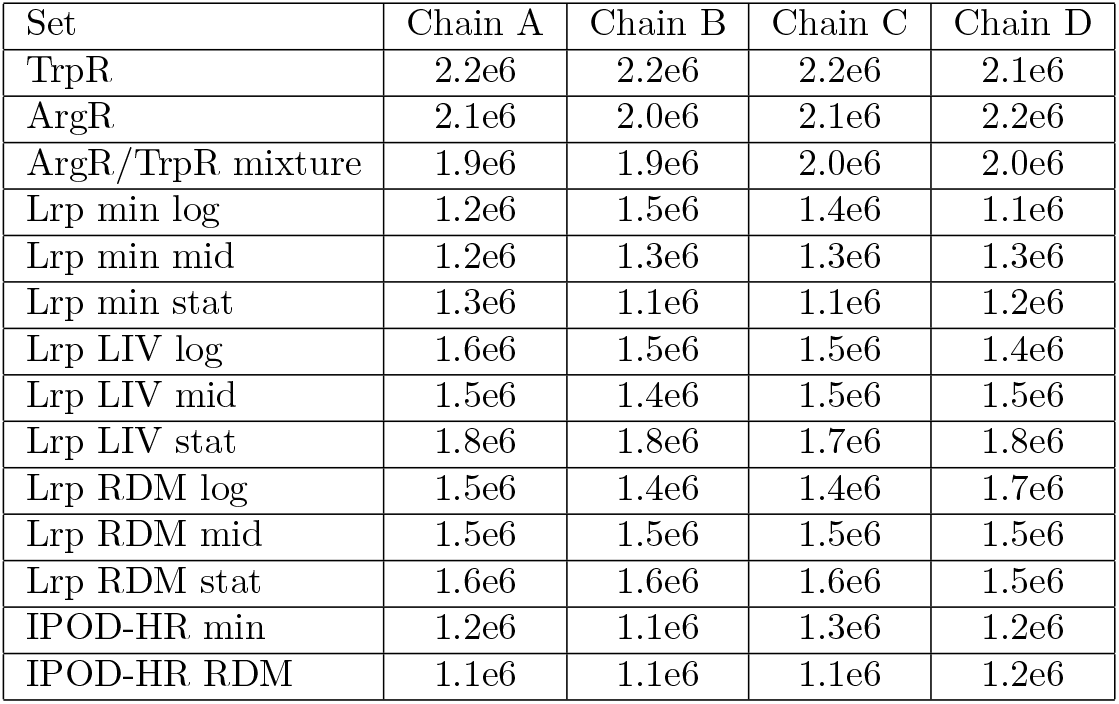

| Set | Chain A | Chain B | Chain C | Chain D |
| --- | --- | --- | --- | --- |
| TrpR | 2.2e6 | 2.2e6 | 2.2e6 | 2.1e6 |
| ArgR | 2.1e6 | 2.0e6 | 2.1e6 | 2.2e6 |
| ArgR/TrpR mixture | 1.9e6 | 1.9e6 | 2.0e6 | 2.0e6 |
| Lrp min log | 1.2e6 | 1.5e6 | 1.4e6 | 1.1e6 |
| Lrp min mid | 1.2e6 | 1.3e6 | 1.3e6 | 1.3e6 |
| Lrp min stat | 1.3e6 | 1.1e6 | 1.1e6 | 1.2e6 |
| Lrp LIV log | 1.6e6 | 1.5e6 | 1.5e6 | 1.4e6 |
| Lrp LIV mid | 1.5e6 | 1.4e6 | 1.5e6 | 1.5e6 |
| Lrp LIV stat | 1.8e6 | 1.8e6 | 1.7e6 | 1.8e6 |
| Lrp RDM log | 1.5e6 | 1.4e6 | 1.4e6 | 1.7e6 |
| Lrp RDM mid | 1.5e6 | 1.5e6 | 1.5e6 | 1.5e6 |
| Lrp RDM stat | 1.6e6 | 1.6e6 | 1.6e6 | 1.5e6 |
| IPOD-HR min | 1.2e6 | 1.1e6 | 1.3e6 | 1.2e6 |
| IPOD-HR RDM | 1.1e6 | 1.1e6 | 1.1e6 | 1.2e6 |

### 5.9 Parallel tempering automatically accounts for reversible jumps

We found other examples of parallel tempering used to accelerate convergence for reversible jump Metropolis Hastings (e.g. [8]), but did not find a clear proof that swapping different dimensional Monte Carlo states between chains does not necessitate additional consideration. To ensure the record is clear, we present this proof here:

Suppose we are considering a swap between two chains with thermodynamic betas *β*_0_ and *β*_1_, with motif sets *M*_0_ (with dimensionality *N*_0_) and *M*_1_ (with dimensionality *N*_1_), respectively. We denote *c*_0_(*M*_0_) to mean “the chain with thermodynamic beta *β*_0_ has motif set *M*_0_ as its current motif set.” If our posterior distribution is denoted *π*(*M*), then to maintain detailed balance in both chains, the following equation must hold:

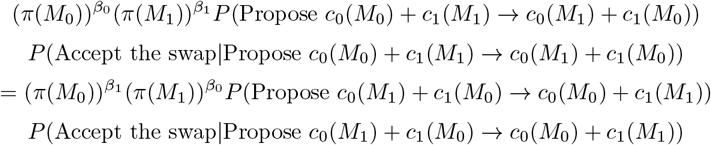

The probability of proposing a replica exchange swap is constant between the *b*_0_ and *b*_1_ chains, regardless of motif set. Cancelling the proposal term and simplifying:

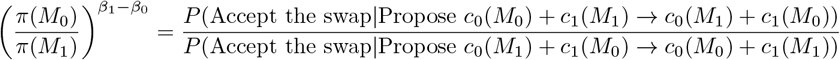

Which is exactly the parallel tempering criterion.

We observe that at no point did the dimensionality of *M*_0_ or *M*_1_ come into play in this calculation. We use the original definition of the reversible jump algorithm from [22], to show why. The ratio of forward and reverse acceptance probabilities for a reversible jump move between states of dimensionalities *N*_0_ and *N*_1_ when the probabilities of proposing the forward and reverse moves are equal should be:

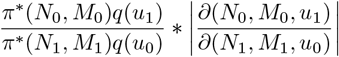

Where *q*(*u*_*i*_) is the probability of generating some vector of variates *u*_*i*_ that are used to generate the dimensionality proposal.

From the perspective of the *β*_0_ chain, 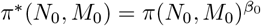 while the ratio 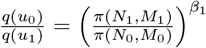 and the magnitude of the Jacobian determinant of the reversible jump move is 1, fulfilling the reversible jump equation from the perspective of the *β*_0_ chain. This logic holds analogously for the *β*_1_ chain.

### 5.10 Data Mixture

To obtain the artificial mixture of the TrpR-ArgR binding data, we took the binary logarithm data from the first ArgR-arg ChIP-chip replicate from [14]. We then took their TrpR-trp ChIP-chip binary logarithm data from the locations of the six main peaks we derived by running macs3 on the TrpR, with some buffer on each side of the peak, ultimate resulting in us taking the windows 1319.75-1322.25 kBp, 1785-1788 kBp, 1983.5-1986.5 kBp, 3302-3305.5 kBp, 3883-3891kBp, and 4629.5-4631.5 kBp from the TrpR data. We then replaced the ArgR data at this locations with the TrpR data multiplied by 4: we performed this multiplication as ArgR binds many more places than TrpR, and we wanted to test our inference with “equal” binding from two transcription factors.

## 6 Acknowledgements

This work was partly funded by the National Institute of General Medical Sciences, National Institutes of Health grant R35 GM128637 awarded to L.F. This work was also partly supported by the National Institute of Health Training Grant NIH 5T32GM070449-14 awarded to A.M.F. TARJIM used the EXPANSE supercomputing cluster at University of California San Diego through allocation MCB160101 from the Advanced Cyberinfrastructure Coordination Ecosystem: Services and Support (ACCESS) program, which is supported by U.S. National Science Foundation grants #2138259, #2138286, #2138307, #2137603, and #2138296. TARJIM also used the Delta advanced computing and data resource which is supported by the National Science Foundation (award OAC 2005572) and the State of Illinois. Delta is a joint effort of the University of Illinois Urbana-Champaign and its National Center for Supercomputing Applications.

## 7 Supplemental Material

- Supplementary File 1: A text file indicating the assignments of the TARJIM inference of IPOD-HR minimal media data set from Freddolino et al. [19] to transcription factors based on similarity in reported occupancy in *E. coli*
- Supplementary File 2: A text file indicating the assignments of the TARJIM inference of IPOD-HR rich defined media data set from Freddolino et al. [19] to transcription factors based on similarity in reported occupancy in *E. coli*
- Supplementary File 3: A folder of the TOMTOM assignments for the Lrp ChIP-chip inference of the TARJIM inference.

## References

[1] Muhammad Idrees Ahmad, Colin David Sinclair, and Barrie D Spurr. Assessment of flood frequency models using empirical distribution function statistics. Water Resources Research, 24(8):1323–1328, 1988.

[2] Kolmogorov An. Sulla determinazione empirica di una legge didistribuzione. Giorn Dell’inst Ital Degli Att, 4:89–91, 1933.

[3] Theodore W Anderson and Donald A Darling. Asymptotic theory of certain” goodness of fit” criteria based on stochastic processes. The annals of mathematical statistics, pages 193–212, 1952.

[4] Gel Mobility Shift Assay. Characterization of cis-regulatory elements and transcription factor binding. Cardiac Gene Expression: Methods and Protocols, 366:183, 2008.

[5] Timothy L. Bailey, James Johnson, Charles E. Grant, and William S. Noble. The MEME Suite. Nucleic Acids Research, 43(W1):W39–W49, 05 2015.

[6] Artem Barski, Suresh Cuddapah, Kairong Cui, Tae-Young Roh, Dustin E Schones, Zhibin Wang, Gang Wei, Iouri Chepelev, and Keji Zhao. High-resolution profiling of histone methylations in the human genome. Cell, 129(4):823–837, 2007.

[7] Leo A Baumgart, Ji Eun Lee, Asaf Salamov, David J Dilworth, Hyunsoo Na, Matthew Mingay, Matthew J Blow, Yu Zhang, Yuko Yoshinaga, Chris G Daum, et al. Persistence and plasticity in bacterial gene regulation. Nature methods, 18(12):1499–1505, 2021.

[8] Cédric Bertrand, Masao Ohmi, Ryoji Suzuki, and Hisashi Kado. A probabilistic solution to the meg inverse problem via mcmc methods: the reversible jump and parallel tempering algorithms. IEEE transactions on biomedical engineering, 48(5):533–542, 2001.

[9] Alan P Boyle, Sean Davis, Hennady P Shulha, Paul Meltzer, Elliott H Margulies, Zhiping Weng, Terrence S Furey, and Gregory E Crawford. High-resolution mapping and characterization of open chromatin across the genome. Cell, 132(2):311–322, 2008.

[10] Douglas F. Browning, Matej Butala, and Stephen J.W. Busby. Bacterial transcription factors: Regulation by pick “n” mix. Journal of Molecular Biology, 431(20):4067–4077, 2019. RNA polymerase reaches 60.

[11] Jason D Buenrostro, Paul G Giresi, Lisa C Zaba, Howard Y Chang, and William J Greenleaf. Transposition of native chromatin for fast and sensitive epigenomic profiling of open chromatin, dna-binding proteins and nucleosome position. Nature methods, 10(12):1213–1218, 2013.

[12] Marina Caldara, Daniel Charlier, and Raymond Cunin. The arginine regulon of escherichia coli: whole-system transcriptome analysis discovers new genes and provides an integrated view of arginine regulation. Microbiology, 152(11):3343–3354, 2006.

[13] Tsu-Pei Chiu, Federico Comoglio, Tianyin Zhou, Lin Yang, Renato Paro, and Remo Rohs. Dnashaper: an r/bioconductor package for dna shape prediction and feature encoding. Bioinformatics, 32(8):1211–1213, 2016.

[14] Byung-Kwan Cho, Stephen Federowicz, Young-Seoub Park, Karsten Zengler, and Bernhard ø Palsson. Deciphering the transcriptional regulatory logic of amino acid metabolism. Nature chemical biology, 8(1):65–71, 2012.

[15] Harald Cramér. On the composition of elementary errors: First paper: Mathematical deductions. Scandinavian Actuarial Journal, 1928(1):13–74, 1928.

[16] Alex de Mendoza and Arnau Sebé-Pedrós. Origin and evolution of eukaryotic transcription factors. Current Opinion in Genetics & Development, 58:25–32, 2019.

[17] Jérôme Déjardin and Robert E Kingston. Purification of proteins associated with specific genomic loci. Cell, 136(1):175–186, 2009.

[18] David J Earl and Michael W Deem. Parallel tempering: Theory, applications, and new perspectives. Physical Chemistry Chemical Physics, 7(23):3910–3916, 2005.

[19] Lydia Freddolino, Haley M Amemiya, Thomas J Goss, and Saeed Tavazoie. Dynamic landscape of protein occupancy across the escherichia coli chromosome. PLoS Biology, 19(6):e3001306, 2021.

[20] Lydia Freddolino, Sasan Amini, and Saeed Tavazoie. Newly identified genetic variations in common escherichia coli mg1655 stock cultures. Journal of bacteriology, 194(2):303–306, 2012.

[21] Charles E Grant, Timothy L Bailey, and William Stafford Noble. Fimo: scanning for occurrences of a given motif. Bioinformatics, 27(7):1017–1018, 2011.

[22] Peter J Green. Reversible jump markov chain monte carlo computation and bayesian model determination. Biometrika, 82(4):711–732, 1995.

[23] Shobhit Gupta, John A Stamatoyannopoulos, Timothy L Bailey, and William Stafford Noble. Quantifying similarity between motifs. Genome biology, 8(2):R24, 2007.

[24] W Keith Hastings. Monte carlo sampling methods using markov chains and their applications. 1970.

[25] Francois Jacob and Jacques Monod. Genetic regulatory mechanisms in the synthesis of proteins. Journal of molecular biology, 3(3):318–356, 1961.

[26] David S Johnson, Ali Mortazavi, Richard M Myers, and Barbara Wold. Genome-wide mapping of in vivo protein-dna interactions. Science, 316(5830):1497–1502, 2007.

[27] Arttu Jolma, Jian Yan, Thomas Whitington, Jarkko Toivonen, Kazuhiro R Nitta, Pasi Rastas, Ekaterina Morgunova, Martin Enge, Mikko Taipale, Gonghong Wei, et al. Dna-binding specificities of human transcription factors. Cell, 152(1):327–339, 2013.

[28] Grace M Kroner, Michael B Wolfe, and Lydia Freddolino. Escherichia coli lrp regulates one-third of the genome via direct, cooperative, and indirect routes. Journal of bacteriology, 201(3):10–1128, 2019.

[29] David J Lee, Stephen D Minchin, and Stephen JW Busby. Activating transcription in bacteria. Annual review of microbiology, 66:125–152, 2012.

[30] Philip Machanick and Timothy L Bailey. Meme-chip: motif analysis of large dna datasets. Bioinformatics, 27(12):1696–1697, 2011.

[31] Sebastian J Maerkl and Stephen R Quake. A systems approach to measuring the binding energy landscapes of transcription factors. Science, 315(5809):233–237, 2007.

[32] Michael P Meers, Karen Adelman, Robert J Duronio, Brian D Strahl, Daniel J McKay, and A Gregory Matera. Transcription start site profiling uncovers divergent transcription and enhancer-associated rnas in drosophila melanogaster. BMC genomics, 19:1–20, 2018.

[33] Sneha Mitra, Anushua Biswas, and Leelavati Narlikar. Diversity in binding, regulation, and evolution revealed from high-throughput chip. PLoS Computational Biology, 14(4):e1006090, 2018.

[34] Lisa R Moore, Ron Caspi, Dana Boyd, Mehmet Berkmen, Amanda Mackie, Suzanne Paley, and Peter D Karp. Revisiting the y-ome of escherichia coli. Nucleic Acids Research, 52(20):12201–12207, 2024.

[35] Mikhail Pachkov, Piotr J Balwierz, Phil Arnold, Evgeniy Ozonov, and Erik Van Nimwegen. Swissregulon, a database of genome-wide annotations of regulatory sites: recent updates. Nucleic acids research, 41(D1):D214–D220, 2012.

[36] Anusri Pampari, Anna Shcherbina, Evgeny Z Kvon, Michael Kosicki, Surag Nair, Soumya Kundu, Arwa S Kathiria, Viviana I Risca, Kristiina Kuningas, Kaur Alasoo, et al. Chrombpnet: bias factorized, base-resolution deep learning models of chromatin accessibility reveal cis-regulatory sequence syntax, transcription factor footprints and regulatory variants. BioRxiv, pages 2024–12, 2025.

[37] G Patikoglou and SK Burley. Eukaryotic transcription factor-dna complexes. Annual review of biophysics and biomolecular structure, 26(1):289–325, 1997.

[38] Roger Pique-Regi, Jacob F Degner, Athma A Pai, Daniel J Gaffney, Yoav Gilad, and Jonathan K Pritchard. Accurate inference of transcription factor binding from dna sequence and chromatin accessibility data. Genome research, 21(3):447–455, 2011.

[39] Ye Ren, Zhouquan Huang, Hao Jiang, Zhuo Wang, Fengsheng Wu, Yufei Xiong, and Jialing Yao. A heat stress responsive nac transcription factor heterodimer plays key roles in rice grain filling. Journal of Experimental Botany, 72(8):2947–2964, 2021.

[40] Heladia Salgado, Socorro Gama-Castro, Paloma Lara, Citlalli Mejia-Almonte, Gabriel Alarcón-Carranza, Andrés G López-Almazo, Felipe Betancourt-Figueroa, Pablo Penña-Loredo, Shirley Alquicira-Hernández, Daniela Ledezma-Tejeida, et al. Regulondb v12. 0: a comprehensive resource of transcriptional regulation in e. coli k-12. Nucleic Acids Research, 52(D1):D255–D264, 2024.

[41] Jeremy W Schroeder, Michael B Wolfe, and Lydia Freddolino. Information theory empowers de novo discovery of structural motifs underpinning protein-dna interactions. Biophysical Journal, 121(3):481a, 2022.

[42] Jeremy W Schroeder, Michael B Wolfe, and Lydia Freddolino. Shapeme: A tool and web front-end for de novo discovery of structural motifs underpinning protein-dna interactions. bioRxiv, 2025.

[43] CD Sinclair, BD Spurr, and MI Ahmad. Modified anderson darling test. Communications in Statistics-Theory and Methods, 19(10):3677–3686, 1990.

[44] Colin J Stirling, G Szatmari, G Stewart, MC Smith, and David J Sherratt. The arginine repressor is essential for plasmid-stabilizing site-specific recombination at the cole1 cer locus. The EMBO journal, 7(13):4389–4395, 1988.

[45] Julian Trouillon, Peter F Doubleday, and Uwe Sauer. Genomic footprinting uncovers global transcription factor responses to amino acids in escherichia coli. Cell Systems, 14(10):860–871, 2023.

[46] Tiffany Vora, Alison K Hottes, and Saeed Tavazoie. Protein occupancy landscape of a bacterial genome. Molecular cell, 35(2):247–253, 2009.

[47] Amy S Weinmann, Stephanie M Bartley, Theresa Zhang, Michael Q Zhang, and Peggy J Farnham. Use of chromatin immunoprecipitation to clone novel e2f target promoters. Molecular and cellular biology, 21(20):6820–6832, 2001.

[48] Nicolae Radu Zabet and Boris Adryan. Estimating binding properties of transcription factors from genome-wide binding profiles. Nucleic Acids Research, 43(1):84–94, 11 2014.

[49] Yong Zhang, Tao Liu, Clifford A Meyer, Jér ôme Eeckhoute, David S Johnson, Bradley E Bernstein, Chad Nusbaum, Richard M Myers, Myles Brown, Wei Li, et al. Model-based analysis of chip-seq (macs). Genome biology, 9(9):R137, 2008.

[50] Christine A Ziegler and Lydia Freddolino. The leucine-responsive regulatory proteins/feast-famine regulatory proteins: an ancient and complex class of transcriptional regulators in bacteria and archaea. Critical reviews in biochemistry and molecular biology, 56(4):373–400, 2021.

[51] Christine A Ziegler and Lydia Freddolino. Escherichia coli leucine-responsive regulatory protein bridges dna in vivo and tunably dissociates in the presence of exogenous leucine. MBio, 14(2):e02690–22, 2023.

[52] Vladimir Mikhailovich Zolotarev. Concerning a certain probability problem. Theory of Probability & Its Applications, 6(2):201–204, 1961.

