## Supplementary File 1 for "The Algorithm for Reversible Jump Inference of Motifs"

Finished outputting binding
Finished regulated loci
Assigned TFs
 

Motif 0 (AATATATAAAT) with peak height 2.2899804723583124 is most likely MIN_EPOD, with log2 fold enrichment 2.186e0 (266 site(s) vs an expected 58.472 site(s)) and p_value 1.622e-114

Factor    	log2 enrichment	p_value 	number of hits	expected number of hits in null distribution
MIN_EPOD  	2.186e0       	1.622e-114	266            	58.472
RDM_EPOD  	2.052e0       	2.244e-100	257            	61.996
Nac       	3.242e0       	9.597e-10	12             	1.269
CRP       	3.158e0       	7.408e-9	11             	1.233
GlaR      	3.519e0       	5.441e-8	8              	0.698
ArcA      	2.825e0       	2.869e-5	6              	0.847
IHF       	2.814e0       	9.566e-5	5              	0.711
FNR       	2.743e0       	1.247e-4	5              	0.747
LeuO      	2.738e0       	3.866e-4	4              	0.599
CpxR      	2.433e0       	9.927e-4	4              	0.741
ArgR      	2.432e0       	9.958e-4	4              	0.741
Fis       	2.132e0       	2.451e-3	4              	0.912
Fur       	2.132e0       	2.456e-3	4              	0.913
SlyA      	2.339e0       	3.194e-3	3              	0.593
LsrR      	1.805e0       	2.039e-2	2              	0.572
YjjQ      	1.795e0       	2.078e-2	2              	0.577
RcsB      	1.790e0       	2.097e-2	2              	0.579
YiaU      	1.783e0       	2.123e-2	2              	0.581
FliZ      	1.764e0       	2.193e-2	2              	0.589
IscR      	1.739e0       	2.293e-2	2              	0.599
NagC      	1.702e0       	2.452e-2	2              	0.615
NsrR      	1.701e0       	2.455e-2	2              	0.615
NarL      	1.675e0       	2.571e-2	2              	0.626
PhoB      	1.673e0       	2.581e-2	2              	0.627
CysB      	1.645e0       	2.710e-2	2              	0.639
OmpR      	1.630e0       	2.780e-2	2              	0.646
Cra       	1.615e0       	2.859e-2	2              	0.653
Lrp       	1.366e0       	4.387e-2	2              	0.776


Motif 1 (TAATATATAAT) with peak height 2.351792369051162 is most likely MIN_EPOD, with log2 fold enrichment 2.255e0 (267 site(s) vs an expected 55.927 site(s)) and p_value 7.045e-122

Factor    	log2 enrichment	p_value 	number of hits	expected number of hits in null distribution
MIN_EPOD  	2.255e0       	7.045e-122	267            	55.927
RDM_EPOD  	2.132e0       	4.354e-109	260            	59.300
Nac       	3.655e0       	2.042e-13	15             	1.191
CRP       	3.250e0       	3.665e-9	11             	1.156
ArcA      	3.515e0       	1.135e-8	9              	0.787
Fur       	3.234e0       	2.802e-7	8              	0.850
IHF       	3.413e0       	4.583e-7	7              	0.657
Lrp       	3.061e0       	1.018e-5	6              	0.719
FNR       	2.532e0       	7.326e-4	4              	0.692
Fis       	2.235e0       	1.807e-3	4              	0.850
NarL      	2.380e0       	2.882e-3	3              	0.576
OmpR      	2.334e0       	3.229e-3	3              	0.595
H-NS      	1.730e0       	2.330e-2	2              	0.603


Motif 2 (TATTAATAAT) with peak height 2.1934417989616932 is most likely MIN_EPOD, with log2 fold enrichment 2.047e0 (203 site(s) vs an expected 49.116 site(s)) and p_value 4.619e-78

Factor    	log2 enrichment	p_value 	number of hits	expected number of hits in null distribution
MIN_EPOD  	2.047e0       	4.619e-78	203            	49.116
RDM_EPOD  	1.984e0       	4.057e-76	206            	52.083
Nac       	3.233e0       	6.822e-8	9              	0.957
CRP       	2.432e0       	3.901e-4	5              	0.927
ArcA      	2.732e0       	3.927e-4	4              	0.602
Fis       	2.606e0       	5.823e-4	4              	0.657
H-NS      	2.770e0       	1.086e-3	3              	0.440
GlaR      	2.654e0       	1.457e-3	3              	0.477
Fur       	2.190e0       	4.582e-3	3              	0.657
RutR      	2.423e0       	6.513e-3	2              	0.373
SlyA      	2.365e0       	7.270e-3	2              	0.388
ArgR      	1.963e0       	1.531e-2	2              	0.513
Lrp       	1.883e0       	1.770e-2	2              	0.542


Motif 3 (TAATAAAATTA) with peak height 2.2424634489610398 is most likely MIN_EPOD, with log2 fold enrichment 2.107e0 (219 site(s) vs an expected 50.844 site(s)) and p_value 1.542e-88

Factor    	log2 enrichment	p_value 	number of hits	expected number of hits in null distribution
MIN_EPOD  	2.107e0       	1.542e-88	219            	50.844
RDM_EPOD  	1.969e0       	7.746e-77	211            	53.913
CRP       	2.793e0       	1.045e-5	7              	1.010
Nac       	2.527e0       	1.029e-4	6              	1.041
H-NS      	2.982e0       	1.788e-4	4              	0.506
Fis       	2.452e0       	9.353e-4	4              	0.731
FNR       	2.354e0       	3.076e-3	3              	0.587
Lrp       	2.293e0       	3.568e-3	3              	0.612
Fur       	2.036e0       	6.631e-3	3              	0.731
SlyA      	2.142e0       	1.102e-2	2              	0.453
LeuO      	2.125e0       	1.138e-2	2              	0.459
NarL      	2.053e0       	1.300e-2	2              	0.482
Cra       	1.985e0       	1.473e-2	2              	0.505
IHF       	1.847e0       	1.889e-2	2              	0.556
CpxR      	1.782e0       	2.124e-2	2              	0.582
ArcA      	1.569e0       	3.093e-2	2              	0.674


Motif 4 (ATATTTATTAA) with peak height 2.1580141101793746 is most likely MIN_EPOD, with log2 fold enrichment 2.054e0 (189 site(s) vs an expected 45.528 site(s)) and p_value 2.821e-73

Factor    	log2 enrichment	p_value 	number of hits	expected number of hits in null distribution
MIN_EPOD  	2.054e0       	2.821e-73	189            	45.528
RDM_EPOD  	1.882e0       	3.581e-60	178            	48.279
ArcA      	3.825e0       	8.872e-9	8              	0.564
FNR       	3.847e0       	4.733e-8	7              	0.486
Nac       	3.162e0       	4.159e-7	8              	0.894
NarL      	3.350e0       	5.475e-5	4              	0.392
CysB      	3.312e0       	6.175e-5	4              	0.403
Fis       	2.700e0       	4.339e-4	4              	0.616
SlyA      	3.034e0       	5.537e-4	3              	0.366
H-NS      	2.857e0       	8.707e-4	3              	0.414
CRP       	2.208e0       	1.948e-3	4              	0.865
GadX      	2.317e0       	7.956e-3	2              	0.401
OmpR      	2.294e0       	8.298e-3	2              	0.408
Fur       	1.699e0       	2.460e-2	2              	0.616
YefM      	1.524e0       	4.801e-2	1              	0.348
YefMB     	1.524e0       	4.801e-2	1              	0.348
FucR      	1.524e0       	4.805e-2	1              	0.348
GlrR      	1.498e0       	4.958e-2	1              	0.354


Motif 5 (ATATAATGATT) with peak height 2.2099437441340872 is most likely MIN_EPOD, with log2 fold enrichment 1.966e0 (152 site(s) vs an expected 38.896 site(s)) and p_value 5.336e-55

Factor    	log2 enrichment	p_value 	number of hits	expected number of hits in null distribution
MIN_EPOD  	1.966e0       	5.336e-55	152            	38.896
RDM_EPOD  	1.843e0       	1.338e-48	148            	41.249
Nac       	3.470e0       	6.986e-8	8              	0.722
Fur       	3.631e0       	7.692e-7	6              	0.484
IHF       	3.101e0       	4.649e-4	3              	0.350
SlyA      	2.884e0       	2.690e-3	2              	0.271
LeuO      	2.861e0       	2.810e-3	2              	0.275
MarA      	2.771e0       	3.345e-3	2              	0.293
GlaR      	2.552e0       	5.095e-3	2              	0.341
CpxR      	2.436e0       	6.345e-3	2              	0.369
Lrp       	2.348e0       	7.502e-3	2              	0.393
ArcA      	2.183e0       	1.020e-2	2              	0.440
PgrR      	1.977e0       	2.723e-2	1              	0.254
AllS      	1.977e0       	2.724e-2	1              	0.254
IlvY      	1.963e0       	2.771e-2	1              	0.256
TorR      	1.940e0       	2.855e-2	1              	0.261
SrsR      	1.914e0       	2.950e-2	1              	0.265
GadW      	1.852e0       	3.190e-2	1              	0.277
CRP       	1.519e0       	3.371e-2	2              	0.698
PhoP      	1.777e0       	3.507e-2	1              	0.292
GadX      	1.732e0       	3.707e-2	1              	0.301
CysB      	1.727e0       	3.731e-2	1              	0.302
OxyR      	1.721e0       	3.760e-2	1              	0.303
SoxS      	1.719e0       	3.768e-2	1              	0.304
OmpR      	1.707e0       	3.828e-2	1              	0.306
LexA      	1.677e0       	3.976e-2	1              	0.313


Motif 6 (TATTACATATA) with peak height 2.418901415992615 is most likely MIN_EPOD, with log2 fold enrichment 2.115e0 (157 site(s) vs an expected 36.248 site(s)) and p_value 1.214e-64

Factor    	log2 enrichment	p_value 	number of hits	expected number of hits in null distribution
MIN_EPOD  	2.115e0       	1.214e-64	157            	36.248
RDM_EPOD  	1.974e0       	5.641e-56	151            	38.441
Nac       	3.606e0       	3.154e-8	8              	0.657
ArcA      	3.926e0       	1.973e-7	6              	0.395
CRP       	2.977e0       	5.087e-5	5              	0.635
Fur       	3.198e0       	8.881e-5	4              	0.436
ArgR      	3.189e0       	3.696e-4	3              	0.329
NagC      	2.998e0       	2.154e-3	2              	0.250
NsrR      	2.997e0       	2.159e-3	2              	0.251
OmpR      	2.890e0       	2.656e-3	2              	0.270
H-NS      	2.863e0       	2.799e-3	2              	0.275
IHF       	2.688e0       	3.921e-3	2              	0.310
CpxR      	2.605e0       	4.599e-3	2              	0.329
FNR       	2.588e0       	4.752e-3	2              	0.333
Fis       	2.199e0       	9.900e-3	2              	0.436
CsqR      	2.163e0       	2.146e-2	1              	0.223
RcsAB     	2.148e0       	2.189e-2	1              	0.226
DpiA      	2.146e0       	2.193e-2	1              	0.226
AcrR      	2.124e0       	2.257e-2	1              	0.229
MntR      	2.122e0       	2.263e-2	1              	0.230
BtsR      	2.121e0       	2.264e-2	1              	0.230
DcuR      	2.116e0       	2.278e-2	1              	0.231
YdeO      	2.110e0       	2.298e-2	1              	0.232
CsgD      	2.059e0       	2.452e-2	1              	0.240
LeuO      	2.054e0       	2.468e-2	1              	0.241
ZraR      	2.052e0       	2.474e-2	1              	0.241
TyrR      	2.051e0       	2.476e-2	1              	0.241
PhoP      	1.964e0       	2.767e-2	1              	0.256
NarL      	1.957e0       	2.793e-2	1              	0.258
GadX      	1.917e0       	2.937e-2	1              	0.265
Cra       	1.867e0       	3.132e-2	1              	0.274
GlaR      	1.727e0       	3.734e-2	1              	0.302


Motif 7 (ATTTAATTATTT) with peak height 2.3287570225478396 is most likely RDM_EPOD, with log2 fold enrichment 1.933e0 (143 site(s) vs an expected 37.439 site(s)) and p_value 3.233e-51

Factor    	log2 enrichment	p_value 	number of hits	expected number of hits in null distribution
RDM_EPOD  	1.933e0       	3.233e-51	143            	37.439
MIN_EPOD  	1.977e0       	5.189e-51	139            	35.304
IHF       	3.987e0       	9.943e-7	5              	0.315
Nac       	3.200e0       	5.365e-6	6              	0.653
Lrp       	3.496e0       	3.374e-5	4              	0.354
BtsR      	3.077e0       	1.842e-3	2              	0.237
GadX      	2.884e0       	2.687e-3	2              	0.271
CRP       	2.249e0       	3.958e-3	3              	0.631
CpxR      	2.586e0       	4.773e-3	2              	0.333
Fis       	2.194e0       	9.999e-3	2              	0.437
Fur       	2.193e0       	1.001e-2	2              	0.437
BluR      	2.121e0       	2.266e-2	1              	0.230
MarR      	2.119e0       	2.270e-2	1              	0.230
MurR      	2.114e0       	2.287e-2	1              	0.231
RcdA      	2.100e0       	2.326e-2	1              	0.233
NhaR      	2.091e0       	2.352e-2	1              	0.235
Ada       	2.090e0       	2.356e-2	1              	0.235
LrhA      	2.060e0       	2.450e-2	1              	0.240
GalS      	2.059e0       	2.451e-2	1              	0.240
GalR      	2.059e0       	2.451e-2	1              	0.240
RcsB-BglJ 	2.054e0       	2.468e-2	1              	0.241
BasR      	2.054e0       	2.468e-2	1              	0.241
LeuO      	2.014e0       	2.597e-2	1              	0.248
FadR      	2.012e0       	2.602e-2	1              	0.248
GlpR      	2.004e0       	2.629e-2	1              	0.249
NagC      	1.960e0       	2.780e-2	1              	0.257
MarA      	1.923e0       	2.915e-2	1              	0.264
OmpR      	1.858e0       	3.165e-2	1              	0.276
Cra       	1.836e0       	3.256e-2	1              	0.280


Motif 8 (TATTAATCTAA) with peak height 2.302875332837443 is most likely MIN_EPOD, with log2 fold enrichment 1.908e0 (136 site(s) vs an expected 36.248 site(s)) and p_value 6.710e-47

Factor    	log2 enrichment	p_value 	number of hits	expected number of hits in null distribution
MIN_EPOD  	1.908e0       	6.710e-47	136            	36.248
RDM_EPOD  	1.780e0       	5.126e-41	132            	38.441
Nac       	3.606e0       	3.154e-8	8              	0.657
ArcA      	3.926e0       	1.973e-7	6              	0.395
Fis       	3.784e0       	3.794e-7	6              	0.436
IHF       	3.688e0       	1.799e-5	4              	0.310
FNR       	3.588e0       	2.500e-5	4              	0.333
CRP       	2.977e0       	5.087e-5	5              	0.635
Fur       	3.198e0       	8.881e-5	4              	0.436
Lrp       	3.097e0       	4.684e-4	3              	0.351
BtsR      	3.121e0       	1.691e-3	2              	0.230
DcuR      	3.116e0       	1.707e-3	2              	0.231
H-NS      	2.863e0       	2.799e-3	2              	0.275
XapR      	2.168e0       	2.133e-2	1              	0.223
RcnR      	2.165e0       	2.140e-2	1              	0.223
DpiA      	2.146e0       	2.193e-2	1              	0.226
SgrR      	2.141e0       	2.206e-2	1              	0.227
DeoR      	2.139e0       	2.213e-2	1              	0.227
NhaR      	2.136e0       	2.222e-2	1              	0.227
ModE      	2.116e0       	2.280e-2	1              	0.231
ExuR      	2.105e0       	2.311e-2	1              	0.232
BasR      	2.096e0       	2.337e-2	1              	0.234
SlyA      	2.078e0       	2.393e-2	1              	0.237
TyrR      	2.051e0       	2.476e-2	1              	0.241
ArgP      	2.037e0       	2.524e-2	1              	0.244
MetJ      	2.020e0       	2.579e-2	1              	0.247
NarL      	1.957e0       	2.793e-2	1              	0.258
CysB      	1.912e0       	2.957e-2	1              	0.266
OxyR      	1.905e0       	2.982e-2	1              	0.267
OmpR      	1.890e0       	3.040e-2	1              	0.270
LexA      	1.858e0       	3.166e-2	1              	0.276
CpxR      	1.605e0       	4.343e-2	1              	0.329
ArgR      	1.604e0       	4.351e-2	1              	0.329


Motif 9 (AATATATTAT) with peak height 2.1976365423003705 is most likely MIN_EPOD, with log2 fold enrichment 2.196e0 (166 site(s) vs an expected 36.228 site(s)) and p_value 4.719e-73

Factor    	log2 enrichment	p_value 	number of hits	expected number of hits in null distribution
MIN_EPOD  	2.196e0       	4.719e-73	166            	36.228
RDM_EPOD  	2.103e0       	2.385e-68	165            	38.421
Nac       	4.557e0       	1.345e-17	15             	0.637
GlaR      	3.824e0       	1.149e-5	4              	0.282
Lrp       	3.181e0       	3.771e-4	3              	0.331
Fur       	2.850e0       	8.829e-4	3              	0.416
McbR      	3.299e0       	1.191e-3	2              	0.203
RcsB-BglJ 	3.224e0       	1.382e-3	2              	0.214
LeuO      	3.178e0       	1.514e-3	2              	0.221
PhoP      	3.080e0       	1.833e-3	2              	0.237
OmpR      	3.000e0       	2.144e-3	2              	0.250
Cra       	2.975e0       	2.253e-3	2              	0.254
H-NS      	2.971e0       	2.270e-3	2              	0.255
IHF       	2.783e0       	3.265e-3	2              	0.290
CRP       	2.286e0       	3.614e-3	3              	0.615
ArcA      	2.415e0       	6.602e-3	2              	0.375
CusR      	2.298e0       	1.803e-2	1              	0.203
FrmR      	2.297e0       	1.806e-2	1              	0.204
HprR      	2.295e0       	1.809e-2	1              	0.204
UxuR      	2.256e0       	1.902e-2	1              	0.209
ExuR      	2.234e0       	1.959e-2	1              	0.213
SlyA      	2.204e0       	2.036e-2	1              	0.217
PurR      	2.172e0       	2.122e-2	1              	0.222
GadW      	2.167e0       	2.135e-2	1              	0.223
NagC      	2.116e0       	2.278e-2	1              	0.231
MarA      	2.074e0       	2.406e-2	1              	0.238
GadX      	2.029e0       	2.547e-2	1              	0.245
CysB      	2.023e0       	2.566e-2	1              	0.246
CpxR      	1.695e0       	3.885e-2	1              	0.309
ArgR      	1.693e0       	3.893e-2	1              	0.309
FNR       	1.677e0       	3.974e-2	1              	0.313


Motif 10 (AATATTATAA) with peak height 2.3341527616466973 is most likely MIN_EPOD, with log2 fold enrichment 2.206e0 (161 site(s) vs an expected 34.906 site(s)) and p_value 1.692e-71

Factor    	log2 enrichment	p_value 	number of hits	expected number of hits in null distribution
MIN_EPOD  	2.206e0       	1.692e-71	161            	34.906
RDM_EPOD  	2.130e0       	9.219e-69	162            	37.019
Fis       	3.667e0       	3.535e-6	5              	0.394
GadX      	4.125e0       	4.221e-6	4              	0.229
CRP       	3.093e0       	3.267e-5	5              	0.586
ArcA      	3.496e0       	3.372e-5	4              	0.354
GadW      	3.853e0       	6.446e-5	3              	0.208
GlrR      	3.375e0       	1.025e-3	2              	0.193
GalR      	3.333e0       	1.114e-3	2              	0.198
GalS      	3.333e0       	1.114e-3	2              	0.198
SlyA      	3.306e0       	1.175e-3	2              	0.202
IHF       	2.873e0       	2.745e-3	2              	0.273
Nac       	2.304e0       	3.456e-3	3              	0.607
Lrp       	2.681e0       	3.975e-3	2              	0.312
HigA      	2.412e0       	1.554e-2	1              	0.188
GadRcs    	2.407e0       	1.566e-2	1              	0.189
HigBA     	2.404e0       	1.572e-2	1              	0.189
BaeR      	2.388e0       	1.603e-2	1              	0.191
Zur       	2.383e0       	1.615e-2	1              	0.192
TorR      	2.373e0       	1.635e-2	1              	0.193
UxuR      	2.360e0       	1.663e-2	1              	0.195
EvgA      	2.360e0       	1.663e-2	1              	0.195
AcrR      	2.357e0       	1.669e-2	1              	0.195
TrpR      	2.344e0       	1.698e-2	1              	0.197
YdeO      	2.342e0       	1.704e-2	1              	0.197
RcsB-BglJ 	2.327e0       	1.737e-2	1              	0.199
IscR      	2.280e0       	1.844e-2	1              	0.206
NtrC      	2.266e0       	1.879e-2	1              	0.208
CytR      	2.242e0       	1.938e-2	1              	0.211
MarA      	2.171e0       	2.123e-2	1              	0.222
PhoB      	2.166e0       	2.137e-2	1              	0.223
CysB      	2.120e0       	2.269e-2	1              	0.230
SoxS      	2.110e0       	2.297e-2	1              	0.232
OmpR      	2.095e0       	2.341e-2	1              	0.234
Cra       	2.069e0       	2.420e-2	1              	0.238
H-NS      	2.065e0       	2.432e-2	1              	0.239
FNR       	1.763e0       	3.565e-2	1              	0.295


Motif 11 (ATAATATAATT) with peak height 2.3827856316294076 is most likely MIN_EPOD, with log2 fold enrichment 2.178e0 (170 site(s) vs an expected 37.572 site(s)) and p_value 1.256e-73

Factor    	log2 enrichment	p_value 	number of hits	expected number of hits in null distribution
MIN_EPOD  	2.178e0       	1.256e-73	170            	37.572
RDM_EPOD  	2.024e0       	1.262e-62	162            	39.844
ArcA      	4.261e0       	6.537e-10	8              	0.417
IHF       	4.408e0       	2.381e-9	7              	0.330
Nac       	3.537e0       	4.724e-8	8              	0.689
CRP       	3.171e0       	6.115e-6	6              	0.666
H-NS      	3.771e0       	1.373e-5	4              	0.293
LexA      	3.351e0       	2.426e-4	3              	0.294
CpxR      	3.105e0       	4.600e-4	3              	0.349
FNR       	3.088e0       	4.802e-4	3              	0.353
Fur       	2.706e0       	1.274e-3	3              	0.460
PgrR      	3.076e0       	1.850e-3	2              	0.237
RcsAB     	3.047e0       	1.958e-3	2              	0.242
DcuR      	3.016e0       	2.077e-3	2              	0.247
LeuO      	2.956e0       	2.337e-3	2              	0.258
NsrR      	2.900e0       	2.604e-3	2              	0.268
NarL      	2.862e0       	2.806e-3	2              	0.275
Fis       	2.122e0       	1.143e-2	2              	0.460
GadRcs    	2.065e0       	2.433e-2	1              	0.239
MprA      	2.056e0       	2.461e-2	1              	0.240
PspF      	2.055e0       	2.466e-2	1              	0.241
BaeR      	2.050e0       	2.481e-2	1              	0.242
DpiA      	2.045e0       	2.496e-2	1              	0.242
Zur       	2.045e0       	2.496e-2	1              	0.242
SoxR      	2.039e0       	2.514e-2	1              	0.243
XylR      	2.027e0       	2.553e-2	1              	0.245
BtsR      	2.021e0       	2.574e-2	1              	0.246
Mlc       	2.019e0       	2.582e-2	1              	0.247
MlrA      	2.014e0       	2.596e-2	1              	0.248
SrsR      	2.010e0       	2.609e-2	1              	0.248
SlyA      	1.979e0       	2.715e-2	1              	0.254
AraC      	1.972e0       	2.740e-2	1              	0.255
Rob       	1.953e0       	2.806e-2	1              	0.258
GadW      	1.947e0       	2.830e-2	1              	0.259
NagC      	1.901e0       	2.997e-2	1              	0.268
MarA      	1.863e0       	3.145e-2	1              	0.275
GadX      	1.823e0       	3.307e-2	1              	0.283
OxyR      	1.812e0       	3.356e-2	1              	0.285
SoxS      	1.810e0       	3.364e-2	1              	0.285
OmpR      	1.797e0       	3.419e-2	1              	0.288
Cra       	1.774e0       	3.518e-2	1              	0.292
GlaR      	1.638e0       	4.171e-2	1              	0.321
ArgR      	1.518e0       	4.837e-2	1              	0.349


Motif 12 (ATTTTATAAAT) with peak height 2.272122816989032 is most likely MIN_EPOD, with log2 fold enrichment 2.103e0 (148 site(s) vs an expected 34.443 site(s)) and p_value 1.742e-60

Factor    	log2 enrichment	p_value 	number of hits	expected number of hits in null distribution
MIN_EPOD  	2.103e0       	1.742e-60	148            	34.443
RDM_EPOD  	1.928e0       	1.337e-49	139            	36.527
ArcA      	4.039e0       	1.165e-7	6              	0.365
CRP       	3.338e0       	2.875e-6	6              	0.593
Fur       	3.630e0       	4.086e-6	5              	0.404
FNR       	3.709e0       	1.679e-5	4              	0.306
Nac       	3.025e0       	4.245e-5	5              	0.614
SlyA      	3.803e0       	7.345e-5	3              	0.215
NarL      	3.677e0       	1.028e-4	3              	0.235
CysB      	3.630e0       	1.164e-4	3              	0.242
LexA      	3.574e0       	1.349e-4	3              	0.252
GlaR      	3.437e0       	1.932e-4	3              	0.277
IHF       	3.398e0       	2.143e-4	3              	0.285
YiaU      	3.265e0       	1.273e-3	2              	0.208
NarP      	3.207e0       	1.427e-3	2              	0.217
NsrR      	3.133e0       	1.651e-3	2              	0.228
Fis       	2.309e0       	8.067e-3	2              	0.404
FucR      	2.316e0       	1.761e-2	1              	0.201
UhpA      	2.312e0       	1.771e-2	1              	0.201
PlaR      	2.293e0       	1.814e-2	1              	0.204
YjjQ      	2.285e0       	1.832e-2	1              	0.205
Mlc       	2.261e0       	1.891e-2	1              	0.209
SrsR      	2.252e0       	1.913e-2	1              	0.210
RcsB-BglJ 	2.238e0       	1.949e-2	1              	0.212
LeuO      	2.193e0       	2.064e-2	1              	0.219
FadR      	2.192e0       	2.068e-2	1              	0.219
GadW      	2.183e0       	2.091e-2	1              	0.220
GlpR      	2.183e0       	2.092e-2	1              	0.220
CytR      	2.159e0       	2.156e-2	1              	0.224
NagC      	2.134e0       	2.226e-2	1              	0.228
MarA      	2.093e0       	2.347e-2	1              	0.234
GadX      	2.050e0       	2.479e-2	1              	0.241
Cra       	1.998e0       	2.652e-2	1              	0.250


Motif 13 (TGTATATATAA) with peak height 2.3182072875167714 is most likely MIN_EPOD, with log2 fold enrichment 2.198e0 (152 site(s) vs an expected 33.120 site(s)) and p_value 2.984e-67

Factor    	log2 enrichment	p_value 	number of hits	expected number of hits in null distribution
MIN_EPOD  	2.198e0       	2.984e-67	152            	33.120
RDM_EPOD  	2.045e0       	1.965e-57	145            	35.125
Nac       	4.362e0       	6.581e-14	12             	0.584
LexA      	4.896e0       	1.708e-10	7              	0.235
Fis       	3.977e0       	1.552e-7	6              	0.381
IHF       	3.907e0       	8.703e-6	4              	0.267
ArgR      	3.818e0       	1.172e-5	4              	0.284
CysB      	3.731e0       	8.887e-5	3              	0.226
Lrp       	3.306e0       	2.723e-4	3              	0.303
CRP       	2.828e0       	2.882e-4	4              	0.563
Fur       	2.976e0       	6.394e-4	3              	0.381
Cbl       	3.404e0       	9.662e-4	2              	0.189
RcsB-BglJ 	3.346e0       	1.085e-3	2              	0.197
LeuO      	3.300e0       	1.189e-3	2              	0.203
TyrR      	3.297e0       	1.196e-3	2              	0.203
NarL      	3.195e0       	1.462e-3	2              	0.218
GadX      	3.152e0       	1.590e-3	2              	0.225
Cra       	3.098e0       	1.770e-3	2              	0.234
FNR       	2.801e0       	3.154e-3	2              	0.287
ArcA      	2.540e0       	5.202e-3	2              	0.344
GadRcs    	2.423e0       	1.533e-2	1              	0.186
IlvY      	2.418e0       	1.542e-2	1              	0.187
MazE      	2.406e0       	1.566e-2	1              	0.189
MazEF     	2.404e0       	1.570e-2	1              	0.189
GcvA      	2.395e0       	1.589e-2	1              	0.190
TorR      	2.391e0       	1.598e-2	1              	0.191
RcsB      	2.387e0       	1.606e-2	1              	0.191
BtsR      	2.373e0       	1.636e-2	1              	0.193
GadE      	2.354e0       	1.676e-2	1              	0.196
BasR      	2.346e0       	1.694e-2	1              	0.197
NarP      	2.314e0       	1.765e-2	1              	0.201
GadW      	2.289e0       	1.823e-2	1              	0.205
ArgP      	2.281e0       	1.843e-2	1              	0.206
NagC      	2.239e0       	1.946e-2	1              	0.212
NsrR      	2.238e0       	1.949e-2	1              	0.212
CpxR      	1.819e0       	3.324e-2	1              	0.283


Motif 14 (TAATTAAATATT) with peak height 2.402706825691282 is most likely MIN_EPOD, with log2 fold enrichment 2.053e0 (127 site(s) vs an expected 30.611 site(s)) and p_value 4.803e-50

Factor    	log2 enrichment	p_value 	number of hits	expected number of hits in null distribution
MIN_EPOD  	2.053e0       	4.803e-50	127            	30.611
RDM_EPOD  	1.990e0       	6.347e-49	129            	32.464
Nac       	4.057e0       	3.083e-10	9              	0.541
CRP       	3.745e0       	7.813e-8	7              	0.522
ArcA      	3.648e0       	2.045e-5	4              	0.319
PhoP      	3.892e0       	5.797e-5	3              	0.202
Lrp       	3.412e0       	2.057e-4	3              	0.282
DcuR      	3.470e0       	8.473e-4	2              	0.180
SlyA      	3.429e0       	9.197e-4	2              	0.186
IscR      	3.405e0       	9.650e-4	2              	0.189
GadW      	3.393e0       	9.886e-4	2              	0.190
GadX      	3.256e0       	1.294e-3	2              	0.209
Cra       	3.202e0       	1.440e-3	2              	0.217
GlaR      	3.053e0       	1.928e-3	2              	0.241
ArgR      	2.924e0       	2.484e-3	2              	0.264
FNR       	2.907e0       	2.565e-3	2              	0.267
Fis       	2.500e0       	5.615e-3	2              	0.354
HcaR      	2.532e0       	1.330e-2	1              	0.173
YefMB     	2.531e0       	1.331e-2	1              	0.173
YefM      	2.531e0       	1.331e-2	1              	0.173
Dan       	2.529e0       	1.335e-2	1              	0.173
MngR      	2.514e0       	1.361e-2	1              	0.175
CaiF      	2.508e0       	1.371e-2	1              	0.176
RcsAB     	2.504e0       	1.378e-2	1              	0.176
PuuR      	2.499e0       	1.387e-2	1              	0.177
TorR      	2.493e0       	1.398e-2	1              	0.178
LeuO      	2.403e0       	1.572e-2	1              	0.189
CytR      	2.368e0       	1.646e-2	1              	0.194
NagC      	2.342e0       	1.701e-2	1              	0.197
MarA      	2.300e0       	1.797e-2	1              	0.203
NarL      	2.299e0       	1.800e-2	1              	0.203
OxyR      	2.243e0       	1.934e-2	1              	0.211
OmpR      	2.227e0       	1.974e-2	1              	0.214
IHF       	2.013e0       	2.599e-2	1              	0.248
CpxR      	1.925e0       	2.906e-2	1              	0.263


Motif 15 (ATAACTAGA) with peak height 2.2921351387658366 is most likely RDM_EPOD, with log2 fold enrichment 1.648e0 (96 site(s) vs an expected 30.642 site(s)) and p_value 1.017e-26

Factor    	log2 enrichment	p_value 	number of hits	expected number of hits in null distribution
RDM_EPOD  	1.648e0       	1.017e-26	96             	30.642
MIN_EPOD  	1.671e0       	5.806e-26	92             	28.890
Nac       	3.106e0       	1.188e-4	4              	0.465
ArcA      	3.556e0       	1.408e-4	3              	0.255
Fur       	3.382e0       	2.227e-4	3              	0.288
NfeR      	4.081e0       	2.487e-4	2              	0.118
LeuO      	3.920e0       	3.443e-4	2              	0.132
CysB      	3.718e0       	5.170e-4	2              	0.152
H-NS      	3.650e0       	5.919e-4	2              	0.159
GlaR      	3.465e0       	8.557e-4	2              	0.181
IHF       	3.414e0       	9.462e-4	2              	0.188
CRP       	2.747e0       	1.142e-3	3              	0.447
PgrR      	3.104e0       	6.240e-3	1              	0.116
StpA      	3.103e0       	6.246e-3	1              	0.116
AllR      	3.081e0       	6.430e-3	1              	0.118
MqsA      	3.056e0       	6.650e-3	1              	0.120
Mlc       	3.015e0       	7.020e-3	1              	0.124
GntR      	2.994e0       	7.226e-3	1              	0.126
ArgP      	2.895e0       	8.241e-3	1              	0.134
NagC      	2.839e0       	8.874e-3	1              	0.140
PhoB      	2.776e0       	9.640e-3	1              	0.146
Cra       	2.655e0       	1.131e-2	1              	0.159
CpxR      	2.306e0       	1.784e-2	1              	0.202
FNR       	2.283e0       	1.836e-2	1              	0.205
Fis       	1.798e0       	3.414e-2	1              	0.288


Motif 16 (TAATTAAAAATAA) with peak height 2.226853696927035 is most likely MIN_EPOD, with log2 fold enrichment 2.270e0 (146 site(s) vs an expected 30.264 site(s)) and p_value 1.080e-68

Factor    	log2 enrichment	p_value 	number of hits	expected number of hits in null distribution
MIN_EPOD  	2.270e0       	1.080e-68	146            	30.264
RDM_EPOD  	2.156e0       	2.069e-62	143            	32.095
IHF       	4.284e0       	3.019e-7	5              	0.257
Nac       	3.457e0       	1.668e-6	6              	0.546
ArcA      	3.197e0       	3.609e-4	3              	0.327
GntR      	3.382e0       	1.009e-3	2              	0.192
LeuO      	3.332e0       	1.115e-3	2              	0.199
PhoP      	3.241e0       	1.335e-3	2              	0.212
NarL      	3.233e0       	1.354e-3	2              	0.213
CysB      	3.188e0       	1.481e-3	2              	0.219
CpxR      	2.878e0       	2.714e-3	2              	0.272
FNR       	2.861e0       	2.806e-3	2              	0.275
Lrp       	2.784e0       	3.254e-3	2              	0.290
StpA      	2.456e0       	1.467e-2	1              	0.182
CitR      	2.455e0       	1.469e-2	1              	0.182
FucR      	2.450e0       	1.479e-2	1              	0.183
ZntR      	2.444e0       	1.490e-2	1              	0.184
IdnR      	2.443e0       	1.493e-2	1              	0.184
NorR      	2.442e0       	1.495e-2	1              	0.184
AsnC      	2.434e0       	1.509e-2	1              	0.185
TorR      	2.416e0       	1.546e-2	1              	0.187
FliZ      	2.372e0       	1.636e-2	1              	0.193
CRP       	1.922e0       	1.645e-2	2              	0.528
NarP      	2.345e0       	1.696e-2	1              	0.197
FadR      	2.330e0       	1.729e-2	1              	0.199
TyrR      	2.329e0       	1.732e-2	1              	0.199
GadW      	2.322e0       	1.748e-2	1              	0.200
NagC      	2.274e0       	1.858e-2	1              	0.207
MarA      	2.235e0       	1.956e-2	1              	0.212
PhoB      	2.230e0       	1.968e-2	1              	0.213
GadX      	2.193e0       	2.064e-2	1              	0.219
H-NS      	2.138e0       	2.214e-2	1              	0.227
GlaR      	2.001e0       	2.641e-2	1              	0.250
ArgR      	1.877e0       	3.091e-2	1              	0.272


Motif 17 (TAAATATAACA) with peak height 2.506090904627015 is most likely MIN_EPOD, with log2 fold enrichment 2.255e0 (130 site(s) vs an expected 27.235 site(s)) and p_value 1.137e-60

Factor    	log2 enrichment	p_value 	number of hits	expected number of hits in null distribution
MIN_EPOD  	2.255e0       	1.137e-60	130            	27.235
RDM_EPOD  	2.043e0       	1.802e-47	119            	28.885
OmpR      	5.207e0       	4.712e-10	6              	0.162
CRP       	4.001e0       	2.010e-8	7              	0.437
Nac       	3.947e0       	2.681e-8	7              	0.454
ArcA      	4.285e0       	2.990e-7	5              	0.257
Fur       	4.121e0       	5.766e-7	5              	0.287
PhoP      	4.300e0       	1.939e-5	3              	0.152
RcsB-BglJ 	3.884e0       	3.697e-4	2              	0.135
LeuO      	3.830e0       	4.128e-4	2              	0.141
GadX      	3.656e0       	5.849e-4	2              	0.159
IHF       	3.374e0       	1.025e-3	2              	0.193
CpxR      	3.274e0       	1.249e-3	2              	0.207
FrlR      	2.993e0       	7.231e-3	1              	0.126
NfeR      	2.971e0       	7.444e-3	1              	0.128
NadR      	2.970e0       	7.460e-3	1              	0.128
MngR      	2.963e0       	7.526e-3	1              	0.128
RstA      	2.955e0       	7.604e-3	1              	0.129
MqsA      	2.949e0       	7.665e-3	1              	0.129
YjjQ      	2.944e0       	7.721e-3	1              	0.130
XylR      	2.925e0       	7.912e-3	1              	0.132
BtsR      	2.917e0       	8.002e-3	1              	0.132
GadW      	2.817e0       	9.132e-3	1              	0.142
NsrR      	2.756e0       	9.900e-3	1              	0.148
MarA      	2.707e0       	1.055e-2	1              	0.153
PhoB      	2.701e0       	1.064e-2	1              	0.154
OxyR      	2.640e0       	1.152e-2	1              	0.160
H-NS      	2.588e0       	1.235e-2	1              	0.166
FNR       	2.253e0       	1.909e-2	1              	0.210
Lrp       	2.163e0       	2.144e-2	1              	0.223
Fis       	1.800e0       	3.404e-2	1              	0.287


Motif 18 (CTAAAATAAATA) with peak height 2.3853548518191667 is most likely MIN_EPOD, with log2 fold enrichment 2.270e0 (121 site(s) vs an expected 25.085 site(s)) and p_value 2.305e-57

Factor    	log2 enrichment	p_value 	number of hits	expected number of hits in null distribution
MIN_EPOD  	2.270e0       	2.305e-57	121            	25.085
RDM_EPOD  	2.149e0       	1.178e-51	118            	26.605
CRP       	3.895e0       	2.211e-7	6              	0.403
Nac       	3.578e0       	4.919e-6	5              	0.419
Fur       	3.500e0       	1.629e-4	3              	0.265
RcsB      	4.046e0       	2.669e-4	2              	0.121
FNR       	3.368e0       	1.036e-3	2              	0.194
ArcA      	3.078e0       	1.833e-3	2              	0.237
Fis       	2.916e0       	2.518e-3	2              	0.265
RcsAB     	3.063e0       	6.582e-3	1              	0.120
NhaR      	3.048e0       	6.717e-3	1              	0.121
UxuR      	3.035e0       	6.833e-3	1              	0.122
MntR      	3.029e0       	6.887e-3	1              	0.122
BtsR      	3.029e0       	6.892e-3	1              	0.123
Mlc       	3.026e0       	6.921e-3	1              	0.123
ExuR      	3.008e0       	7.083e-3	1              	0.124
SlyA      	2.973e0       	7.423e-3	1              	0.127
CsgD      	2.949e0       	7.668e-3	1              	0.130
FadR      	2.940e0       	7.755e-3	1              	0.130
NagC      	2.870e0       	8.509e-3	1              	0.137
NsrR      	2.869e0       	8.525e-3	1              	0.137
GadX      	2.769e0       	9.729e-3	1              	0.147
OmpR      	2.735e0       	1.017e-2	1              	0.150
Cra       	2.706e0       	1.057e-2	1              	0.153
H-NS      	2.701e0       	1.063e-2	1              	0.154
GlaR      	2.534e0       	1.324e-2	1              	0.173
IHF       	2.488e0       	1.407e-2	1              	0.178
CpxR      	2.388e0       	1.602e-2	1              	0.191


Motif 19 (ATTATTATATAC) with peak height 2.3831272228221128 is most likely MIN_EPOD, with log2 fold enrichment 2.315e0 (138 site(s) vs an expected 27.727 site(s)) and p_value 1.708e-67

Factor    	log2 enrichment	p_value 	number of hits	expected number of hits in null distribution
MIN_EPOD  	2.315e0       	1.708e-67	138            	27.727
RDM_EPOD  	2.188e0       	2.761e-60	134            	29.406
Fur       	3.707e0       	1.676e-5	4              	0.306
Nac       	3.072e0       	1.322e-4	4              	0.476
GlaR      	3.294e0       	1.201e-3	2              	0.204
CitR      	2.817e0       	9.138e-3	1              	0.142
QseB      	2.780e0       	9.585e-3	1              	0.146
DpiA      	2.780e0       	9.585e-3	1              	0.146
YiaU      	2.753e0       	9.939e-3	1              	0.148
DcuR      	2.745e0       	1.004e-2	1              	0.149
BasR      	2.722e0       	1.036e-2	1              	0.152
SlyA      	2.700e0       	1.066e-2	1              	0.154
TyrR      	2.668e0       	1.111e-2	1              	0.157
LexA      	2.444e0       	1.491e-2	1              	0.184
ArgR      	2.155e0       	2.166e-2	1              	0.224
ArcA      	1.864e0       	3.141e-2	1              	0.275
Fis       	1.708e0       	3.818e-2	1              	0.306


Motif 20 (ATTAAATATGCA) with peak height 2.2835483276231643 is most likely MIN_EPOD, with log2 fold enrichment 2.044e0 (97 site(s) vs an expected 23.526 site(s)) and p_value 1.603e-38

Factor    	log2 enrichment	p_value 	number of hits	expected number of hits in null distribution
MIN_EPOD  	2.044e0       	1.603e-38	97             	23.526
RDM_EPOD  	1.867e0       	1.642e-31	91             	24.952
ArgR      	5.340e0       	1.472e-11	7              	0.173
NsrR      	5.037e0       	1.923e-7	4              	0.122
CRP       	3.750e0       	2.489e-6	5              	0.372
LeuO      	4.700e0       	6.557e-6	3              	0.115
Fis       	3.632e0       	1.148e-4	3              	0.242
PhoB      	3.980e0       	3.043e-4	2              	0.127
Cra       	3.866e0       	3.829e-4	2              	0.137
H-NS      	3.861e0       	3.864e-4	2              	0.138
CpxR      	3.535e0       	7.429e-4	2              	0.173
CadC      	3.266e0       	5.023e-3	1              	0.104
IclR      	3.255e0       	5.095e-3	1              	0.105
Cbl       	3.247e0       	5.151e-3	1              	0.105
MntR      	3.207e0       	5.431e-3	1              	0.108
LrhA      	3.181e0       	5.625e-3	1              	0.110
RcsB-BglJ 	3.173e0       	5.688e-3	1              	0.111
BasR      	3.173e0       	5.688e-3	1              	0.111
SlyA      	3.147e0       	5.883e-3	1              	0.113
NarP      	3.133e0       	5.998e-3	1              	0.114
TyrR      	3.111e0       	6.176e-3	1              	0.116
GlpR      	3.101e0       	6.259e-3	1              	0.117
CytR      	3.071e0       	6.516e-3	1              	0.119
PhoP      	2.994e0       	7.215e-3	1              	0.126
NarL      	2.985e0       	7.309e-3	1              	0.126
CysB      	2.925e0       	7.909e-3	1              	0.132
IHF       	2.638e0       	1.155e-2	1              	0.161
Lrp       	2.421e0       	1.536e-2	1              	0.187
Fur       	2.046e0       	2.491e-2	1              	0.242


Motif 21 (ATTTATTATAAT) with peak height 2.2446292484265977 is most likely MIN_EPOD, with log2 fold enrichment 2.248e0 (114 site(s) vs an expected 24.006 site(s)) and p_value 3.262e-53

Factor    	log2 enrichment	p_value 	number of hits	expected number of hits in null distribution
MIN_EPOD  	2.248e0       	3.262e-53	114            	24.006
RDM_EPOD  	2.124e0       	1.008e-47	111            	25.460
Nac       	3.921e0       	1.956e-7	6              	0.396
Fur       	4.005e0       	6.225e-6	4              	0.249
ArgR      	4.072e0       	3.560e-5	3              	0.178
CysB      	3.874e0       	3.769e-4	2              	0.136
LexA      	3.805e0       	4.332e-4	2              	0.143
Dan       	3.218e0       	5.355e-3	1              	0.107
EvgA      	3.158e0       	5.805e-3	1              	0.112
MlrA      	3.141e0       	5.931e-3	1              	0.113
DnaA      	3.122e0       	6.085e-3	1              	0.115
YfeC      	3.118e0       	6.120e-3	1              	0.115
SlyA      	3.093e0       	6.330e-3	1              	0.117
NarL      	2.932e0       	7.833e-3	1              	0.131
CpxR      	2.489e0       	1.405e-2	1              	0.178
Lrp       	2.376e0       	1.628e-2	1              	0.193
ArcA      	2.172e0       	2.120e-2	1              	0.222


Motif 22 (ACAAAAATAAGAA) with peak height 2.822618233296507 is most likely MIN_EPOD, with log2 fold enrichment 2.323e0 (71 site(s) vs an expected 14.190 site(s)) and p_value 3.146e-36

Factor    	log2 enrichment	p_value 	number of hits	expected number of hits in null distribution
MIN_EPOD  	2.323e0       	3.146e-36	71             	14.190
RDM_EPOD  	2.176e0       	1.516e-31	68             	15.051
Fur       	4.581e0       	8.879e-6	3              	0.125
H-NS      	5.008e0       	3.731e-5	2              	0.062
Lrp       	4.444e0       	1.179e-4	2              	0.092
UhpA      	4.585e0       	8.374e-4	1              	0.042
FliZ      	4.437e0       	1.026e-3	1              	0.046
CsgD      	4.367e0       	1.128e-3	1              	0.048
NsrR      	4.248e0       	1.327e-3	1              	0.053
OxyR      	4.081e0       	1.664e-3	1              	0.059
IHF       	3.717e0       	2.730e-3	1              	0.076
ArgR      	3.583e0       	3.267e-3	1              	0.083
FNR       	3.559e0       	3.376e-3	1              	0.085
ArcA      	3.194e0       	5.509e-3	1              	0.109
Fis       	2.997e0       	7.169e-3	1              	0.125
CRP       	2.297e0       	1.799e-2	1              	0.204


Motif 23 (TTTTATAGTATT) with peak height 2.3496336312364225 is most likely MIN_EPOD, with log2 fold enrichment 2.282e0 (69 site(s) vs an expected 14.187 site(s)) and p_value 3.705e-34

Factor    	log2 enrichment	p_value 	number of hits	expected number of hits in null distribution
MIN_EPOD  	2.282e0       	3.705e-34	69             	14.187
RDM_EPOD  	2.155e0       	1.405e-30	67             	15.048
GlaR      	4.840e0       	5.259e-5	2              	0.070
KdpE      	4.691e0       	7.244e-4	1              	0.039
CusR      	4.688e0       	7.273e-4	1              	0.039
HprR      	4.682e0       	7.330e-4	1              	0.039
PepA      	4.657e0       	7.589e-4	1              	0.040
RcsAB     	4.651e0       	7.648e-4	1              	0.040
DeoR      	4.632e0       	7.853e-4	1              	0.040
EvgA      	4.604e0       	8.162e-4	1              	0.041
YiaU      	4.597e0       	8.242e-4	1              	0.041
ModE      	4.582e0       	8.414e-4	1              	0.042
MarA      	4.259e0       	1.307e-3	1              	0.052
OxyR      	4.158e0       	1.500e-3	1              	0.056
SoxS      	4.155e0       	1.507e-3	1              	0.056
Cra       	4.086e0       	1.653e-3	1              	0.059
ArgR      	3.637e0       	3.039e-3	1              	0.080
FNR       	3.612e0       	3.144e-3	1              	0.082
Lrp       	3.493e0       	3.692e-3	1              	0.089
Fis       	3.033e0       	6.838e-3	1              	0.122
CRP       	2.318e0       	1.749e-2	1              	0.200
Nac       	2.257e0       	1.895e-2	1              	0.209


Motif 24 (AAAATCGTGCAAATTCAA) with peak height 9.607002297643668 is most likely IHF, with log2 fold enrichment 7.942e0 (2 site(s) vs an expected 0.008 site(s)) and p_value 7.690e-8

Factor    	log2 enrichment	p_value 	number of hits	expected number of hits in null distribution
IHF       	7.942e0       	7.690e-8	2              	0.008
NsrR      	7.966e0       	7.599e-6	1              	0.004
Fur       	5.892e0       	1.336e-4	1              	0.017
Nac       	4.958e0       	4.837e-4	1              	0.032


Motif 25 (ATGCTAATTAT) with peak height 2.1361116888182803 is most likely MIN_EPOD, with log2 fold enrichment 1.855e0 (47 site(s) vs an expected 12.989 site(s)) and p_value 5.419e-17

Factor    	log2 enrichment	p_value 	number of hits	expected number of hits in null distribution
MIN_EPOD  	1.855e0       	5.419e-17	47             	12.989
RDM_EPOD  	1.801e0       	1.006e-16	48             	13.778
CRP       	4.073e0       	3.470e-5	3              	0.178
Fur       	4.229e0       	1.820e-4	2              	0.107
StpA      	5.087e0       	4.207e-4	1              	0.029
LeuO      	4.775e0       	6.450e-4	1              	0.037
TyrR      	4.769e0       	6.508e-4	1              	0.037
PhoP      	4.571e0       	8.535e-4	1              	0.042
Nac       	3.424e0       	9.157e-4	2              	0.186
CysB      	4.458e0       	9.950e-4	1              	0.045
H-NS      	4.358e0       	1.142e-3	1              	0.049
FNR       	3.846e0       	2.289e-3	1              	0.070
ArcA      	3.444e0       	3.940e-3	1              	0.092


Motif 26 (ACTTAAAAAATAT) with peak height 2.8228870524478364 is most likely MIN_EPOD, with log2 fold enrichment 2.391e0 (55 site(s) vs an expected 10.486 site(s)) and p_value 3.388e-30

Factor    	log2 enrichment	p_value 	number of hits	expected number of hits in null distribution
MIN_EPOD  	2.391e0       	3.388e-30	55             	10.486
RDM_EPOD  	2.252e0       	9.408e-27	53             	11.123
DpiA      	5.374e0       	2.830e-4	1              	0.024
DcuR      	5.294e0       	3.157e-4	1              	0.025
NtrC      	5.108e0       	4.076e-4	1              	0.029
CRP       	3.807e0       	4.236e-4	2              	0.143
NagC      	5.001e0       	4.723e-4	1              	0.031
NsrR      	4.999e0       	4.740e-4	1              	0.031
OmpR      	4.762e0       	6.555e-4	1              	0.037
Cra       	4.712e0       	7.016e-4	1              	0.038
GlaR      	4.434e0       	1.026e-3	1              	0.046
FNR       	4.182e0       	1.448e-3	1              	0.055
Lrp       	4.052e0       	1.729e-3	1              	0.060
Nac       	2.743e0       	1.001e-2	1              	0.149


Motif 27 (AATTATCTCAATGT) with peak height 2.7970697108046227 is most likely RDM_EPOD, with log2 fold enrichment 2.229e0 (48 site(s) vs an expected 10.238 site(s)) and p_value 5.099e-24

Factor    	log2 enrichment	p_value 	number of hits	expected number of hits in null distribution
RDM_EPOD  	2.229e0       	5.099e-24	48             	10.238
MIN_EPOD  	2.253e0       	4.244e-23	46             	9.652
H-NS      	5.832e0       	6.775e-6	2              	0.035
Fis       	4.679e0       	7.217e-5	2              	0.078
RcsAB     	5.509e0       	2.346e-4	1              	0.022
XylR      	5.455e0       	2.527e-4	1              	0.023
AraC      	5.307e0       	3.102e-4	1              	0.025
Nac       	3.864e0       	3.766e-4	2              	0.137
IHF       	4.488e0       	9.530e-4	1              	0.045
CpxR      	4.337e0       	1.171e-3	1              	0.049
CRP       	2.929e0       	7.824e-3	1              	0.131


Motif 28 (TAAATTGCAATAT) with peak height 2.6898577754008803 is most likely MIN_EPOD, with log2 fold enrichment 1.847e0 (36 site(s) vs an expected 10.008 site(s)) and p_value 1.876e-13

Factor    	log2 enrichment	p_value 	number of hits	expected number of hits in null distribution
MIN_EPOD  	1.847e0       	1.876e-13	36             	10.008
RDM_EPOD  	1.721e0       	6.158e-12	35             	10.616
NsrR      	6.697e0       	2.651e-8	3              	0.029
ArgR      	5.888e0       	2.455e-7	3              	0.051
CRP       	4.469e0       	1.176e-5	3              	0.135
IHF       	5.459e0       	1.461e-5	2              	0.045
Fur       	4.639e0       	7.844e-5	2              	0.080
CaiF      	5.517e0       	2.324e-4	1              	0.022
PlaR      	5.511e0       	2.343e-4	1              	0.022
YdeO      	5.401e0       	2.726e-4	1              	0.024
GadW      	5.229e0       	3.450e-4	1              	0.027
CytR      	5.173e0       	3.730e-4	1              	0.028
Nac       	3.820e0       	4.125e-4	2              	0.142
PhoP      	5.035e0       	4.504e-4	1              	0.030
GadX      	4.928e0       	5.217e-4	1              	0.033


Motif 29 (TTCATCAAATGATA) with peak height 4.315678937619924 is most likely MIN_EPOD, with log2 fold enrichment 2.820e0 (21 site(s) vs an expected 2.975 site(s)) and p_value 7.333e-18

Factor    	log2 enrichment	p_value 	number of hits	expected number of hits in null distribution
MIN_EPOD  	2.820e0       	7.333e-18	21             	2.975
RDM_EPOD  	2.664e0       	1.007e-15	20             	3.156
Cra       	7.252e0       	2.057e-5	1              	0.007
H-NS      	7.240e0       	2.091e-5	1              	0.007
IHF       	6.713e0       	4.337e-5	1              	0.010
Nac       	4.712e0       	6.828e-4	1              	0.038


Motif 30 (TTTTTTATGCGGTT) with peak height 5.375968590548846 is most likely RDM_EPOD, with log2 fold enrichment 2.276e0 (11 site(s) vs an expected 2.272 site(s)) and p_value 1.444e-7

Factor    	log2 enrichment	p_value 	number of hits	expected number of hits in null distribution
RDM_EPOD  	2.276e0       	1.444e-7	11             	2.272
ArgR      	7.040e0       	2.713e-5	1              	0.008
Lrp       	6.816e0       	3.701e-5	1              	0.009
MIN_EPOD  	1.901e0       	8.420e-5	8              	2.141
Fis       	6.166e0       	9.082e-5	1              	0.014
CRP       	5.278e0       	3.087e-4	1              	0.026
Nac       	5.206e0       	3.412e-4	1              	0.027


Motif 31 (TATCACTATTCATT) with peak height 4.435696670023551 is most likely MIN_EPOD, with log2 fold enrichment 2.619e0 (19 site(s) vs an expected 3.094 site(s)) and p_value 3.624e-14

Factor    	log2 enrichment	p_value 	number of hits	expected number of hits in null distribution
MIN_EPOD  	2.619e0       	3.624e-14	19             	3.094
RDM_EPOD  	2.373e0       	3.738e-11	17             	3.282
Nac       	6.237e0       	7.962e-8	3              	0.040
ArgR      	6.430e0       	6.424e-5	1              	0.012


Motif 32 (TAAGCGGAGCGCATCAGG) with peak height 2.5199166521832312 is most likely IHF, with log2 fold enrichment 6.093e0 (2 site(s) vs an expected 0.029 site(s)) and p_value 3.886e-6

Factor    	log2 enrichment	p_value 	number of hits	expected number of hits in null distribution
IHF       	6.093e0       	3.886e-6	2              	0.029
NsrR      	5.775e0       	1.619e-4	1              	0.018


Motif 33 (AACCTGTTGAAACT) with peak height 4.308277898169546 is most likely Lrp, with log2 fold enrichment 6.816e0 (1 site(s) vs an expected 0.009 site(s)) and p_value 3.701e-5

Factor    	log2 enrichment	p_value 	number of hits	expected number of hits in null distribution
Lrp       	6.816e0       	3.701e-5	1              	0.009
Fur       	6.165e0       	9.102e-5	1              	0.014
Nac       	5.206e0       	3.412e-4	1              	0.027
MIN_EPOD  	1.223e0       	1.475e-2	5              	2.141
RDM_EPOD  	1.138e0       	1.940e-2	5              	2.272


Motif 34 (GGTATATGCTTTTT) with peak height 3.8961635166233872 is most likely MIN_EPOD, with log2 fold enrichment 2.486e0 (18 site(s) vs an expected 3.213 site(s)) and p_value 2.317e-12

Factor    	log2 enrichment	p_value 	number of hits	expected number of hits in null distribution
MIN_EPOD  	2.486e0       	2.317e-12	18             	3.213
RDM_EPOD  	2.401e0       	6.687e-12	18             	3.408
Nac       	5.595e0       	1.024e-5	2              	0.041
ArgR      	6.365e0       	7.029e-5	1              	0.012
ArcA      	5.797e0       	1.540e-4	1              	0.018


Motif 35 (TAATATTGAGTA) with peak height 2.8876400637324187 is most likely RDM_EPOD, with log2 fold enrichment 2.438e0 (26 site(s) vs an expected 4.797 site(s)) and p_value 1.562e-16

Factor    	log2 enrichment	p_value 	number of hits	expected number of hits in null distribution
RDM_EPOD  	2.438e0       	1.562e-16	26             	4.797
MIN_EPOD  	2.282e0       	1.382e-12	22             	4.522
CysB      	6.677e0       	4.623e-5	1              	0.010
ArcA      	5.268e0       	3.226e-4	1              	0.026
Fur       	5.007e0       	4.618e-4	1              	0.031
CRP       	4.157e0       	1.477e-3	1              	0.056
Nac       	4.086e0       	1.626e-3	1              	0.059


Motif 36 (AATTCTCAGCTAAT) with peak height 4.883948913884732 is most likely MIN_EPOD, with log2 fold enrichment 2.569e0 (12 site(s) vs an expected 2.022 site(s)) and p_value 1.425e-9

Factor    	log2 enrichment	p_value 	number of hits	expected number of hits in null distribution
MIN_EPOD  	2.569e0       	1.425e-9	12             	2.022
RDM_EPOD  	2.221e0       	7.668e-7	10             	2.145
GlaR      	7.476e0       	1.480e-5	1              	0.006
FNR       	7.093e0       	2.514e-5	1              	0.007


Motif 37 (TATGTTATTGATA) with peak height 3.31111502360704 is most likely MIN_EPOD, with log2 fold enrichment 2.171e0 (15 site(s) vs an expected 3.332 site(s)) and p_value 1.186e-8

Factor    	log2 enrichment	p_value 	number of hits	expected number of hits in null distribution
MIN_EPOD  	2.171e0       	1.186e-8	15             	3.332
RDM_EPOD  	2.085e0       	2.782e-8	15             	3.534
AraC      	7.934e0       	8.045e-6	1              	0.004
CRP       	5.617e0       	9.830e-6	2              	0.041
OxyR      	7.208e0       	2.196e-5	1              	0.007
ArcA      	5.751e0       	1.643e-4	1              	0.019
Nac       	4.545e0       	8.611e-4	1              	0.043


Motif 38 (GATATCAACAAGATA) with peak height 5.484474109992438 is most likely IHF, with log2 fold enrichment 7.668e0 (1 site(s) vs an expected 0.005 site(s)) and p_value 1.119e-5

Factor    	log2 enrichment	p_value 	number of hits	expected number of hits in null distribution
IHF       	7.668e0       	1.119e-5	1              	0.005
FNR       	7.393e0       	1.637e-5	1              	0.006
Lrp       	7.205e0       	2.124e-5	1              	0.007
ArcA      	6.825e0       	3.593e-5	1              	0.009
CRP       	5.650e0       	1.820e-4	1              	0.020
Nac       	5.577e0       	2.013e-4	1              	0.021
MIN_EPOD  	1.586e0       	3.628e-3	5              	1.665
RDM_EPOD  	1.501e0       	4.897e-3	5              	1.767


Motif 39 (CTTATCATGTTGATT) with peak height 5.997214549525336 is most likely GadE, with log2 fold enrichment 9.410e0 (1 site(s) vs an expected 0.001 site(s)) and p_value 1.008e-6

Factor    	log2 enrichment	p_value 	number of hits	expected number of hits in null distribution
GadE      	9.410e0       	1.008e-6	1              	0.001
LrhA      	9.397e0       	1.026e-6	1              	0.001
RDM_EPOD  	2.249e0       	1.675e-6	9              	1.893
LexA      	8.113e0       	6.079e-6	1              	0.004
MIN_EPOD  	2.165e0       	1.213e-5	8              	1.784
CpxR      	7.327e0       	1.803e-5	1              	0.006


Motif 40 (AAATATGGTGATTTT) with peak height 4.901808612748266 is most likely RDM_EPOD, with log2 fold enrichment 2.861e0 (11 site(s) vs an expected 1.514 site(s)) and p_value 1.630e-11

Factor    	log2 enrichment	p_value 	number of hits	expected number of hits in null distribution
RDM_EPOD  	2.861e0       	1.630e-11	11             	1.514
MIN_EPOD  	2.809e0       	7.212e-10	10             	1.427


Motif 41 (TTTGGTAATCCATTG) with peak height 5.548975977235349 is most likely MIN_EPOD, with log2 fold enrichment 2.657e0 (9 site(s) vs an expected 1.427 site(s)) and p_value 2.977e-8

Factor    	log2 enrichment	p_value 	number of hits	expected number of hits in null distribution
MIN_EPOD  	2.657e0       	2.977e-8	9              	1.427
RDM_EPOD  	2.571e0       	5.296e-8	9              	1.514


Motif 42 (AATATAAATTTGCAAA) with peak height 4.145717969334324 is most likely RDM_EPOD, with log2 fold enrichment 2.887e0 (14 site(s) vs an expected 1.893 site(s)) and p_value 3.280e-14

Factor    	log2 enrichment	p_value 	number of hits	expected number of hits in null distribution
RDM_EPOD  	2.887e0       	3.280e-14	14             	1.893
MIN_EPOD  	2.865e0       	1.516e-12	13             	1.784
GadE      	9.363e0       	1.076e-6	1              	0.002
GadW      	8.961e0       	1.877e-6	1              	0.002
GadX      	8.328e0       	4.511e-6	1              	0.003
H-NS      	8.113e0       	6.079e-6	1              	0.004
Lrp       	7.086e0       	2.516e-5	1              	0.007


Motif 43 (CATATAAAATACCAT) with peak height 5.731606152522131 is most likely MIN_EPOD, with log2 fold enrichment 2.612e0 (8 site(s) vs an expected 1.308 site(s)) and p_value 2.090e-7

Factor    	log2 enrichment	p_value 	number of hits	expected number of hits in null distribution
MIN_EPOD  	2.612e0       	2.090e-7	8              	1.308
RDM_EPOD  	2.527e0       	3.506e-7	8              	1.388


Motif 44 (ACAGTTGTAGTTATT) with peak height 4.926674684573577 is most likely MIN_EPOD, with log2 fold enrichment 2.724e0 (11 site(s) vs an expected 1.665 site(s)) and p_value 5.790e-10

Factor    	log2 enrichment	p_value 	number of hits	expected number of hits in null distribution
MIN_EPOD  	2.724e0       	5.790e-10	11             	1.665
RDM_EPOD  	2.638e0       	1.159e-9	11             	1.767
EvgA      	9.746e0       	6.289e-7	1              	0.001
Nac       	5.577e0       	2.013e-4	1              	0.021


Motif 45 (CATAATATTGCGTT) with peak height 5.856915610105799 is most likely MIN_EPOD, with log2 fold enrichment 2.782e0 (9 site(s) vs an expected 1.308 site(s)) and p_value 5.561e-9

Factor    	log2 enrichment	p_value 	number of hits	expected number of hits in null distribution
MIN_EPOD  	2.782e0       	5.561e-9	9              	1.308
Lrp       	7.589e0       	1.222e-5	1              	0.005
RDM_EPOD  	2.112e0       	1.054e-4	6              	1.388


Motif 46 (TAAGTTGTTGATACAT) with peak height 5.697809314919233 is most likely MIN_EPOD, with log2 fold enrichment 2.264e0 (8 site(s) vs an expected 1.665 site(s)) and p_value 5.430e-6

Factor    	log2 enrichment	p_value 	number of hits	expected number of hits in null distribution
MIN_EPOD  	2.264e0       	5.430e-6	8              	1.665
RDM_EPOD  	1.986e0       	9.490e-5	7              	1.767


Motif 47 (ACCCAATAATTGAAA) with peak height 4.574691164945593 is most likely MIN_EPOD, with log2 fold enrichment 2.934e0 (10 site(s) vs an expected 1.308 site(s)) and p_value 6.743e-11

Factor    	log2 enrichment	p_value 	number of hits	expected number of hits in null distribution
MIN_EPOD  	2.934e0       	6.743e-11	10             	1.308
RDM_EPOD  	2.849e0       	1.291e-10	10             	1.388
ArgR      	7.816e0       	8.924e-6	1              	0.004


Motif 48 (ATACTGGTGTTAAAT) with peak height 5.288567007614533 is most likely RDM_EPOD, with log2 fold enrichment 2.834e0 (9 site(s) vs an expected 1.262 site(s)) and p_value 1.023e-9

Factor    	log2 enrichment	p_value 	number of hits	expected number of hits in null distribution
RDM_EPOD  	2.834e0       	1.023e-9	9              	1.262
MIN_EPOD  	2.750e0       	4.255e-8	8              	1.189
Nac       	6.075e0       	9.823e-5	1              	0.015


Motif 49 (AAATAATAAGCTCTA) with peak height 4.447379946439174 is most likely MIN_EPOD, with log2 fold enrichment 2.541e0 (9 site(s) vs an expected 1.546 site(s)) and p_value 1.151e-7

Factor    	log2 enrichment	p_value 	number of hits	expected number of hits in null distribution
MIN_EPOD  	2.541e0       	1.151e-7	9              	1.546
RDM_EPOD  	2.456e0       	2.033e-7	9              	1.640


Motif 50 (TTTTATGATGATATT) with peak height 5.137527929306289 is most likely MIN_EPOD, with log2 fold enrichment 2.487e0 (6 site(s) vs an expected 1.070 site(s)) and p_value 9.733e-6

Factor    	log2 enrichment	p_value 	number of hits	expected number of hits in null distribution
MIN_EPOD  	2.487e0       	9.733e-6	6              	1.070
RDM_EPOD  	2.402e0       	1.451e-5	6              	1.136


Motif 51 (ATCTTATTTCCTGA) with peak height 5.31077977424697 is most likely MIN_EPOD, with log2 fold enrichment 2.487e0 (8 site(s) vs an expected 1.427 site(s)) and p_value 7.469e-7

Factor    	log2 enrichment	p_value 	number of hits	expected number of hits in null distribution
MIN_EPOD  	2.487e0       	7.469e-7	8              	1.427
RDM_EPOD  	2.401e0       	1.243e-6	8              	1.514
Nac       	5.808e0       	1.445e-4	1              	0.018


Motif 52 (AGTATTTTTACATAA) with peak height 6.26385150336751 is most likely RDM_EPOD, with log2 fold enrichment 2.817e0 (8 site(s) vs an expected 1.136 site(s)) and p_value 8.107e-9

Factor    	log2 enrichment	p_value 	number of hits	expected number of hits in null distribution
RDM_EPOD  	2.817e0       	8.107e-9	8              	1.136
MIN_EPOD  	2.709e0       	3.224e-7	7              	1.070


Motif 53 (TCAGATGGAATTC) with peak height 5.0776629528232675 is most likely MIN_EPOD, with log2 fold enrichment 2.434e0 (9 site(s) vs an expected 1.665 site(s)) and p_value 3.594e-7

Factor    	log2 enrichment	p_value 	number of hits	expected number of hits in null distribution
MIN_EPOD  	2.434e0       	3.594e-7	9              	1.665
RDM_EPOD  	2.349e0       	6.300e-7	9              	1.767
FNR       	7.414e0       	1.591e-5	1              	0.006
Fur       	6.555e0       	5.216e-5	1              	0.011


Motif 54 (TTTCAAACATTA) with peak height 3.0059366582555644 is most likely MIN_EPOD, with log2 fold enrichment 2.419e0 (21 site(s) vs an expected 3.927 site(s)) and p_value 2.355e-13

Factor    	log2 enrichment	p_value 	number of hits	expected number of hits in null distribution
MIN_EPOD  	2.419e0       	2.355e-13	21             	3.927
RDM_EPOD  	2.263e0       	1.016e-11	20             	4.166
YdeO      	8.844e0       	1.247e-8	2              	0.004
GadW      	8.500e0       	2.550e-8	2              	0.006
GadX      	7.974e0       	7.603e-8	2              	0.008
CadC      	8.189e0       	5.681e-6	1              	0.003
ArgP      	7.463e0       	1.551e-5	1              	0.006
H-NS      	6.787e0       	3.955e-5	1              	0.009
GlaR      	6.377e0       	6.963e-5	1              	0.012
FNR       	6.026e0       	1.130e-4	1              	0.015
ArcA      	5.499e0       	2.338e-4	1              	0.022
Fis       	5.235e0       	3.362e-4	1              	0.027


Motif 55 (GAGTAATATTGTTT) with peak height 3.597045819077894 is most likely RDM_EPOD, with log2 fold enrichment 2.308e0 (15 site(s) vs an expected 3.029 site(s)) and p_value 1.112e-9

Factor    	log2 enrichment	p_value 	number of hits	expected number of hits in null distribution
RDM_EPOD  	2.308e0       	1.112e-9	15             	3.029
MIN_EPOD  	2.187e0       	6.911e-8	13             	2.856
GadW      	8.075e0       	6.571e-6	1              	0.004
GadX      	7.513e0       	1.432e-5	1              	0.005
Fis       	5.718e0       	1.710e-4	1              	0.019
Fur       	5.716e0       	1.714e-4	1              	0.019


Motif 56 (TATTGATCATCTTG) with peak height 6.2578006137548705 is most likely MIN_EPOD, with log2 fold enrichment 2.657e0 (9 site(s) vs an expected 1.427 site(s)) and p_value 2.977e-8

Factor    	log2 enrichment	p_value 	number of hits	expected number of hits in null distribution
MIN_EPOD  	2.657e0       	2.977e-8	9              	1.427
RDM_EPOD  	2.571e0       	5.295e-8	9              	1.514


Motif 57 (ATAGGGACATTGA) with peak height 5.45727563847912 is most likely MIN_EPOD, with log2 fold enrichment 3.072e0 (8 site(s) vs an expected 0.951 site(s)) and p_value 0.000e0

Factor    	log2 enrichment	p_value 	number of hits	expected number of hits in null distribution
MIN_EPOD  	3.072e0       	0.000e0 	8              	0.951
RDM_EPOD  	2.987e0       	0.000e0 	8              	1.009
EvgA      	1.102e1       	1.019e-7	1              	0.000
LeuO      	1.030e1       	2.749e-7	1              	0.001


Motif 58 (GTACTTTCACAAAT) with peak height 3.2928237677253254 is most likely RDM_EPOD, with log2 fold enrichment 2.472e0 (14 site(s) vs an expected 2.524 site(s)) and p_value 2.714e-10

Factor    	log2 enrichment	p_value 	number of hits	expected number of hits in null distribution
RDM_EPOD  	2.472e0       	2.714e-10	14             	2.524
MIN_EPOD  	2.450e0       	2.179e-9	13             	2.379


Motif 59 (TGAAAATGATGTGATTA) with peak height 4.4805140239457675 is most likely NarP, with log2 fold enrichment 1.008e1 (1 site(s) vs an expected 0.001 site(s)) and p_value 3.768e-7

Factor    	log2 enrichment	p_value 	number of hits	expected number of hits in null distribution
NarP      	1.008e1       	3.768e-7	1              	0.001
NarL      	9.395e0       	9.797e-7	1              	0.001
FNR       	8.073e0       	6.116e-6	1              	0.004
MIN_EPOD  	2.224e0       	1.724e-4	5              	1.070
RDM_EPOD  	1.817e0       	2.591e-3	4              	1.136


Motif 60 (TTGATTTCATGATTT) with peak height 4.504134400225303 is most likely MIN_EPOD, with log2 fold enrichment 2.946e0 (11 site(s) vs an expected 1.427 site(s)) and p_value 8.024e-12

Factor    	log2 enrichment	p_value 	number of hits	expected number of hits in null distribution
MIN_EPOD  	2.946e0       	8.024e-12	11             	1.427
RDM_EPOD  	2.723e0       	1.371e-9	10             	1.514
GlaR      	8.034e0       	6.660e-6	1              	0.004
Nac       	5.806e0       	1.450e-4	1              	0.018


Motif 61 (TGTCAGAAAGATATCGC) with peak height 6.139433881230946 is most likely MIN_EPOD, with log2 fold enrichment 3.072e0 (7 site(s) vs an expected 0.833 site(s)) and p_value 0.000e0

Factor    	log2 enrichment	p_value 	number of hits	expected number of hits in null distribution
MIN_EPOD  	3.072e0       	0.000e0 	7              	0.833
RDM_EPOD  	2.987e0       	0.000e0 	7              	0.883


Motif 62 (TTATTTACGTTATA) with peak height 3.532782051953324 is most likely MIN_EPOD, with log2 fold enrichment 2.901e0 (16 site(s) vs an expected 2.141 site(s)) and p_value 3.060e-15

Factor    	log2 enrichment	p_value 	number of hits	expected number of hits in null distribution
MIN_EPOD  	2.901e0       	3.060e-15	16             	2.141
RDM_EPOD  	2.816e0       	8.286e-15	16             	2.272
PdeL      	9.588e0       	7.963e-7	1              	0.001
GadW      	8.670e0       	2.841e-6	1              	0.002
ArgP      	8.625e0       	3.024e-6	1              	0.003
GadX      	8.047e0       	6.736e-6	1              	0.004


Motif 63 (ATTCAGTCAATTACAA) with peak height 5.545985820946878 is most likely RDM_EPOD, with log2 fold enrichment 2.501e0 (5 site(s) vs an expected 0.883 site(s)) and p_value 2.519e-5

Factor    	log2 enrichment	p_value 	number of hits	expected number of hits in null distribution
RDM_EPOD  	2.501e0       	2.519e-5	5              	0.883
MIN_EPOD  	2.264e0       	4.058e-4	4              	0.833


Motif 64 (ATAATTAAGGTTTTTA) with peak height 5.168152426160848 is most likely MIN_EPOD, with log2 fold enrichment 2.879e0 (7 site(s) vs an expected 0.952 site(s)) and p_value 4.005e-8

Factor    	log2 enrichment	p_value 	number of hits	expected number of hits in null distribution
MIN_EPOD  	2.879e0       	4.005e-8	7              	0.952
RDM_EPOD  	2.794e0       	6.424e-8	7              	1.009
Nac       	6.402e0       	6.085e-5	1              	0.012


Motif 65 (AAGGCATATGCTAA) with peak height 5.572051724931704 is most likely MIN_EPOD, with log2 fold enrichment 2.586e0 (5 site(s) vs an expected 0.833 site(s)) and p_value 1.779e-5

Factor    	log2 enrichment	p_value 	number of hits	expected number of hits in null distribution
MIN_EPOD  	2.586e0       	1.779e-5	5              	0.833
RDM_EPOD  	2.501e0       	2.518e-5	5              	0.883
CRP       	6.675e0       	4.086e-5	1              	0.010


Motif 66 (TCCTTTGTTTTTATT) with peak height 5.5285782000339765 is most likely RDM_EPOD, with log2 fold enrichment 2.308e0 (5 site(s) vs an expected 1.009 site(s)) and p_value 8.989e-5

Factor    	log2 enrichment	p_value 	number of hits	expected number of hits in null distribution
RDM_EPOD  	2.308e0       	8.989e-5	5              	1.009
MIN_EPOD  	2.072e0       	9.755e-4	4              	0.951


Motif 67 (TTATGTTCTTTGCTA) with peak height 4.904568187630259 is most likely MIN_EPOD, with log2 fold enrichment 2.879e0 (7 site(s) vs an expected 0.951 site(s)) and p_value 4.004e-8

Factor    	log2 enrichment	p_value 	number of hits	expected number of hits in null distribution
MIN_EPOD  	2.879e0       	4.004e-8	7              	0.951
RDM_EPOD  	2.794e0       	6.424e-8	7              	1.009


Motif 68 (AAGATATTTAAAATG) with peak height 3.5754828537167995 is most likely MIN_EPOD, with log2 fold enrichment 2.809e0 (10 site(s) vs an expected 1.427 site(s)) and p_value 7.212e-10

Factor    	log2 enrichment	p_value 	number of hits	expected number of hits in null distribution
MIN_EPOD  	2.809e0       	7.212e-10	10             	1.427
RDM_EPOD  	2.723e0       	1.371e-9	10             	1.514
NarP      	9.590e0       	7.714e-7	1              	0.001
NsrR      	9.139e0       	1.440e-6	1              	0.002
PhoP      	8.967e0       	1.828e-6	1              	0.002
NarL      	8.930e0       	1.924e-6	1              	0.002
Fis       	6.780e0       	3.774e-5	1              	0.009


Motif 69 (TTAACCTACTGTATT) with peak height 5.0373242563429175 is most likely BtsR, with log2 fold enrichment 1.042e1 (1 site(s) vs an expected 0.001 site(s)) and p_value 2.385e-7

Factor    	log2 enrichment	p_value 	number of hits	expected number of hits in null distribution
BtsR      	1.042e1       	2.385e-7	1              	0.001
MIN_EPOD  	2.557e0       	1.442e-6	7              	1.189
RDM_EPOD  	2.249e0       	4.306e-5	6              	1.262


Motif 70 (ACTGCATTATCATAT) with peak height 4.516095348025802 is most likely RDM_EPOD, with log2 fold enrichment 2.794e0 (7 site(s) vs an expected 1.009 site(s)) and p_value 6.424e-8

Factor    	log2 enrichment	p_value 	number of hits	expected number of hits in null distribution
RDM_EPOD  	2.794e0       	6.424e-8	7              	1.009
SlyA      	1.045e1       	2.237e-7	1              	0.001
MIN_EPOD  	2.657e0       	2.413e-6	6              	0.951


Motif 71 (TCAGATATTCTACTG) with peak height 5.077236845481441 is most likely MIN_EPOD, with log2 fold enrichment 2.657e0 (6 site(s) vs an expected 0.951 site(s)) and p_value 2.413e-6

Factor    	log2 enrichment	p_value 	number of hits	expected number of hits in null distribution
MIN_EPOD  	2.657e0       	2.413e-6	6              	0.951
RDM_EPOD  	2.308e0       	8.989e-5	5              	1.009


Motif 72 (AGAGAATCGTAACG) with peak height 4.451765407549692 is most likely MIN_EPOD, with log2 fold enrichment 2.920e0 (9 site(s) vs an expected 1.189 site(s)) and p_value 5.667e-10

Factor    	log2 enrichment	p_value 	number of hits	expected number of hits in null distribution
MIN_EPOD  	2.920e0       	5.667e-10	9              	1.189
RDM_EPOD  	2.665e0       	7.187e-8	8              	1.262


Motif 73 (ATTGCTTATTGACATA) with peak height 4.2344646946076265 is most likely RDM_EPOD, with log2 fold enrichment 2.624e0 (7 site(s) vs an expected 1.136 site(s)) and p_value 5.134e-7

Factor    	log2 enrichment	p_value 	number of hits	expected number of hits in null distribution
RDM_EPOD  	2.624e0       	5.134e-7	7              	1.136
MIN_EPOD  	2.487e0       	9.734e-6	6              	1.070


Motif 74 (CAAAATGCACCTGTA) with peak height 5.440904269363602 is most likely MIN_EPOD, with log2 fold enrichment 2.879e0 (7 site(s) vs an expected 0.951 site(s)) and p_value 4.004e-8

Factor    	log2 enrichment	p_value 	number of hits	expected number of hits in null distribution
MIN_EPOD  	2.879e0       	4.004e-8	7              	0.951
RDM_EPOD  	2.794e0       	6.424e-8	7              	1.009
CpxR      	8.315e0       	4.308e-6	1              	0.003


Motif 75 (CCTTAAGACATG) with peak height 5.255745084111004 is most likely MIN_EPOD, with log2 fold enrichment 2.557e0 (7 site(s) vs an expected 1.189 site(s)) and p_value 1.441e-6

Factor    	log2 enrichment	p_value 	number of hits	expected number of hits in null distribution
MIN_EPOD  	2.557e0       	1.441e-6	7              	1.189
RDM_EPOD  	2.472e0       	2.279e-6	7              	1.262
FNR       	7.946e0       	7.382e-6	1              	0.004


Motif 76 (ACAAAGTGCGCTTTGTTTA) with peak height 5.55529622741491 has no hits!


Motif 77 (ATTACAGACAAGCCT) with peak height 3.1808519547651626 is most likely RDM_EPOD, with log2 fold enrichment 2.665e0 (8 site(s) vs an expected 1.262 site(s)) and p_value 7.188e-8

Factor    	log2 enrichment	p_value 	number of hits	expected number of hits in null distribution
RDM_EPOD  	2.665e0       	7.188e-8	8              	1.262
RcsAB     	1.073e1       	1.568e-7	1              	0.001
MIN_EPOD  	2.557e0       	1.442e-6	7              	1.189


Motif 78 (TATTGCTCGTGTA) with peak height 3.749305889256381 is most likely MIN_EPOD, with log2 fold enrichment 2.223e0 (10 site(s) vs an expected 2.141 site(s)) and p_value 9.598e-7

Factor    	log2 enrichment	p_value 	number of hits	expected number of hits in null distribution
MIN_EPOD  	2.223e0       	9.598e-7	10             	2.141
NagC      	8.451e0       	3.851e-6	1              	0.003
PhoB      	8.235e0       	5.193e-6	1              	0.003
OmpR      	7.962e0       	7.574e-6	1              	0.004
RDM_EPOD  	1.986e0       	1.698e-5	9              	2.272
CRP       	5.282e0       	3.071e-4	1              	0.026
Nac       	5.210e0       	3.394e-4	1              	0.027


Motif 79 (ATTAATCTCGATTTG) with peak height 4.55457527483134 is most likely ZraR, with log2 fold enrichment 1.024e1 (1 site(s) vs an expected 0.001 site(s)) and p_value 3.006e-7

Factor    	log2 enrichment	p_value 	number of hits	expected number of hits in null distribution
ZraR      	1.024e1       	3.006e-7	1              	0.001
Lrp       	8.071e0       	6.035e-6	1              	0.004
Nac       	6.403e0       	6.071e-5	1              	0.012
MIN_EPOD  	2.394e0       	6.394e-5	5              	0.951
RDM_EPOD  	2.308e0       	8.989e-5	5              	1.009


Motif 80 (TAGTAAAAAAACAAGT) with peak height 7.043769373373997 is most likely MIN_EPOD, with log2 fold enrichment 3.072e0 (5 site(s) vs an expected 0.595 site(s)) and p_value 0.000e0

Factor    	log2 enrichment	p_value 	number of hits	expected number of hits in null distribution
MIN_EPOD  	3.072e0       	0.000e0 	5              	0.595
RDM_EPOD  	2.987e0       	0.000e0 	5              	0.631


Motif 81 (ATGTGTTATTTTTTT) with peak height 3.817746581922925 is most likely MIN_EPOD, with log2 fold enrichment 2.831e0 (11 site(s) vs an expected 1.546 site(s)) and p_value 9.288e-11

Factor    	log2 enrichment	p_value 	number of hits	expected number of hits in null distribution
MIN_EPOD  	2.831e0       	9.288e-11	11             	1.546
RDM_EPOD  	2.745e0       	1.873e-10	11             	1.640
IHF       	7.788e0       	9.419e-6	1              	0.005


Motif 82 (CTATCTGATGGTAA) with peak height 6.386667767123331 is most likely OxyR, with log2 fold enrichment 1.013e1 (1 site(s) vs an expected 0.001 site(s)) and p_value 3.173e-7

Factor    	log2 enrichment	p_value 	number of hits	expected number of hits in null distribution
OxyR      	1.013e1       	3.173e-7	1              	0.001
Nac       	7.092e0       	2.143e-5	1              	0.007
MIN_EPOD  	2.335e0       	9.050e-4	3              	0.595
RDM_EPOD  	2.250e0       	1.139e-3	3              	0.631


Motif 83 (TTATCTTTATGCTAA) with peak height 6.367392710319348 is most likely MIN_EPOD, with log2 fold enrichment 3.072e0 (5 site(s) vs an expected 0.595 site(s)) and p_value 0.000e0

Factor    	log2 enrichment	p_value 	number of hits	expected number of hits in null distribution
MIN_EPOD  	3.072e0       	0.000e0 	5              	0.595
RDM_EPOD  	2.665e0       	3.197e-5	4              	0.631


Motif 84 (TCAGTCCTGCAAAATT) with peak height 4.55575019573233 is most likely MIN_EPOD, with log2 fold enrichment 2.487e0 (4 site(s) vs an expected 0.714 site(s)) and p_value 1.286e-4

Factor    	log2 enrichment	p_value 	number of hits	expected number of hits in null distribution
MIN_EPOD  	2.487e0       	1.286e-4	4              	0.714
RDM_EPOD  	2.402e0       	1.717e-4	4              	0.757


Motif 85 (GGCAAGATTATA) with peak height 4.277923309606436 is most likely ArcA, with log2 fold enrichment 7.356e0 (1 site(s) vs an expected 0.006 site(s)) and p_value 1.672e-5

Factor    	log2 enrichment	p_value 	number of hits	expected number of hits in null distribution
ArcA      	7.356e0       	1.672e-5	1              	0.006
MIN_EPOD  	2.335e0       	2.907e-5	6              	1.189
RDM_EPOD  	2.250e0       	4.304e-5	6              	1.262


Motif 86 (ATAAATTGCAGTAAACA) with peak height 6.332053704318874 is most likely RDM_EPOD, with log2 fold enrichment 2.250e0 (3 site(s) vs an expected 0.631 site(s)) and p_value 1.139e-3

Factor    	log2 enrichment	p_value 	number of hits	expected number of hits in null distribution
RDM_EPOD  	2.250e0       	1.139e-3	3              	0.631
MIN_EPOD  	1.750e0       	1.396e-2	2              	0.595


Motif 87 (AGCGGTGTAGGCCTGATAAG) with peak height 3.9162503882081943 is most likely MIN_EPOD, with log2 fold enrichment 2.394e0 (5 site(s) vs an expected 0.952 site(s)) and p_value 6.397e-5

Factor    	log2 enrichment	p_value 	number of hits	expected number of hits in null distribution
MIN_EPOD  	2.394e0       	6.397e-5	5              	0.952
RDM_EPOD  	2.308e0       	8.992e-5	5              	1.009


Motif 88 (ACATTGAAGGAAACAT) with peak height 4.285760185876079 is most likely MIN_EPOD, with log2 fold enrichment 3.072e0 (9 site(s) vs an expected 1.070 site(s)) and p_value 0.000e0

Factor    	log2 enrichment	p_value 	number of hits	expected number of hits in null distribution
MIN_EPOD  	3.072e0       	0.000e0 	9              	1.070
RDM_EPOD  	2.817e0       	8.108e-9	8              	1.136


Motif 89 (AATAATTCTCTGGACT) with peak height 5.505674175487495 is most likely MIN_EPOD, with log2 fold enrichment 2.750e0 (4 site(s) vs an expected 0.595 site(s)) and p_value 2.379e-5

Factor    	log2 enrichment	p_value 	number of hits	expected number of hits in null distribution
MIN_EPOD  	2.750e0       	2.379e-5	4              	0.595
RDM_EPOD  	2.665e0       	3.197e-5	4              	0.631


Motif 90 (TTTTCAATAATACCCA) with peak height 4.620848610622331 is most likely MIN_EPOD, with log2 fold enrichment 3.072e0 (6 site(s) vs an expected 0.714 site(s)) and p_value 0.000e0

Factor    	log2 enrichment	p_value 	number of hits	expected number of hits in null distribution
MIN_EPOD  	3.072e0       	0.000e0 	6              	0.714
RDM_EPOD  	2.987e0       	0.000e0 	6              	0.757


Motif 91 (CTATTGGACTTATT) with peak height 3.7760940171225923 is most likely MIN_EPOD, with log2 fold enrichment 2.657e0 (6 site(s) vs an expected 0.951 site(s)) and p_value 2.413e-6

Factor    	log2 enrichment	p_value 	number of hits	expected number of hits in null distribution
MIN_EPOD  	2.657e0       	2.413e-6	6              	0.951
GlaR      	8.687e0       	2.573e-6	1              	0.002
RDM_EPOD  	2.571e0       	3.623e-6	6              	1.009
Fis       	7.393e0       	1.543e-5	1              	0.006


Motif 92 (TATTCCATGAGCA) with peak height 3.2806044783562167 is most likely MIN_EPOD, with log2 fold enrichment 2.557e0 (7 site(s) vs an expected 1.189 site(s)) and p_value 1.442e-6

Factor    	log2 enrichment	p_value 	number of hits	expected number of hits in null distribution
MIN_EPOD  	2.557e0       	1.442e-6	7              	1.189
RDM_EPOD  	2.472e0       	2.279e-6	7              	1.262


Motif 93 (AATTAATGTTATTTTT) with peak height 4.39288113222096 is most likely MIN_EPOD, with log2 fold enrichment 2.586e0 (5 site(s) vs an expected 0.833 site(s)) and p_value 1.779e-5

Factor    	log2 enrichment	p_value 	number of hits	expected number of hits in null distribution
MIN_EPOD  	2.586e0       	1.779e-5	5              	0.833
RDM_EPOD  	2.501e0       	2.519e-5	5              	0.883


Motif 94 (TATACATCTCACTG) with peak height 5.018382673973644 is most likely MIN_EPOD, with log2 fold enrichment 3.072e0 (6 site(s) vs an expected 0.714 site(s)) and p_value 0.000e0

Factor    	log2 enrichment	p_value 	number of hits	expected number of hits in null distribution
MIN_EPOD  	3.072e0       	0.000e0 	6              	0.714
RDM_EPOD  	2.987e0       	0.000e0 	6              	0.757
Nac       	6.826e0       	3.225e-5	1              	0.009


Motif 95 (ATTGATTCTGCACAT) with peak height 5.080101306445308 is most likely MIN_EPOD, with log2 fold enrichment 1.750e0 (2 site(s) vs an expected 0.595 site(s)) and p_value 1.396e-2

Factor    	log2 enrichment	p_value 	number of hits	expected number of hits in null distribution
MIN_EPOD  	1.750e0       	1.396e-2	2              	0.595


Motif 96 (TTTGTGCCGTTATTTCT) with peak height 6.680916632484972 is most likely MIN_EPOD, with log2 fold enrichment 3.072e0 (4 site(s) vs an expected 0.476 site(s)) and p_value 0.000e0

Factor    	log2 enrichment	p_value 	number of hits	expected number of hits in null distribution
MIN_EPOD  	3.072e0       	0.000e0 	4              	0.476
RDM_EPOD  	2.987e0       	0.000e0 	4              	0.505


Motif 97 (AGATATAGAGGATTTTGGG) with peak height 4.964564758744229 is most likely MIN_EPOD, with log2 fold enrichment 3.072e0 (5 site(s) vs an expected 0.595 site(s)) and p_value 0.000e0

Factor    	log2 enrichment	p_value 	number of hits	expected number of hits in null distribution
MIN_EPOD  	3.072e0       	0.000e0 	5              	0.595
RDM_EPOD  	2.987e0       	0.000e0 	5              	0.631


Motif 98 (GTGCACCATATTTAT) with peak height 3.4123520521644917 is most likely MIN_EPOD, with log2 fold enrichment 2.657e0 (6 site(s) vs an expected 0.951 site(s)) and p_value 2.413e-6

Factor    	log2 enrichment	p_value 	number of hits	expected number of hits in null distribution
MIN_EPOD  	2.657e0       	2.413e-6	6              	0.951
RDM_EPOD  	2.571e0       	3.623e-6	6              	1.009


Motif 99 (TCAAAATAAGAAGAGG) with peak height 5.803919314389294 is most likely MIN_EPOD, with log2 fold enrichment 2.750e0 (4 site(s) vs an expected 0.595 site(s)) and p_value 2.379e-5

Factor    	log2 enrichment	p_value 	number of hits	expected number of hits in null distribution
MIN_EPOD  	2.750e0       	2.379e-5	4              	0.595
RDM_EPOD  	2.665e0       	3.197e-5	4              	0.631


Motif 100 (ATTTACTGTCATA) with peak height 4.893821616291213 is most likely ArcA, with log2 fold enrichment 8.114e0 (1 site(s) vs an expected 0.004 site(s)) and p_value 5.418e-6

Factor    	log2 enrichment	p_value 	number of hits	expected number of hits in null distribution
ArcA      	8.114e0       	5.418e-6	1              	0.004
RDM_EPOD  	2.402e0       	1.716e-4	4              	0.757
MIN_EPOD  	2.072e0       	2.458e-3	3              	0.714


Motif 101 (TACGGTTCTGTTCACT) with peak height 4.936804235265678 is most likely MIN_EPOD, with log2 fold enrichment 1.072e0 (1 site(s) vs an expected 0.476 site(s)) and p_value 7.200e-2

Factor    	log2 enrichment	p_value 	number of hits	expected number of hits in null distribution
MIN_EPOD  	1.072e0       	7.200e-2	1              	0.476


Motif 102 (AAATAAAAACAAAAACC) with peak height 5.739853203745972 is most likely MIN_EPOD, with log2 fold enrichment 2.657e0 (3 site(s) vs an expected 0.476 site(s)) and p_value 2.000e-4

Factor    	log2 enrichment	p_value 	number of hits	expected number of hits in null distribution
MIN_EPOD  	2.657e0       	2.000e-4	3              	0.476
RDM_EPOD  	2.572e0       	2.534e-4	3              	0.505


Motif 103 (TGAGTTGAAATTTC) with peak height 4.142656298823346 is most likely PhoP, with log2 fold enrichment 1.016e1 (1 site(s) vs an expected 0.001 site(s)) and p_value 3.190e-7

Factor    	log2 enrichment	p_value 	number of hits	expected number of hits in null distribution
PhoP      	1.016e1       	3.190e-7	1              	0.001
MIN_EPOD  	2.809e0       	2.830e-6	5              	0.714
RDM_EPOD  	2.724e0       	4.033e-6	5              	0.757


Motif 104 (CACGTATATCAGGCG) with peak height 5.464361492246326 is most likely MIN_EPOD, with log2 fold enrichment 1.750e0 (2 site(s) vs an expected 0.595 site(s)) and p_value 1.396e-2

Factor    	log2 enrichment	p_value 	number of hits	expected number of hits in null distribution
MIN_EPOD  	1.750e0       	1.396e-2	2              	0.595
RDM_EPOD  	1.665e0       	1.647e-2	2              	0.631


Motif 105 (AGTCAAGTCGGGT) with peak height 5.717151143611043 is most likely MIN_EPOD, with log2 fold enrichment 3.072e0 (4 site(s) vs an expected 0.476 site(s)) and p_value 0.000e0

Factor    	log2 enrichment	p_value 	number of hits	expected number of hits in null distribution
MIN_EPOD  	3.072e0       	0.000e0 	4              	0.476
RDM_EPOD  	2.987e0       	0.000e0 	4              	0.505


Motif 106 (TTATTGGAGGATA) with peak height 3.768015867333595 is most likely MIN_EPOD, with log2 fold enrichment 2.809e0 (5 site(s) vs an expected 0.714 site(s)) and p_value 2.829e-6

Factor    	log2 enrichment	p_value 	number of hits	expected number of hits in null distribution
MIN_EPOD  	2.809e0       	2.829e-6	5              	0.714
RDM_EPOD  	2.724e0       	4.033e-6	5              	0.757


Motif 107 (CCTCGCAACTCTTT) with peak height 4.618167530456412 is most likely MIN_EPOD, with log2 fold enrichment 1.487e0 (1 site(s) vs an expected 0.357 site(s)) and p_value 3.906e-2

Factor    	log2 enrichment	p_value 	number of hits	expected number of hits in null distribution
MIN_EPOD  	1.487e0       	3.906e-2	1              	0.357
RDM_EPOD  	1.402e0       	4.373e-2	1              	0.378


Motif 108 (TGATTTTTTTCATGATGT) with peak height 4.980430590217372 is most likely MIN_EPOD, with log2 fold enrichment 2.487e0 (2 site(s) vs an expected 0.357 site(s)) and p_value 1.682e-3

Factor    	log2 enrichment	p_value 	number of hits	expected number of hits in null distribution
MIN_EPOD  	2.487e0       	1.682e-3	2              	0.357
RDM_EPOD  	1.402e0       	4.373e-2	1              	0.378


Motif 109 (ATGGGTCAAAGAATA) with peak height 4.623840055004482 is most likely MIN_EPOD, with log2 fold enrichment 2.657e0 (3 site(s) vs an expected 0.476 site(s)) and p_value 2.000e-4

Factor    	log2 enrichment	p_value 	number of hits	expected number of hits in null distribution
MIN_EPOD  	2.657e0       	2.000e-4	3              	0.476
RDM_EPOD  	2.572e0       	2.533e-4	3              	0.505


Motif 110 (TACTTGGTTGGTAAT) with peak height 5.1083727965005 is most likely MIN_EPOD, with log2 fold enrichment 2.487e0 (2 site(s) vs an expected 0.357 site(s)) and p_value 1.682e-3

Factor    	log2 enrichment	p_value 	number of hits	expected number of hits in null distribution
MIN_EPOD  	2.487e0       	1.682e-3	2              	0.357
RDM_EPOD  	2.402e0       	2.008e-3	2              	0.378


Motif 111 (GGGTTTGATAACAATT) with peak height 4.267701778069021 is most likely MIN_EPOD, with log2 fold enrichment 3.072e0 (3 site(s) vs an expected 0.357 site(s)) and p_value 0.000e0

Factor    	log2 enrichment	p_value 	number of hits	expected number of hits in null distribution
MIN_EPOD  	3.072e0       	0.000e0 	3              	0.357
RDM_EPOD  	2.987e0       	0.000e0 	3              	0.378


Motif 112 (GGGGTCGAATAGTGA) with peak height 4.510017503103599 is most likely MIN_EPOD, with log2 fold enrichment 3.072e0 (3 site(s) vs an expected 0.357 site(s)) and p_value 0.000e0

Factor    	log2 enrichment	p_value 	number of hits	expected number of hits in null distribution
MIN_EPOD  	3.072e0       	0.000e0 	3              	0.357
RDM_EPOD  	2.402e0       	2.008e-3	2              	0.378


Motif 113 (TGACATTGATTTGTCT) with peak height 4.9543269358087585 is most likely Nac, with log2 fold enrichment 7.834e0 (1 site(s) vs an expected 0.004 site(s)) and p_value 6.400e-6

Factor    	log2 enrichment	p_value 	number of hits	expected number of hits in null distribution
Nac       	7.834e0       	6.400e-6	1              	0.004
MIN_EPOD  	2.487e0       	1.682e-3	2              	0.357
RDM_EPOD  	2.402e0       	2.008e-3	2              	0.378


Motif 114 (GCAACAGCGTGGTGCGCAGA) with peak height 4.466879715467592 is most likely MIN_EPOD, with log2 fold enrichment 3.072e0 (3 site(s) vs an expected 0.357 site(s)) and p_value 0.000e0

Factor    	log2 enrichment	p_value 	number of hits	expected number of hits in null distribution
MIN_EPOD  	3.072e0       	0.000e0 	3              	0.357
RDM_EPOD  	2.987e0       	0.000e0 	3              	0.378


Motif 115 (AAAATAAGTGTAAC) with peak height 3.4099664024905563 is most likely McbR, with log2 fold enrichment 1.155e1 (1 site(s) vs an expected 0.000 site(s)) and p_value 4.890e-8

Factor    	log2 enrichment	p_value 	number of hits	expected number of hits in null distribution
McbR      	1.155e1       	4.890e-8	1              	0.000
RDM_EPOD  	2.794e0       	6.423e-8	7              	1.009
H-NS      	9.195e0       	1.273e-6	1              	0.002
MIN_EPOD  	2.657e0       	2.413e-6	6              	0.951


Motif 116 (GTGGTTCCATAACAAC) with peak height 5.078013338234484 is most likely MIN_EPOD, with log2 fold enrichment 3.072e0 (4 site(s) vs an expected 0.476 site(s)) and p_value 0.000e0

Factor    	log2 enrichment	p_value 	number of hits	expected number of hits in null distribution
MIN_EPOD  	3.072e0       	0.000e0 	4              	0.476
RDM_EPOD  	2.987e0       	0.000e0 	4              	0.505


Motif 117 (ATTTTGTTGTTTATTTTT) with peak height 5.26681497055797 is most likely MIN_EPOD, with log2 fold enrichment 2.487e0 (2 site(s) vs an expected 0.357 site(s)) and p_value 1.682e-3

Factor    	log2 enrichment	p_value 	number of hits	expected number of hits in null distribution
MIN_EPOD  	2.487e0       	1.682e-3	2              	0.357
RDM_EPOD  	2.402e0       	2.008e-3	2              	0.378


Motif 118 (AAGTTGGAAAAATTTTC) with peak height 3.865549293854575 is most likely AllR, with log2 fold enrichment 1.444e1 (2 site(s) vs an expected 0.000 site(s)) and p_value 2.685e-14

Factor    	log2 enrichment	p_value 	number of hits	expected number of hits in null distribution
AllR      	1.444e1       	2.685e-14	2              	0.000
MIN_EPOD  	1.487e0       	3.906e-2	1              	0.357
RDM_EPOD  	1.402e0       	4.373e-2	1              	0.378


Motif 119 (ACCCGCATCAGCTGCAGCA) with peak height 3.541262658055977 is most likely RDM_EPOD, with log2 fold enrichment 2.987e0 (5 site(s) vs an expected 0.631 site(s)) and p_value 0.000e0

Factor    	log2 enrichment	p_value 	number of hits	expected number of hits in null distribution
RDM_EPOD  	2.987e0       	0.000e0 	5              	0.631
MIN_EPOD  	2.750e0       	2.379e-5	4              	0.595


Motif 120 (ACATATAGTTACGAC) with peak height 5.2623266155473525 is most likely MIN_EPOD, with log2 fold enrichment 2.072e0 (1 site(s) vs an expected 0.238 site(s)) and p_value 1.414e-2

Factor    	log2 enrichment	p_value 	number of hits	expected number of hits in null distribution
MIN_EPOD  	2.072e0       	1.414e-2	1              	0.238
RDM_EPOD  	1.987e0       	1.592e-2	1              	0.252


Motif 121 (ATGTCTTAGAGAG) with peak height 3.3175521927195475 is most likely MIN_EPOD, with log2 fold enrichment 2.072e0 (2 site(s) vs an expected 0.476 site(s)) and p_value 6.127e-3

Factor    	log2 enrichment	p_value 	number of hits	expected number of hits in null distribution
MIN_EPOD  	2.072e0       	6.127e-3	2              	0.476
RDM_EPOD  	1.987e0       	7.272e-3	2              	0.505


Motif 122 (GCTCAAAAAGCTAAAG) with peak height 3.9886857415852104 is most likely MIN_EPOD, with log2 fold enrichment 3.072e0 (2 site(s) vs an expected 0.238 site(s)) and p_value 0.000e0

Factor    	log2 enrichment	p_value 	number of hits	expected number of hits in null distribution
MIN_EPOD  	3.072e0       	0.000e0 	2              	0.238
RDM_EPOD  	2.987e0       	0.000e0 	2              	0.252


Motif 123 (GTTGCTGCGACTGACGGCCC) with peak height 4.982142339464048 is most likely RDM_EPOD, with log2 fold enrichment 2.987e0 (2 site(s) vs an expected 0.252 site(s)) and p_value 0.000e0

Factor    	log2 enrichment	p_value 	number of hits	expected number of hits in null distribution
RDM_EPOD  	2.987e0       	0.000e0 	2              	0.252


Motif 124 (GCGGCGTTCTGCCTGTG) with peak height 4.433004830952716 has no hits!


Motif 125 (CAATTCCTTGAAAAGGA) with peak height 3.9708534120821364 is most likely MIN_EPOD, with log2 fold enrichment 3.072e0 (2 site(s) vs an expected 0.238 site(s)) and p_value 0.000e0

Factor    	log2 enrichment	p_value 	number of hits	expected number of hits in null distribution
MIN_EPOD  	3.072e0       	0.000e0 	2              	0.238
RDM_EPOD  	2.987e0       	0.000e0 	2              	0.252


Motif 126 (GTAATAACCTTTATCCA) with peak height 4.300368189753921 is most likely RDM_EPOD, with log2 fold enrichment 2.987e0 (3 site(s) vs an expected 0.378 site(s)) and p_value 0.000e0

Factor    	log2 enrichment	p_value 	number of hits	expected number of hits in null distribution
RDM_EPOD  	2.987e0       	0.000e0 	3              	0.378
MIN_EPOD  	2.487e0       	1.682e-3	2              	0.357


Motif 127 (TATGACAAGAGTCGCCGCA) with peak height 3.602025372435953 is most likely MIN_EPOD, with log2 fold enrichment 3.072e0 (4 site(s) vs an expected 0.476 site(s)) and p_value 0.000e0

Factor    	log2 enrichment	p_value 	number of hits	expected number of hits in null distribution
MIN_EPOD  	3.072e0       	0.000e0 	4              	0.476
RDM_EPOD  	2.987e0       	0.000e0 	4              	0.505


Motif 128 (TGTTTTTTCTCTGTTT) with peak height 3.1165330373270703 is most likely MIN_EPOD, with log2 fold enrichment 1.849e0 (3 site(s) vs an expected 0.833 site(s)) and p_value 5.196e-3

Factor    	log2 enrichment	p_value 	number of hits	expected number of hits in null distribution
MIN_EPOD  	1.849e0       	5.196e-3	3              	0.833
RDM_EPOD  	1.764e0       	6.456e-3	3              	0.883


Motif 129 (GAATCTGCGACCGAAA) with peak height 2.9401595756965127 is most likely MIN_EPOD, with log2 fold enrichment 3.072e0 (2 site(s) vs an expected 0.238 site(s)) and p_value 0.000e0

Factor    	log2 enrichment	p_value 	number of hits	expected number of hits in null distribution
MIN_EPOD  	3.072e0       	0.000e0 	2              	0.238
RDM_EPOD  	2.987e0       	0.000e0 	2              	0.252


Motif 130 (GAAACCAGGGGCGGCTAAA) with peak height 3.078894749978733 is most likely MIN_EPOD, with log2 fold enrichment 3.072e0 (4 site(s) vs an expected 0.476 site(s)) and p_value 0.000e0

Factor    	log2 enrichment	p_value 	number of hits	expected number of hits in null distribution
MIN_EPOD  	3.072e0       	0.000e0 	4              	0.476
RDM_EPOD  	2.987e0       	0.000e0 	4              	0.505


Motif 131 (GTAAGTGAGTGCTC) with peak height 3.612164358877541 is most likely MIN_EPOD, with log2 fold enrichment 2.072e0 (1 site(s) vs an expected 0.238 site(s)) and p_value 1.414e-2

Factor    	log2 enrichment	p_value 	number of hits	expected number of hits in null distribution
MIN_EPOD  	2.072e0       	1.414e-2	1              	0.238
RDM_EPOD  	1.987e0       	1.592e-2	1              	0.252


Motif 132 (ACGTGGAGAATAAGCTGCA) with peak height 3.455624344676073 is most likely MIN_EPOD, with log2 fold enrichment 3.072e0 (4 site(s) vs an expected 0.476 site(s)) and p_value 0.000e0

Factor    	log2 enrichment	p_value 	number of hits	expected number of hits in null distribution
MIN_EPOD  	3.072e0       	0.000e0 	4              	0.476
RDM_EPOD  	2.987e0       	0.000e0 	4              	0.505


Motif 133 (GAGCCCTTAACAT) with peak height 3.148486244311354 is most likely MIN_EPOD, with log2 fold enrichment 3.072e0 (6 site(s) vs an expected 0.714 site(s)) and p_value 0.000e0

Factor    	log2 enrichment	p_value 	number of hits	expected number of hits in null distribution
MIN_EPOD  	3.072e0       	0.000e0 	6              	0.714
RDM_EPOD  	2.987e0       	0.000e0 	6              	0.757
NarL      	1.013e1       	3.323e-7	1              	0.001
FNR       	8.716e0       	2.355e-6	1              	0.002


Motif 134 (GTAGAGATATCGGAGCATT) with peak height 4.328922313240391 is most likely MIN_EPOD, with log2 fold enrichment 3.072e0 (2 site(s) vs an expected 0.238 site(s)) and p_value 0.000e0

Factor    	log2 enrichment	p_value 	number of hits	expected number of hits in null distribution
MIN_EPOD  	3.072e0       	0.000e0 	2              	0.238
RDM_EPOD  	2.987e0       	0.000e0 	2              	0.252
